# Drying parameters and aging modulate protective properties of vitrified trehalose

**DOI:** 10.64898/2026.01.16.700019

**Authors:** UGVSS Kumara, Thomas C Boothby

## Abstract

Storage of biological materials is essential for medical, research, and biotechnological applications. While cold-chain preservation is effective, it is costly, infrastructure-dependent, and vulnerable to disruption. Room-temperature dry storage, inspired by desiccation-tolerant organisms, provides an alternative by stabilizing biomolecules in vitrified (“glass-like”) matrices that limit molecular motion which if not reduced can lead to breakdown and loss of integrity. Trehalose is widely used as a vitrifying agent, but its protective capacity depends on glassy properties shaped by drying methods, environmental conditions, storage duration, and the type of preserved molecule. Systematic studies linking these factors to short and long-term stability remain limited. Here, we examine how drying conditions and storage duration influence the stability of DNA, RNA, and enzymes in vitrified trehalose systems. DNA remained stable under all conditions, independent of trehalose or drying parameters, reflecting its intrinsic resistance to desiccation-induced damage. RNA showed moderate sensitivity to drying without trehalose but was stabilized in its presence, although RNA integrity did not consistently correlate with measured vitrified properties. In contrast, enzymes were highly sensitive to drying without trehalose and were strongly protected under conditions that promoted favorable vitrified properties. Enzyme protection after 30 min correlated with high glass transition temperature. However, during prolonged drying, increased glass transition temperature was inversely correlated with enzyme protection and was a better indicator of detrimental physical aging of the vitrified system. These findings present the first insight into how drying methods, environmental conditions, and storage duration shape vitrified properties and stability. They guide optimization of room-temperature preservation.

**Graphical abstract:** 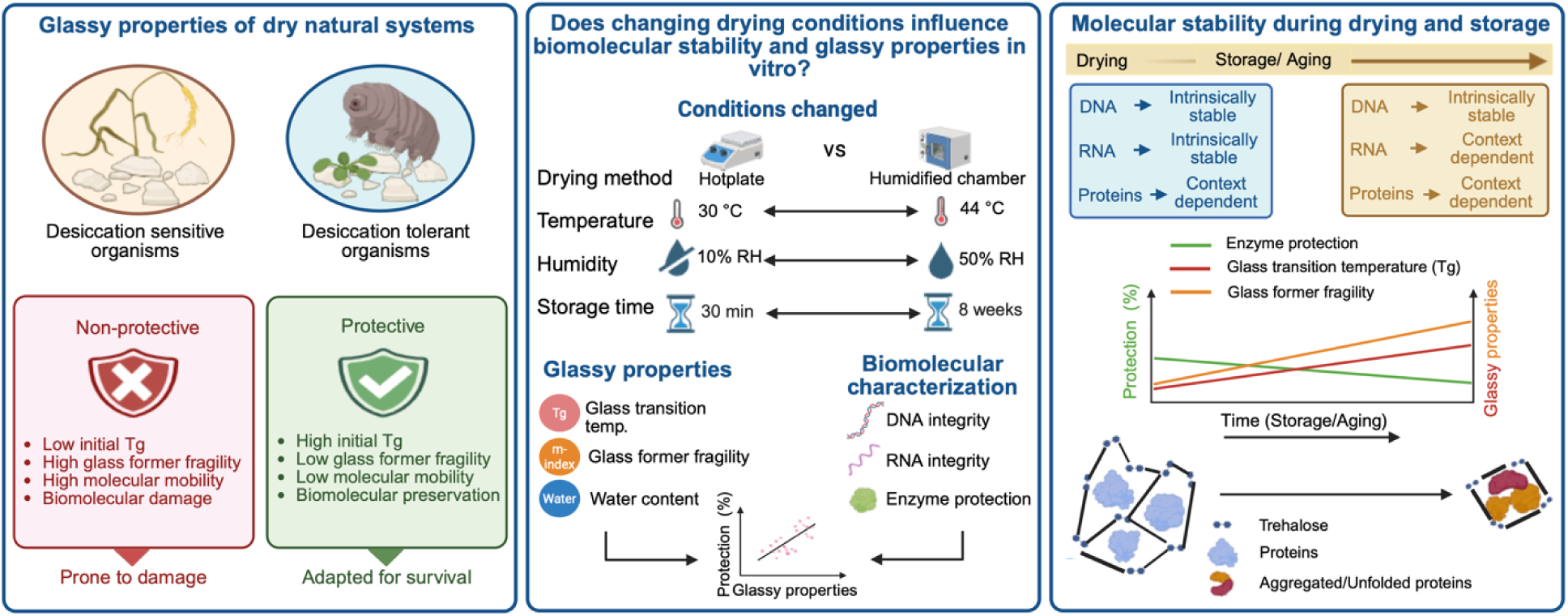

**Highlights:**

- Nucleic acids are generally stable under various drying conditions/vitrified systems
- Protein preservation benefits from specific mediators and vitrified properties
- Drying method and parameters tune vitrified properties and protection
- Vitrified systems age over time, changing properties and protectiveness
- Initially high Tg is protective, increasing Tg and fragility over time indicate aging

## Introduction

Cryopreservation and other cold-chain storage methods, including freezing and refrigeration, are widely used for preserving biological materials in medicine, biotechnology, and research[1–7]. Cold-chain storage effectively slows or even halts molecular motion and biochemical degradation[1–7]. However, it requires specialized equipment as well as stable electrical and maintenance infrastructure, making it costly and susceptible to failures such as power outages or equipment malfunctions[8–13]. Both the cost and logistical considerations of the cold-chain limits its use in remote and developing parts of the world, as well as under austere conditions such as on the battlefield or during natural disasters. These limitations have driven interest in alternative storage strategies that enable stable long-term preservation of biological material at room temperature[14].

One promising approach draws inspiration from biological systems that have evolved to survive extreme desiccation or drying[15–23]. To survive extreme drying, organisms undergo anhydrobiosis, a term from Greek meaning “life without water.” Anhydrobiotic organisms such as tardigrades, nematodes, and certain seeds, employ vitrification to stabilize cellular components in a dry glass-like state[23–29]. Similar to the cold-chain, dry vitrification can prevent biological degradation by dramatically reducing molecular motion, diffusion and chemical reaction rates, thereby preserving biomolecules in a physically arrested state[23,27,30–33]. These vitrified, or glassy, states are characterized by key material properties such as glass transition temperature (Tg), glass former fragility (m-index), and water content, which collectively influence the stability and protective capacity of vitrified systems[23,28,29,34–43].

Numerous drying methods have been used to induce vitrification in biological materials[44–49] with varying success in replicating the natural dehydration processes observed in desiccation-tolerant organisms. However, the final quality of dried samples, including structural integrity, functional preservation, and long-term stability, is shaped by several factors, such as drying rate, environmental conditions, and the presence or absence of protective agents like disaccharides such as trehalose[44–48,50–55].

Several sugars, including trehalose, are known to be able to form stable glassy states that protect biological molecules[28,34,45]. The protective capacity of these glasses for diverse clients in part depends on their material properties, which in turn are influenced by drying methods, environmental parameters, and time in the vitrified state[45,48,56–58]. However, systematic studies that allow for direct comparison of how drying parameters influence resulting material properties of vitrified systems, and how these in turn modulate protective capacity for diverse clients are lacking.

This study systematically compares two desiccation techniques: humidified chamber drying and rapid hotplate drying. We examine how variations in temperature and humidity affect the glassy properties and protective capacity of vitrified trehalose, both immediately after drying and during storage. To evaluate protective capacity for diverse clients we examine these two drying methods, environmental parameters, and the influence of time in the vitrified state on DNA, RNA, and two different enzymes.

Our results indicate that the drying method and environmental conditions directly shape the physical state of the glassy matrix and its evolution during prolonged storage, which in turn controls the stability of embedded biomolecules. DNA maintains its integrity under all conditions regardless of the presence of trehalose, suggesting its structure is inherently resistant to changes in molecular mobility. RNA is more sensitive, with milder drying conditions and the presence of trehalose reducing perturbations RNA integrity. Protein stability was strongly perturbed by drying, but trehalose could ameliorate these perturbations to varying degrees depending on drying method/parameters. Rapid hotplate drying at mild temperatures and low humidity minimized intermediate hydration, limiting enzymatic dysfunction, whereas slower drying at higher temperatures and humidity increased damage. These findings demonstrate that controlling drying rate and environmental conditions can be used to tune the glassy properties of vitrified trehalose and that by modulating these properties protection of diverse biological macromolecules can be optimized outside of the cold-chain.

## Results

### 1. Effects of drying method on the protection of diverse biological client molecules, after immediate drying

#### 1.1. The effects of drying method and environmental parameters on nucleic acid stability with and without trehalose, after immediate drying

DNA integrity was evaluated using hotplate and humidified chamber drying across a range of temperatures and humidity levels, immediately after 30 min of drying (see methods). Considering all variables four experimental setups were created: (1) hotplate with trehalose (**Figure 1A**), (2) humidified chamber with trehalose (**Figure 1B**), (3) hotplate without trehalose **(Supplementary Figure S1A)**, and (4) humidified chamber without trehalose **(Supplementary Figure S1B)**.

**Figure 1.**
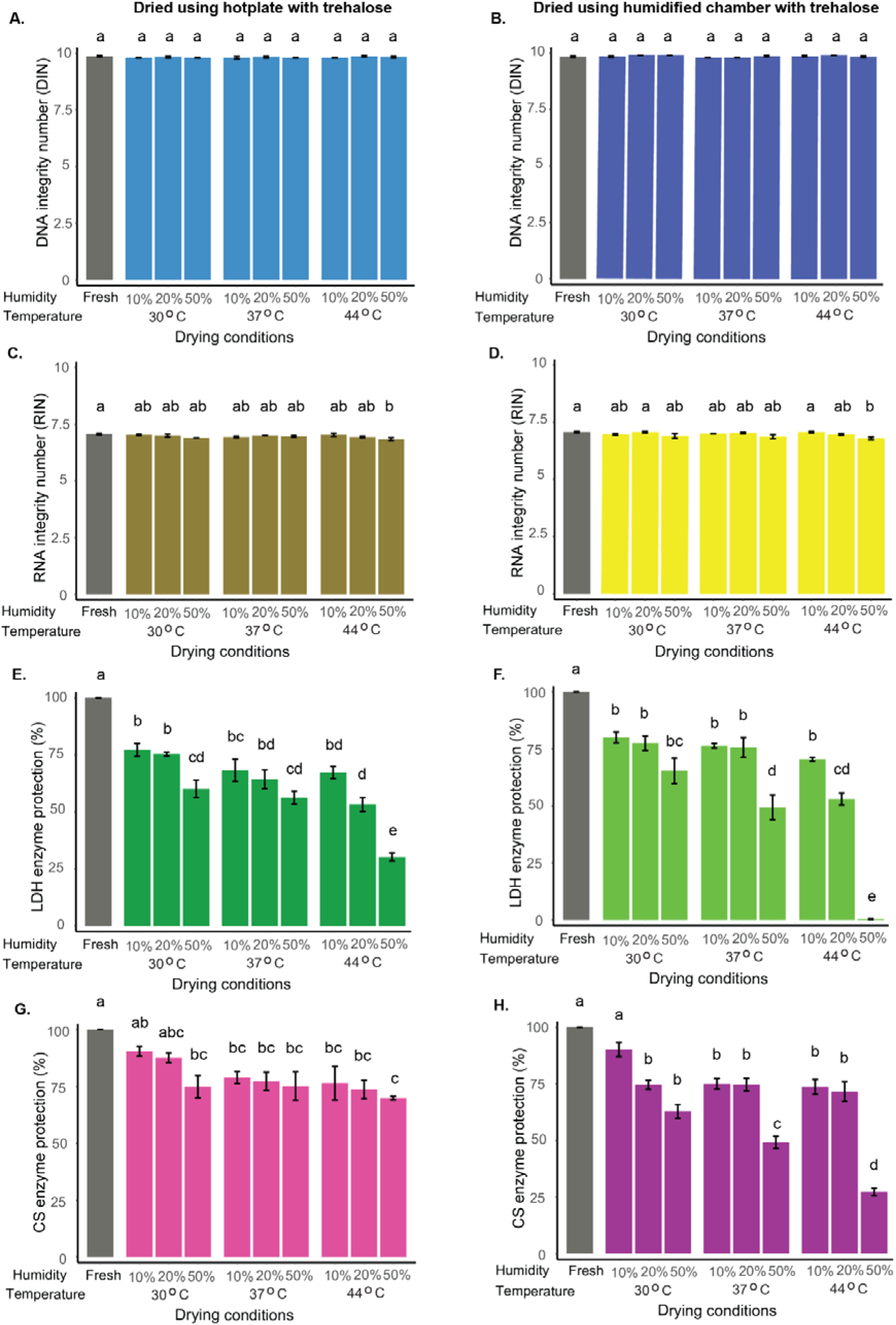
Differential stability of nucleic acids and proteins under various drying conditions in the presence of trehalose. DNA (A,B), RNA (C,D), lactate dehydrogenase (LDH; E,F), and citrate synthase (CS; G,H) samples were dried using two methods-hotplate (A,C,E,G) and humidified chamber (B,D,F,H)-across nine temperature-humidity combinations, all in the presence of 300 mM trehalose. DNA and RNA integrity were assessed using DNA Integrity Number (DIN) and RNA Integrity Number (RIN), respectively, while enzyme activity retention was used to evaluate protein function. Different letters indicate statistically significant differences among drying conditions within each panel, as determined by one-way ANOVA followed by Tukey’s post hoc test (α = 0.05). Data represent mean ± SE from three independent replicates per condition.

Across all drying conditions, DNA integrity remained within the high-quality range, regardless of method or the presence of trehalose, and was comparable to undried controls (**Figure 1A-B; Supplementary Figures S1A-D; S2A-D)**. When trehalose was included **(Supplementary Figure S2A)**, no differences in DNA integrity were observed between drying methods. In contrast, without trehalose **(Supplementary Figure S2B)**, significant differences emerged at 37 °C/50% relative humidity (RH) and 44 °C/10% RH, where the humidified chamber preserved slightly higher integrity. However, even under these conditions, DNA still maintained high overall integrity.

RNA stability was assessed under the same conditions. Control RNA (freshly taken from the freezer) had an initial RNA Integrity Number (RIN) of ∼7 (**Figure 1C,D; Supplementary Figure S1C,D)**. In the presence of trehalose, RNA stability was better maintained during drying conditions on the hotplate, resulting in higher RIN values compared to samples dried without trehalose (**Figure 1C; Supplementary Figures S1C;S3C)**. In contrast, the humidified chamber preserved high RIN values (6.8-7.2) even without trehalose, showing minimal degradation and little added benefit from trehalose was observed (**Figure 1D; Supplementary Figures S1D;S3D).**

When comparing protection between drying methods, RIN values did not differ in the presence of trehalose **(Supplementary Figure S3A)**, whereas without trehalose, the humidified chamber maintained higher RIN values than the hotplate **(Supplementary Figure S3B)**.

To quantify trehalose protection, we calculated fold protection (the ratio of nucleic acid integrity with vs. without trehalose) across drying methods and conditions. Values were near 1 for both DNA and RNA, with slightly higher RNA fold protection under harsher hotplate conditions **(Supplementary Figure S4A,B).** Overall, these findings indicate that RNA is more sensitive than DNA to drying, but both remain largely intact, with trehalose providing extra protection only under severe drying conditions.

#### 1.2. Effects of drying method and environmental parameters on enzyme function with and without trehalose, after immediate drying

To assess how drying methods and environmental conditions affect the protective capacity of vitrified trehalose, we first examined lactate dehydrogenase (LDH) immediately after 30 min of drying.

In the presence of trehalose, LDH activity was substantially preserved, although protection decreased as humidity and temperature increased for both drying methods (**Figure 1E,F**). The highest LDH activity occurred at 30 °C and 10% RH, while the lowest activity was observed at 44 °C and 50% RH, particularly in samples dried in the humidified chamber (**Figure 1F; Supplementary Figure S5A)**. LDH samples dried without trehalose showed minimal activity under all temperatures, humidity levels, and drying methods tested **(Supplementary Figures S1E,F).** When comparing drying methods, LDH activity at 50% RH differed significantly between the hotplate and humidified chamber across all temperatures tested (30 °C, 37 °C, and 44 °C) in the presence of trehalose, whereas no method-dependent differences were observed without trehalose **(Supplementary Figures S5A,B)**.

Citrate synthase (CS) displayed similar trends after 30 min drying, with trehalose providing strong protection that declined at higher humidity and temperature for both drying methods (**Figure 1G,H; Supplementary Figure S6A,B)**. CS activity was highest at 30 °C and 10% RH and lowest under humidified chamber drying at 44 °C and 50% RH (**Figure 1G,H; Supplementary Figure S6A)**. Unlike LDH, CS retained some activity even without trehalose under mild conditions for both drying methods **(Supplementary Figures S1G,H; S6B-D; S7C,D)**, though trehalose consistently improved stability. When comparing drying methods, significant differences in CS activity between the hotplate and humidified chamber were observed at 37 °C and 44 °C in the presence of trehalose, whereas no method-dependent differences occurred without trehalose **(Supplementary Figures S6A,B)**.

Fold-protection analysis further quantified the extent of stabilization across drying methods and conditions, showing that LDH reached its highest protection (∼588-fold) at 37 °C and 20% RH **(Supplementary Figure S4C)**, while CS showed its maximum (∼102-fold) at 44 °C and 50% RH **(Supplementary Figure S4D)**, with both peaks occurring under hotplate drying. Together, these results demonstrate that trehalose is a strong stabilizer for both enzymes, although its effectiveness is shaped by the protein, drying method, temperature, and humidity.

### 2. Effect of drying methods and environmental parameters on water content and protection, after immediate drying

#### 2.1. Effect of drying methods and environmental parameters on water content in dried samples, after immediate drying

After observing that the drying method and environmental parameters during desiccation affect the integrity and function of biomolecules embedded in trehalose glasses, we next examined whether these drying protocols promote vitrified properties that correlate with protection, immediately after 30 min drying. Accurately measuring water content in dried samples is critical for understanding the physical state and stability of vitrified materials as water content can affect both Tg and glass former fragility[28,34,38,59].

Hotplate-dried samples showed consistently low water content with no significant differences across temperatures or humidity levels, immediately after 30 min drying **(Supplementary Figure S8A)**. Humidified chamber-dried samples retained more water than those dried on a hotplate, and water content increased with higher humidity and lower temperature **(Supplementary Figure S8B)**.

Direct comparison of the two drying methods showed no differences at lower humidity, but at 50% RH, humidified chamber samples retained significantly more water than hotplate-dried samples **(Supplementary Figure S8C)**. These results highlight the greater environmental sensitivity of the humidified chamber method compared to the more consistent drying achieved with the hotplate.

#### 2.2. Effects of water content on the stability of biomolecules in dry trehalose glasses, after immediate drying

Excess water in glasses can increase molecular mobility and promote degradation of client molecules, however low water levels can also be detrimental, making an optimal water balance essential[28,34]. To examine if water content affects the protective capacity of our dry trehalose glasses, we evaluated how water content correlates with the stability of nucleic acids and proteins under different drying methods, temperatures, and humidities, immediately after 30 min drying **(Figure 2; Supplementary Figures S9; S10; S11)**.

Correlation analyses varied one condition while keeping the other constant within each drying method. For temperature analyses, humidity was fixed, and for humidity analyses, temperature was fixed, allowing assessment of how changes in water content affect molecular protection.

DNA integrity was largely unaffected by residual water content across drying conditions, with only a slight decline at 44 °C on the hotplate **(Supplementary Figure 9A-L)**. RNA, which is more sensitive to water, showed a slight decline in integrity at 44 °C when dried in a humidified chamber, with higher humidity correlating with reduced RIN **(Supplementary Figure S10L)**. This trend was not seen at other temperatures, and overall, RIN values remained close to fresh controls **(Supplementary Figure 10A-K)**. These results demonstrate no correlation between DNA protection and water content. Similarly, for RNA our data indicates no strong correlation between RNA integrity of water content.

Unlike nucleic acids, protein stability was highly sensitive to water content. Under hotplate drying at low humidity (10% RH), both LDH and CS showed significant positive correlations between residual water and enzymatic activity across different temperatures **(Figure 2A,D)**. LDH showed improved preservation of activity at moderate humidity **(20% RH, Figure 2B**), whereas CS did not. This indicates that LDH’s stability is enhanced by retaining slightly higher moisture levels during drying. Similarly, under humidified chamber conditions at 50% RH, both enzymes exhibited strong positive correlations between water content and activity across all tested temperatures **(Figure 2I,L)**, consistent with previous findings that adequate water stabilizes enzymes by maintaining hydration shells, structural flexibility, and hydrogen bonding networks[28].

**Figure 2.**
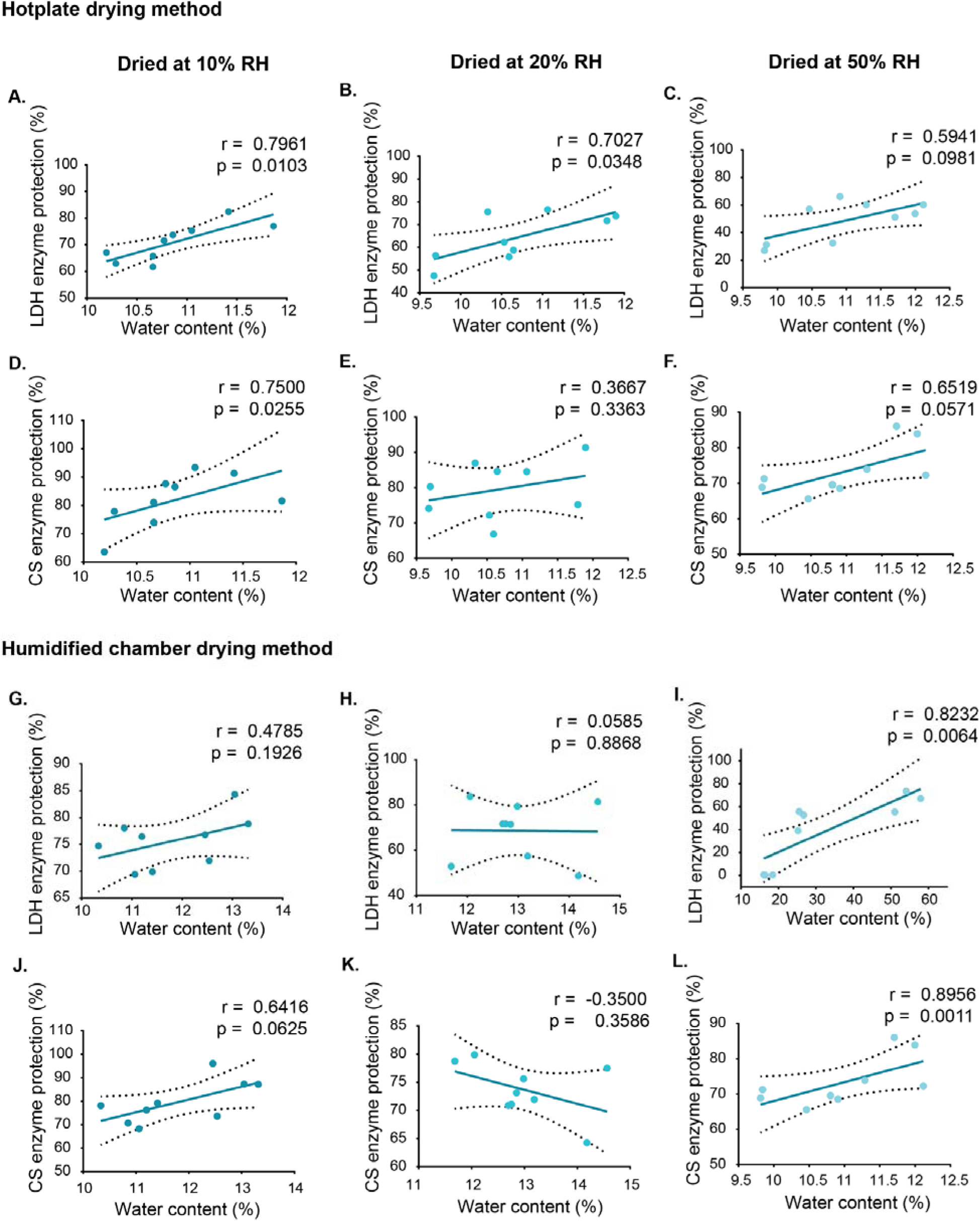
Relationship between residual water content and enzyme activity of LDH and CS under constant humidity and varying temperature conditions using two drying methods. Enzymatic activity of lactate dehydrogenase (LDH; A-C, G-I) and citrate synthase (CS; D-F, J-L) was assessed after drying under three different temperatures at each of the following constant relative humidity levels: 10%, 20%, and 50%. Drying was performed using either the hotplate method (A-F) or the humidified chamber method (G-L). Residual water content was measured in each sample post-drying and plotted against enzyme activity to evaluate their relationship. Correlation coefficients (r) and significance values (p) were calculated using Pearson correlation for normally distributed data and Spearman correlation for non-normally distributed data. Each data point represents an individual replicate. Dashed lines indicate 95% confidence interval (CI).

However, at high temperatures in the humidified chamber (37 °C and 44 °C), higher water content reduced the activity of both LDH and CS **(Supplementary Figures S11H,I; S11K,L)**. This is different from the positive effect seen when temperature varies at constant humidity. Excess moisture under these conditions may prevent proper glass formation and weaken the dried matrix, making enzymes more prone to inactivation. These results highlight that the effect of water on proteins is highly context-dependent: while moderate moisture can be protective, excess water under thermal stress can be detrimental.

These findings highlight that the effect of water content on stability is specific to different client molecule types (protein vs. nucleic acid), emphasizing the need to optimize drying protocols based on what client(s) are being preserved.

### 3. The effect of drying conditions on glass transition, and the role of water in determining glass transition temperature and the stability of biological molecules, after immediate drying

#### 3.1. Effect of drying method and environmental conditions on the glass transition temperature of trehalose glasses, after immediate drying

A glass transition is a physical change in which a vitrified material shifts from a hard, glassy state to a soft, rubbery state[60–62]. The glass transition temperature (Tg) refers to specific points within this transition range[28,34,41,43]. Here we report on three Tg values: onset, midpoint, and endset which represent the beginning, middle, and end of the transition range, respectively, in each drying regime after 30 min of drying. Ultimately, Tg values reflect the temperature/energy needed to induce molecular mobility and serve as a key indicator of the physical stability of dry systems, with higher Tg typically considered to be more protective to biological macromolecules[28,34,63].

In this study, Tg was strongly affected by drying methods and conditions. Hotplate drying produced higher Tg values (onset, midpoint, and endset) under lower humidity and higher temperatures, indicating more stable glassy states **(Supplementary Figure S12A-C)**. In contrast, under humidified chamber drying at 50% RH, Tg values were not detectable, likely due to high water content **(Supplementary Figure S12D-F)**. Overall, Tg was consistently lower in humidified chamber samples than hotplate-dried samples **(Supplementary Figure S12G-I)**. These results highlight that trehalose vitrifies with a higher resulting Tg when dried under low humidity/high temperature conditions.

#### 3.2. Effect of water content on glass transition temperature in dry samples, after immediate drying

In many glasses, including trehalose glasses, water acts as a plasticizer lowering Tg and increasing molecular mobility, which can reduce the stability of embedded molecules[64–67]. Because of the importance of water content on glassy properties, we assessed how residual water content affects Tg in samples dried by hotplate and humidified chamber methods, while varying temperatures at constant humidity as well as varying humidity at constant temperature, immediately after drying.

Under hotplate drying at 50% RH, higher water content showed a significant correlation with lowered Tg values, consistent with plasticizing effect of water[64–67] **(Supplementary Figure S13C,F,I)**. However, at fixed temperatures in both drying methods, correlations between water content and Tg were generally weaker. Notably, under certain humidified chamber conditions (37 °C/10-20% RH), higher water content was associated with slightly increased Tg, suggesting an anti-plasticizing effect[28] **(Supplementary Figure S14N,Q)**. These results highlight that water content strongly affects trehalose glass formation, with its impact (plasticization or anti-plasticization) depending on the drying conditions. Overall these results demonstrate that careful control of both temperature and humidity is essential to achieve consistent Tg, optimal vitrification, and sample stability.

#### 3.3. Effect of glass transition temperature on molecular stability, after immediate drying

After seeing that water content can affect Tg, we next wanted to assess how changes in Tg affect stability of diverse clients embedded within the glassy matrix. To assess whether Tg predicts molecular stability, we examined correlations between Tg and DNA/RNA integrity and LDH/CS activity across the described drying conditions, after immediate drying.

DNA **(Supplementary Figures S15; S16)** and RNA **(Supplementary Figures S17; S18)** remained stable across all conditions and showed no correlation with glassy properties, except for a slight decrease in RNA integrity at a fixed temperature of 44 °C across all humidity levels in the humidified chamber **(Supplementary Figure S18M)**.

Correlation analyses of Tg and protein stability revealed a complex, condition-dependent relationship: LDH and CS showed different sensitivities to Tg changes under varying temperature and humidity during hotplate and humidified chamber drying.

LDH under hot-plate drying at constant humidity maintained high activity even at lower Tg (20% and 50% RH; **Figure 3B,C,E,F,H,I)**, suggesting that protein protection does not increase with Tg. At constant temperatures, LDH stability was higher at 30 °C and 37 °C (**Figure 3J,K,M,N,P,Q**), supporting the classical view that a glassy matrix provides protection[28,34,63]. In contrast, no clear correlations between Tg and protection of proteins were observed with the humidified chamber **(Supplementary Figure S19A-O)**.

**Figure 3.**
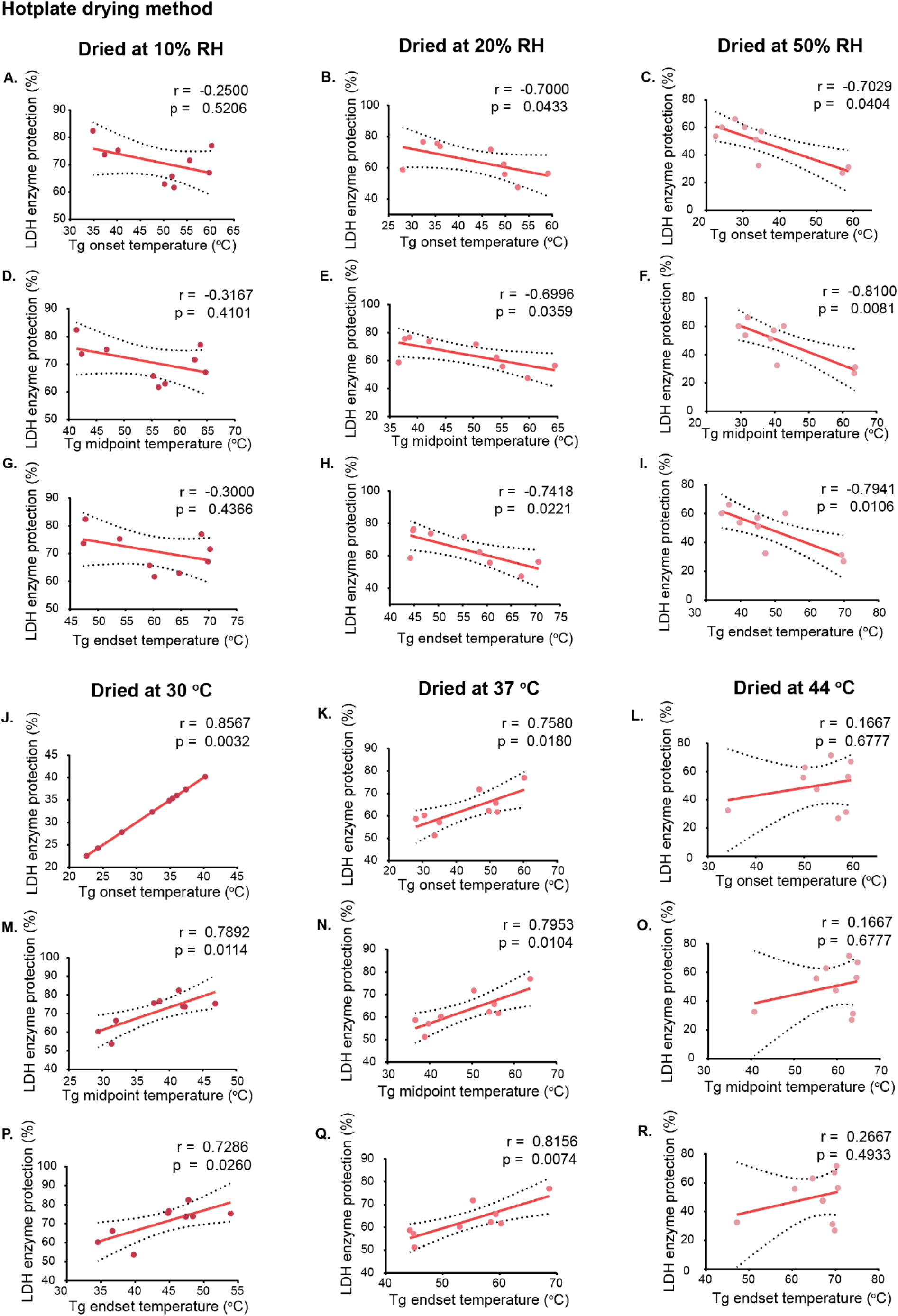
Correlation between glass transition temperature (Tg) and LDH activity in samples dried using the hotplate method under constant humidity and temperature conditions. Lactate dehydrogenase (LDH) activity was assessed in samples dried using the hotplate method. A-I show correlations between Tg values (onset, midset, and endset) and LDH activity under constant humidity conditions (10%, 20%, and 50% RH) at varying temperatures, while panels J-R show correlations under constant temperature conditions (30 °C, 37 °C, and 44 °C) at varying humidity levels. Glass transition temperatures were measured for each dried sample and plotted against LDH activity to assess their relationship. Correlation coefficients (r) and significance values (p) were calculated using Pearson correlation for normally distributed data and Spearman correlation for non-normally distributed data. Each data point represents an individual replicate. Dashed lines indicate 95% confidence interval (CI).

Under hotplate drying at constant humidity, CS exhibited a significant negative correlation with Tg onset at 10% RH and with both Tg onset and Tg midpoint at 20% RH **(Supplementary Figure S20A,B,E**). At constant temperatures, a positive correlation between Tg values and CS activity was observed at 30 °C, indicating improved stability with higher Tg under this condition **(Supplementary Figure S20J,M,P)**. However, no correlations were found at 37 °C or 44 °C. In the humidified chamber, correlations were generally weak, with only Tg endset showing a negative correlation at 10% RH and no significant correlation across temperatures **(Supplementary Figure S21E,G-O)**.

Overall, Tg was generally not a consistent predictor of protein stability, but higher Tg was mostly associated with better protection, influenced by humidity, temperature, drying method, and protein type. At high humidity, lower Tg improved protection, whereas at constant temperatures, higher Tg supported stability, consistent with the vitrification hypothesis. Significant correlations were mainly seen with hotplate drying, where rapid water removal influenced outcomes. CS stability correlated less with Tg than LDH, likely due to its higher resistance to drying stress and tendency to aggregate rather than unfold[68–70].

### 4. Effect of drying methods and conditions on glass former fragility and biomolecular protection, after immediate drying

#### 4.1. Effect of drying methods and conditions on glass former fragility, after immediate drying

Glass former fragility (m-index), classically described in relation to cold-induced vitrification, reflects how rapidly viscosity changes near Tg[35,42,71].

In our study, glass former fragility remained consistent across temperatures and humidity levels with hotplate drying **(Supplementary Figure S22A)**, and no significant differences were seen in the humidified chamber at 10 and 20% RH, immediately after drying **(Supplementary Figure S22B,C)**. At 50% RH, glass former fragility could not be determined due to high residual water preventing glass formation **(Supplementary Figure S22B,C)**. Overall, drying method and temperature had little impact on glass former fragility under low to moderate humidity during short-term drying, while high humidity hindered glass formation and assessment of glassy properties.

#### 4.2. Impact of glass former fragility on molecular stability across different drying conditions, after immediate drying

To assess whether glass former fragility predicts molecular stability, we examined its correlation with nucleic acid integrity and enzyme activity under different drying conditions, after immediate drying. Although higher glass former fragility is generally expected to reduce protection due to increased molecular mobility near Tg[28,29], no correlation was found with DNA **(Supplementary Figure S23)** or RNA **(Supplementary Figure S24)**. For proteins, a positive correlation was observed only for LDH at 37 °C under hotplate drying **(Supplementary Figure S25G)**, while CS showed no correlation **(Supplementary Figure S26)**. These results suggest that glass former fragility alone is not a reliable predictor of biomolecular stability and must be interpreted in the context of drying conditions, residual water, drying time, and biomolecule-specific sensitivity.

### 5. Effect of initial drying conditions and trehalose on long-term stability in trehalose glasses

To assess the long-term stability of dried biological materials, we conducted an aging experiment to evaluate how well the initial drying conditions preserve molecular integrity. Aging studies are important because short-term protection does not always reflect how well samples will last. Monitoring biomolecular stability helps assess the robustness of the vitrified state and its potential for room-temperature storage.

For the aging experiment, three conditions giving the highest enzyme protection after immediate 30 min drying were selected, as proteins are more sensitive than nucleic acids: 37 °C at 20% RH for LDH (highest fold protection; **Supplementary Figure S4C)**, 44 °C at 50% RH for CS (highest fold protection; **Supplementary Figure S4D)**, and 30 °C at 10% RH for both enzymes (protection; **Supplementary Figure S7A)**. The hotplate was chosen over the humidified chamber for its faster drying, which reduces exposure to intermediate moisture that can promote protein unfolding and ensures consistent glass formation[45]. To protect biomolecules during drying and storage, trehalose was used as a vitrifying mediator. Samples, prepared both with and without trehalose, were dried for 30 minutes and stored in LiCl desiccators at 10% RH at room temperature for up to 8 weeks, matching the residual water content **(Supplementary Figure S8A)** of the dried samples.

#### 5.1. Effect of initial drying conditions and trehalose on the long-term preservation of nucleic acid integrity

To evaluate how storage conditions affect nucleic acid integrity, we monitored both DNA and RNA samples over 8 weeks under different drying conditions and in the presence or absence of trehalose. DNA remained stable for 8 weeks, with no significant changes in integrity across drying methods or trehalose presence (**Figure 4A,B**). Minor fluctuations appeared in individual conditions **(Supplementary Figure S27A-F)**, but overall structure was well preserved.

**Figure 4.**
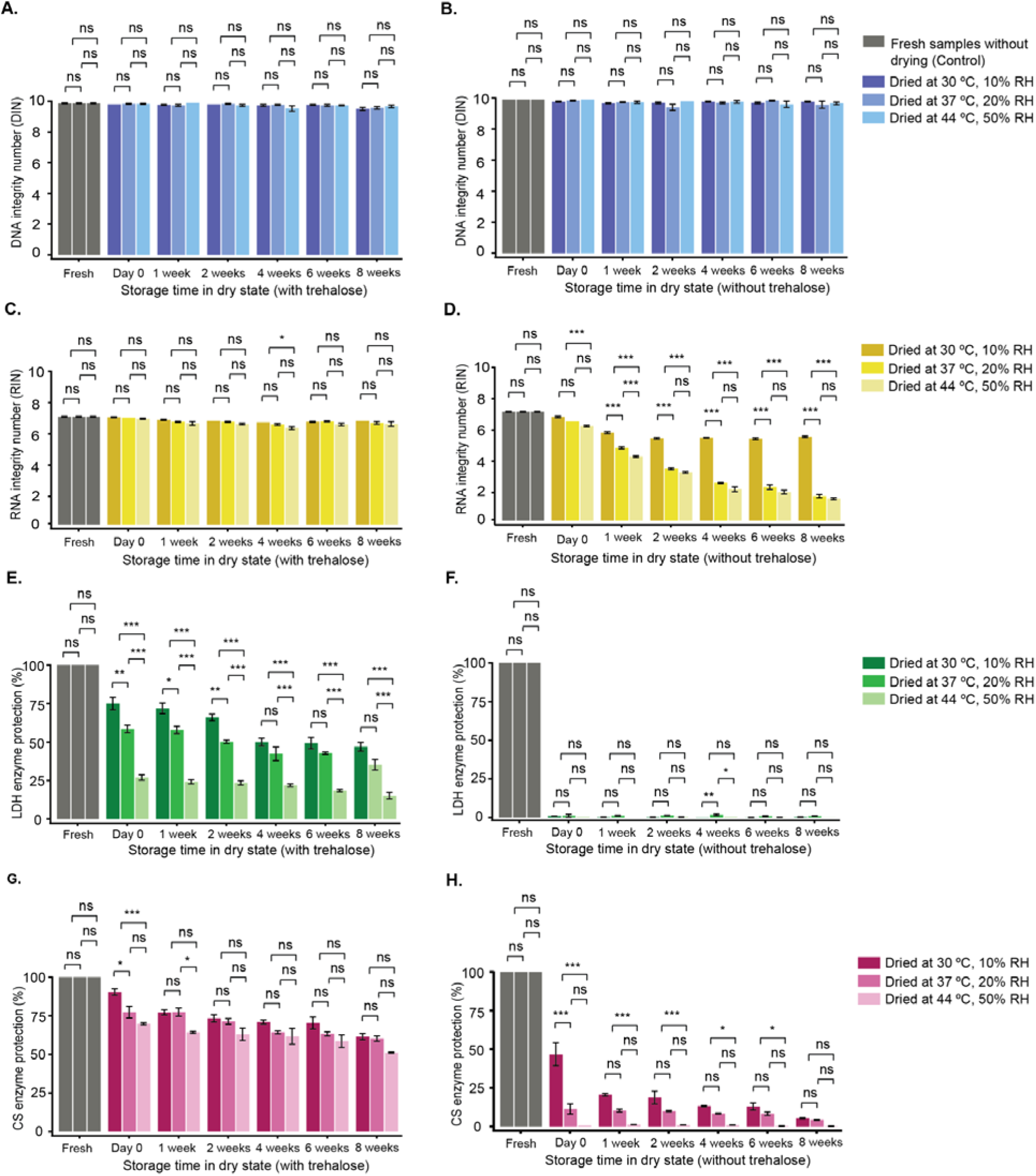
Molecular stability during storage under different drying conditions in the presence and absence of trehalose. DNA (A,B), RNA (C,D), lactate dehydrogenase (LDH; E,F), and citrate synthase (CS; G,H) samples were dried using the hotplate method under three conditions: 30 °C at 10% relative humidity, 37 °C at 20% relative humidity, and 44 °C at 50% relative humidity. Samples were stored at room temperature in a 10% RH LiCl jar, and molecular stability was evaluated at Day 0, Week 1, Week 2, Week 4, and Week 8. A, C, E, and G show samples dried and stored with 300 mM trehalose; B, D, F, and H show samples without trehalose. Nucleic acid integrity and enzyme activity were assessed at each time point. Data represent the mean ± SE from three independent experiments. A two-way ANOVA, followed by multiple comparisons, was performed to assess the effects of drying condition and storage time. However, only the statistical comparisons among the three drying conditions at each individual time point are presented graphically (significance set at p < 0.05).

However, RNA showed slightly greater sensitivity to both factors. In the presence of trehalose, RNA was generally well maintained across all conditions, with only minor differences at 4 weeks where 30 °C and 10% humidity samples had higher RIN values than those at 44 °C and 50% humidity (**Figure 4C**). Without trehalose, differences were evident by week 1: samples dried at 30 °C/10% RH remained most stable over 8 weeks, while higher temperature and humidity conditions (37 °C/20% RH and 44 °C/50% RH) led to gradual declines, highlighting trehalose’s protective role and the importance of initial drying conditions(**Figure 4D**).

Overall, these results show that DNA integrity is largely unaffected by drying conditions or trehalose over 8 weeks, whereas RNA stability is moderately influenced by both. This indicates that trehalose and initial drying parameters play a greater role in preserving RNA than DNA under the conditions tested.

#### 5.2. Effect of initial drying conditions and trehalose on the long-term preservation of protein activity

To evaluate how drying conditions and trehalose affect enzyme activity in the dry state, we measured LDH and CS activity over 8 weeks of storage.

When LDH activity was measured across the three drying conditions in the presence of trehalose, LDH activity was highest at Day 0 in samples dried at 30 °C/10% humidity, followed by 37 °C/20%, and lowest at 44 °C/50% (**Figure 4E**). These differences persisted through week 2, but by week 4 and through week 8, activity in the 30 °C/10% and 37 °C/20% groups became similar, both showing good long-term protection, while 44 °C/50% remained the lowest (**Figure 4E; Supplementary Figure S27M-O)**. Without trehalose, LDH activity dropped to near zero after drying (**Figure 4F; Supplementary Figure S27P-R)**, and minor fluctuations during storage likely reflect handling variability rather than true degradation.

Similar to LDH, CS activity declined over time under all drying conditions, but trehalose slowed this decline (**Figure 4G,H; Supplementary Figure S27S-U)**. Day 0 activity was highest at 30 °C/10%, and by Week 1, only 44 °C/50% samples showed noticeable decline. From Week 2 onward, CS activity was comparable across conditions with trehalose. Interestingly, without trehalose, CS still retained low but measurable activity, indicating greater inherent stability than LDH during desiccation and storage (**Figure 4H; Supplementary Figure S27V,W)**.

These results demonstrate that trehalose provides substantial protection for both enzymes, though the extent depends on the initial drying method. CS is inherently more stable than LDH, and long-term dry-state stability is influenced by both the enzyme itself and the drying conditions.

### 6. Effect of initial drying conditions on residual water content and its role in modulating molecular protection during storage

#### 6.1. Influence of initial drying conditions on long-term water retention during storage

Maintaining optimal water content during storage is crucial[37,44,49]. Too little water weakens hydrogen bonding and destabilizes the material, while too much water increases molecular mobility, making biomolecules more prone to damage[58,63,72]. Water also directly influences the glassy properties of the dried material-affecting parameters such as Tg and glass former fragility-which in turn impact stability and protective capacity[28,34,38,59]. Therefore, water content was monitored throughout storage to track changes over time.

At Day 0, immediately after drying water content ranged from 10–11% across all drying conditions **(Supplementary Figures 28A-D, 29A)**. By Week 1, it dropped to ∼4.5% and remained stable for the remainder of storage **(Supplementary Figures 28A-D, 29B)**. These results indicate that drying removed most of the water initially, with only minor additional loss during early storage, after which water content stabilized irrespective of the drying method or storage duration.

#### 6.2. Effect of water content on molecular stability during long-term storage

To assess the effect of residual water on molecular stability, we measured DNA, RNA, and protein integrity over time and analyzed correlations with water content. Across all drying conditions, water levels showed no significant correlation with DNA (**Figure 5A-C**) or RNA integrity (**Figure 5D-F**), indicating that both remain largely stable for at least up to 8 weeks.

**Figure 5.**
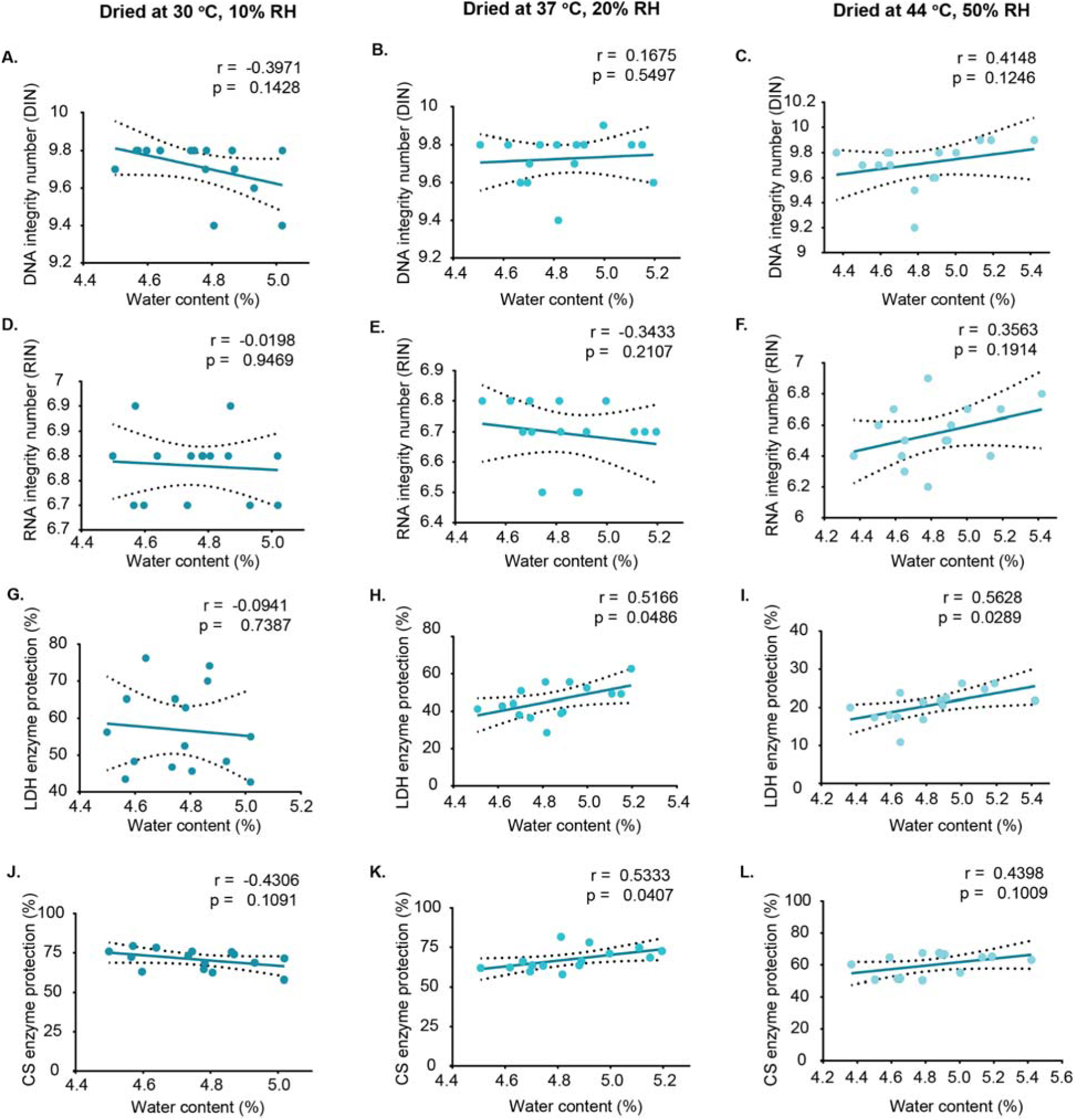
Correlation between residual water content and molecular integrity/activity in dried samples over storage. A-C show the correlation between water content and DNA integrity, D-F show RNA integrity correlations, G-I show lactate dehydrogenase (LDH) activity correlations, and J-L show citrate synthase (CS) activity correlations. Samples were dried using the hotplate method under three different drying conditions. Correlation coefficients (r) and significance values (p) were calculated using Pearson correlation for normally distributed data and Spearman correlation for non-normally distributed data. Each data point represents an individual replicate. Dashed lines indicate 95% confidence interval (CI).

In contrast, protein stability showed a variable relationship with residual water content (**Figure 5G-L**). For LDH, higher water content helped preserve activity in samples dried at 37 °C/20% and 44 °C/50% (**Figure 5H,I**), but no correlation was seen at 30 °C/10% (**Figure 5G**), suggesting that stability under milder conditions is less influenced by water content. For CS, a positive correlation with water content was observed only at 37 °C/20% (**Figure 5K**), with no significant correlation under the other conditions.

These results indicate that residual water has little impact on nucleic acid stability but can modulate protein stability, with the effect dependent on both the specific enzyme and the initial drying environment.

### 7. Effect of initial drying conditions on glass transition temperature and its role in molecular stability during long-term storage

#### 7.1. Influence of initial drying conditions on the long-term behavior of glass transition temperature during storage

Even dried materials that seem stable can change gradually over time[73]. Tg is a key indicator of the physical stability of dried biological materials during storage[28,34,74–77]. Keeping Tg above the storage temperature helps maintain a rigid structure and limits molecular motion, thereby slowing degradation[73,78]. Conversely, a decline in Tg signals reduced stability and increased risk of biomolecular damage[79]. Regular Tg measurements thus provide important insight into the physical state, quality, and shelf life of dried samples[74–77,79]. Here, we aimed to investigate how initial drying conditions influence the long-term behavior of Tg during storage.

At Day 0, immediately after drying, all samples showed broad glass transition events **(Supplementary Figure S29C)**. From Week 1, the Tg region narrowed, and a relaxation event appeared, indicating physical aging, consistently across all drying conditions **(Supplementary Figure S29D)**. At Day 0, significant differences in Tg midpoint and endset were observed only between samples dried at 30 °C/10% and 44 °C/50% (**Figure 6B,C**). From Week 1 onward, Tg values became similar across all drying conditions (**Figure 6A-C**). Tg was lowest at Day 0 and generally increased during storage, most clearly in samples dried at 30 °C/10%, which showed steady increases in onset, midpoint, and endset (**Figure 6A-C; Supplementary Figure S30A-C)**. Samples dried at 37 °C/20% showed variable changes in onset and midpoint, with stable endset values (**Figure 6A-C; Supplementary Figure S30D-F)**. In contrast, samples dried at 44 °C/50% showed minimal changes, suggesting limited physical alterations **(Supplementary Figure S30G-I)**.

**Figure 6.**
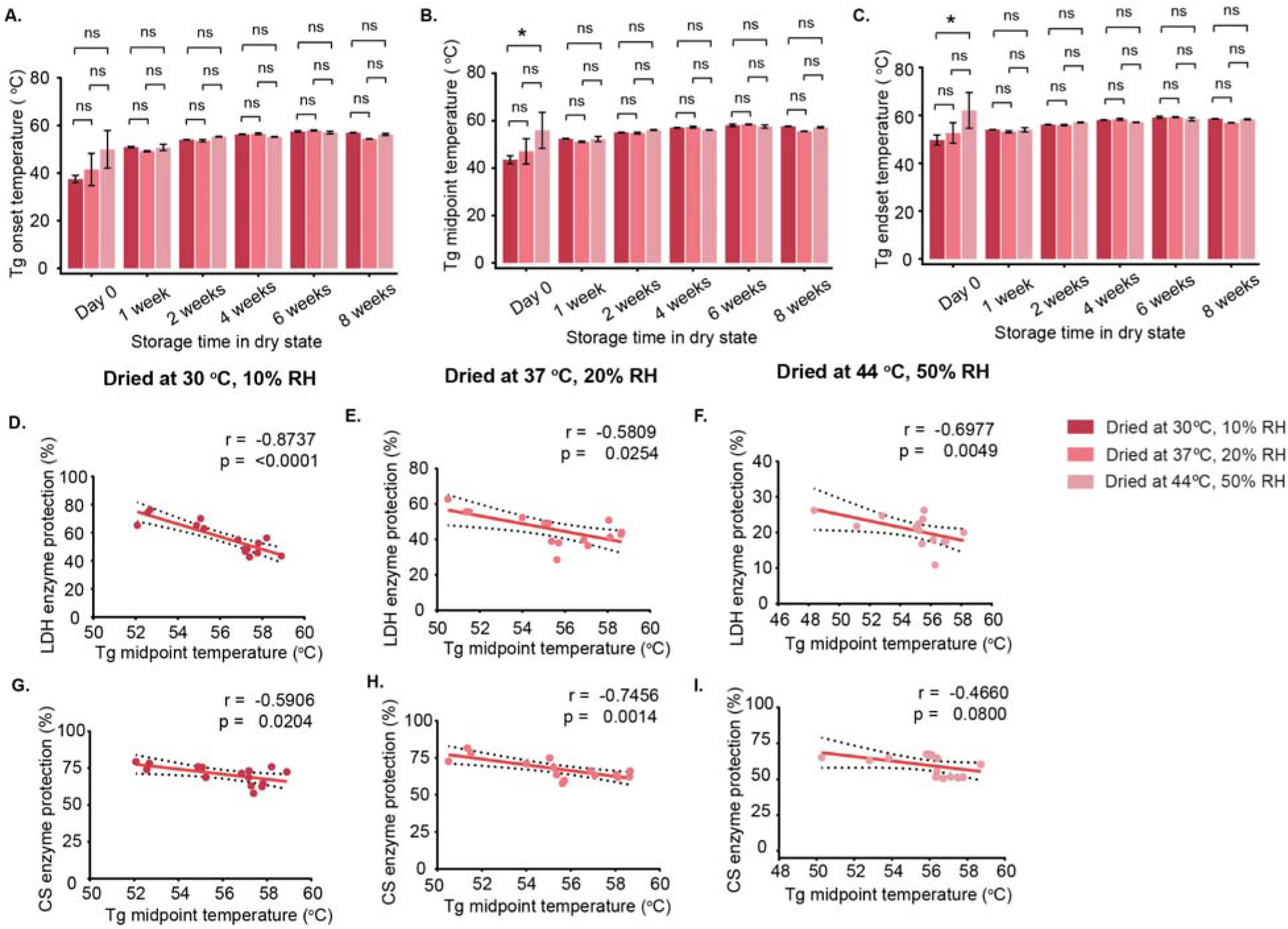
Glass transition temperatures (Tg) under different drying conditions and their comparison over storage time. Samples were dried using the hotplate method at 30 °C with 10% humidity, 37 °C with 20% humidity, and 44 °C with 50% humidity. A-C show comparisons of Tg onset (A), midpoint (B), and endset (C) values across all drying conditions at each storage time point. Bar graphs display mean Tg values with standard error (SE). Correlation analyses were conducted using only the Tg midpoint data to examine the relationship between Tg and enzyme activity for lactate dehydrogenase (LDH; D-F)) and citrate synthase (CS; G-L)). Correlation graphs include 95% confidence intervals (CI), and Pearson correlation was used for analysis. Statistical significance was determined at the 0.05 level.

These results show that while initial drying conditions affect Tg at Day 0, immediately after drying, storage under the same conditions leads to more uniform Tg values. The relaxation event following Tg indicates structural reorganization. Which suggests initial drying conditions influence the long-term behavior of Tg, with values gradually converging during storage due to structural reorganization.

#### 7.2. Impact of evolving glass transition temperature on molecular stability during long-term storage

Tg indicates how rigid and immobile a vitrified matrix becomes, with higher values generally reducing molecular motion and slowing degradation[28,34,63]. We assessed whether Tg correlates with long-term protection by comparing Tg values with the stability of nucleic acids and enzymes during storage.

Despite differences in Tg among samples, nucleic acids remained largely stable. DNA integrity **(Supplementary Figure S31)** and RNA stability **(Supplementary Figure S32)** showed no clear relationship with Tg, indicating that a higher Tg is not required to maintain nucleic acid stability in the dry state.

In contrast, LDH activity showed a strong negative correlation with Tg across all drying conditions (**Figure 6D-F; Supplementary Figure S33A-F)**. This does not mean higher Tg is harmful; rather, the activity decline is consistent with physical aging of the glassy matrix, as indicated by relaxation during storage **(Supplementary Figure S29D)**.

CS activity also negatively correlated with Tg, but significantly only under certain drying conditions **(30 °C/10% and 37 °C/20%; Figure 6G,H; Supplementary Figure S33G,H,J,K)**, with no correlation at 44 °C/50% (**Supplementary Figure S33I,L)**. These findings indicate that the relationship between Tg and enzyme stability is both enzyme-specific and dependent on the initial drying environment. Although high Tg is often associated with improved stability in vitrified systems, our results show that Tg alone does not fully predict long-term enzyme preservation, as factors such as matrix relaxation and aging during storage also play important roles.

### 8. Effect of initial drying conditions on glass former fragility, and their role in molecular stability during long-term storage

#### 8.1. Effect of initial drying conditions on glass former fragility during long-term storage

Assessing glass fragility during storage provides insight into how the physical properties of dried samples change over time, especially alongside shifts in Tg[28,29,34,35,42,71,74–77,79]. In our study, Tg shifted during storage, and the Tg region in DSC curves narrowed **(Supplementary Figure S29D)**, suggesting increased fragility and decreased structural stability[28,29,34,71]. To explore this, we evaluated glass former fragility for each drying condition at all storage time points.

Glass former fragility increased over storage time, showing distinct patterns depending on the initial drying conditions. Samples dried at 30 °C/10% increased in glass former fragility up to 4 weeks before stabilizing **(Supplementary Figure S34A)**. Samples dried at 37 °C/20% increased gradually until 6 weeks, then dropped suddenly **(Supplementary Figure S34B)**. Samples dried at 44 °C/50% showed a non-monotonic increase, peaking at 6 weeks before slightly decreasing **(Supplementary Figure S34C)**. Comparing all conditions, significant differences were only observed at 8 weeks between 37 °C, 20% RH and 44 °C, 50% RH samples **(Supplementary Figure S34D)**.

These results indicate that while initial drying conditions influence glass former fragility, storage time is the major factor affecting physical stability. Molecules gradually rearrange into a more stable state, but physical aging can increase glass former fragility due to molecular relaxation and heterogeneity. The sudden drop at 8 weeks remains unexplained, as longer storage was not tested.

#### 8.2. Impact of glass former fragility on molecular stability during storage

Given the role of glass former fragility in physical aging, we examined correlations between glass former fragility and molecular stability during storage. No significant correlation was observed for DNA **(Supplementary Figure S35A-C)** or RNA **(Supplementary Figure S35D-F)**.

In contrast, LDH and CS activity showed a strong negative correlation with glass former fragility in samples dried at 30 °C/10% **(Supplementary Figure S35G,J)**, but not in samples dried at 37 °C/20% or 44 °C/50%. This suggests that the initial drying condition strongly influences the glass former fragility profile.

### 9. Effect of aging on the relationship between Tg and glass former fragility

In this study we observed that in samples dried and assessed immediately for their protective capacity, higher Tg was generally correlated with increased protection (**Figure 3J,K,M,N,P,Q; Supplementary Figurers S20JMP; S21J,M,P)**, indicating that Tg can predict stability immediately after drying. Conversely, after prolonged storage, we observed that higher Tg did not correlate with protection (**Figure 6D-H; Supplementary Figure S33)**. To investigate why higher Tg did not consistently confer protection during aging, we examined the relationship between Tg midpoint and glass former fragility across all three drying conditions during storage.

Under mild drying conditions, strong and statistically significant positive correlations were observed: at 30 °C/10%, Tg midpoint and glass former fragility were highly correlated (r = 0.9337, p < 0.0001), and a similar correlation was seen at 37 °C/20% (r = 0.9195, p < 0.0001) **(Supplementary Figure S36A-B)**. In contrast, at 44 °C/50%, the correlation was weak and not statistically significant **(Supplementary Figure S36C)**, likely due to the glassy state being compromised at high temperature and humidity. These results indicate that under mild drying conditions followed by prolonged storage, higher Tg reflects structural compaction of the dried matrix, likely leading to increased glass former fragility and physical aging.

Overall, although higher Tg is generally linked to increased protection in dried systems, which we confirmed for samples left in the dry state for only a short time, our findings show that increased Tg during aging has an inverse relationship with the protection of enzymes and that this increased Tg is more indicative of physical aging of the glass, as confirmed by a concomitant increase in glass former fragility.

## Discussion

### 1. Impact of drying method and environmental condition on biomolecular stability in the presence of trehalose, immediately after drying

Our study demonstrates that the stability of biomolecules in the dry state depends on drying method, environmental conditions, and the presence of trehalose. DNA remained highly intact across all tested conditions, even without trehalose, reflecting its intrinsic chemical stability and resistance to short-term desiccation, with any minor structural changes during drying fully reversed upon rehydration[45,80]. This indicates that external protectants are not essential for preserving DNA under controlled drying and storage conditions[45,80].

In contrast, RNA was more sensitive to drying stress due to its single-stranded structure and labile phosphodiester backbone, making it prone to hydrolysis and structural degradation under heat and moisture[81–83]. Trehalose mitigated these effects by forming a glassy matrix that restricts molecular mobility, stabilizing local structure of RNA and reducing hydrolytic cleavage[81]. Under gentler humidified drying, RNA remained largely intact even without trehalose, highlighting that environmental stress dictates the necessity of protective mediators.

Proteins were the most vulnerable macromolecules. LDH lost nearly all activity without trehalose, whereas CS retained partial function, reflecting enzyme-specific sensitivities to structural perturbation and hydration. Trehalose stabilized both enzymes by restricting conformational flexibility and maintaining optimal moisture, thereby reducing hydrolytic, thermal, and oxidative damage. Protection also depended on the drying method and conditions, leading to protein destabilization both directly and indirectly through altered glassy properties.

These results establish a hierarchy of dry-state stability-DNA is largely resilient, RNA is moderately sensitive, and proteins are highly vulnerable[16,28,34,68,69,84]. Effective preservation relies on the interplay of trehalose and drying methods, which control water content, molecular mobility, and exposure to destabilizing intermediate states[45,85,86]. Optimizing these factors is essential for maintaining macromolecular integrity, and future work should address drying kinetics, rehydration stress, and hydrolytic damage for labile biomolecules[23].

### 2. Differential impact of residual water content on DNA, RNA, and protein stability across drying conditions, immediately after drying

In desiccation-tolerant systems, protection relies on maintaining small amounts of tightly bound water that limit molecular mobility and prevent damage, a process enhanced by mediators such as trehalose[23,37,63,87]. Residual water also determines key physical properties of the dried matrix, including Tg and glass former fragility, which govern biomolecular stability[28,34]. According to the water entrapment theory, trehalose forms a glassy matrix that traps minimal water, preserving local hydration and structural integrity without promoting degradation[63,88].

Our study demonstrates that drying methods and environmental conditions shape residual water content and thus influence macromolecular stability. DNA remained highly resistant to desiccation, while RNA showed modest sensitivity, likely due to hydrolysis of its labile backbone under higher water content[81,83]. Proteins were the most susceptible: moderate water helped maintain activity, and under harsher drying conditions, slightly higher water content buffered thermal and dehydration stress, supporting better enzyme preservation. Excessive water content increased molecular mobility[89] and promoted destabilization, while very low water caused stress-induced structural damage[90]. These results demonstrate that maintaining an optimal balance of residual water is essential for biomolecular stability, as both excess and deficiency can promote degradation[89,90]. Among the methods tested, hotplate drying provided more consistent control of water content, thereby reducing the destabilizing intermediate water and preserving protein structure[52] compared to the slow humidified chamber drying method. Together, these findings clarify how residual water, drying kinetics, and protective mediators interact, offering mechanistic insight into optimizing glassy-state preservation.

### 3. Drying techniques determine glass transition and preservation outcomes for biomolecules during short-term drying

Our findings show that Tg varies with drying method and environmental conditions, particularly temperature and humidity, reflecting the influence of residual water[34,58]. Hotplate drying generally produced higher Tg, especially under high-temperature, low-humidity conditions, favoring a stronger and more stable glassy matrix. In contrast, humidified chamber drying often resulted in lower Tg, likely due to higher retained water.

Water plays a dual role in dried systems: small amounts of tightly bound water help maintain structural integrity[56], while excess water acts as a plasticizer, increasing molecular mobility and lowering Tg[34,66,67]. In some cases, residual water may even enhance intermolecular interactions and raise Tg, a phenomenon known as anti-plasticization, depending on its amount, binding state, and distribution[28,65,91]. These findings show that water can either weaken or strengthen the glassy matrix, depending on how it is incorporated during drying.

We found that Tg had little effect on nucleic acids, which remained stable during short-term drying, indicating their preservation relies on intrinsic structural resilience rather than matrix rigidity. In contrast, protein stability varied with conditions. Higher Tg values were associated with improved protection due to reduced molecular mobility. However, LDH remained stable even in lower Tg, high-humidity environments, consistent with stabilization by hydration shells[63,92].

These results emphasize that protein stability depends on the interplay between Tg, residual water, and intrinsic protein properties such as structure and hydration requirements.

### 4. After short-term drying, glass former fragility reflects the drying conditions but does not reliably predict biomolecular stability

Since glass former fragility is a property associated with glass formation[35,42,71], it could not be determined at 50% humidity, where excess water inhibited vitrification[37,93]. Under low to moderate humidity, glass former fragility was largely unaffected by drying method or temperature. Its influence on stability was molecule-specific: DNA and RNA remained stable regardless of glass former fragility, reflecting their intrinsic resilience, whereas LDH stability improved with higher glass former fragility under hotplate drying at 37 °C, suggesting that a more flexible matrix can alleviate conformational stress or enable beneficial interactions with residual water. CS showed no clear trend, likely due to its tighter structural constraints and distinct hydration requirements. Overall, these results indicate that glass former fragility alone cannot universally predict biomolecular preservation; stability depends on the interplay between residual water, drying environment, and molecule-specific properties.

### 5. Biomolecular preservation during extended storage is influenced by the presence of trehalose and initial drying conditions

Our study demonstrates that the long-term stability of biomolecules during drying is governed by the interplay between drying conditions and the presence of trehalose. DNA showed remarkable resilience, with its double-stranded structure and chemical composition conferring strong protection against hydrolytic and oxidative damage, ensuring robust preservation under all tested conditions[94].

RNA stability is highly sensitive to both environmental conditions and storage duration. Its single-stranded structure and chemical lability make it prone to hydrolytic degradation over time[81]. Mild, controlled drying can reduce stress and partially preserve RNA even in the absence of trehalose, while trehalose further enhances stability by forming a glassy matrix that limits molecular motion. These observations highlight the combined importance of drying conditions, protective additives, and stable storage for maintaining RNA integrity.

Proteins are the most sensitive to drying-induced damage. Their stability depends on residual water, drying parameters, and trehalose. Sensitive enzymes such as LDH rapidly lose activity without trehalose, whereas more stable proteins like CS retain partial function under mild conditions. Trehalose enhances stability by limiting structural stress and conformational changes, and optimal preservation requires balancing water removal to reduce molecular mobility while maintaining sufficient hydration.

Together, these findings show that effective dry-state preservation arises from matching drying protocols and protectants to the molecular characteristics of each biomolecule. Trehalose stabilizes molecules while modulating mobility, and controlled temperature and humidity minimize stress and degradation. Understanding these mechanistic relationships enables rational design of drying strategies that maintain both structural and functional integrity, advancing practical room-temperature biopreservation.

### 6. Protein stability is affected by changes in water content over time, whereas nucleic acids are not

Our findings highlight the critical role of residual water in long-term biomolecular stability, with effects that are molecule- and condition-specific. Samples were stored at 10% relative humidity (using a LiCl desiccator) at room temperature, based on TGA (Thermogravimetric analysis) measurements taken immediately after 30 min drying[95]. Brief exposure to air during sampling, however, may have removed some water, so the actual residual content could have been slightly higher than that of the loaded samples[96]. Therefore, under these controlled conditions, additional water loss during the first week may reflect the equilibration of initially higher post-drying moisture along with the hygroscopic action of LiCl[95,97,98]. All samples-regardless of initial drying conditions-ultimately reached a similar moisture level during storage.

Nucleic acids remained highly stable, with their structural resilience and embedding in a trehalose glass limiting molecular mobility and protecting against stress. This indicates that DNA and RNA integrity in the dry state depends more on molecular architecture and trehalose stabilization than on residual water[45,81].

Proteins showed enzyme- and condition-specific responses. LDH preservation increased with residual water under harsh drying, indicating modest moisture helps maintain conformation, while under mild conditions it remained stable regardless of water. CS activity was less water-dependent, reflecting higher structural stability. These enzyme- and condition-specific patterns indicate that protein stability depends on residual water, drying conditions, and matrix formation, which together modulate molecular mobility and susceptibility to damage.

Collectively, these observations emphasize three points. First, even in seemingly stable low-humidity conditions, residual water can continue to equilibrate. Second, DNA and RNA are robust across varying hydration levels due to intrinsic structure and trehalose protection. Third, protein preservation depends on a dynamic interplay between residual water, initial drying stress, and matrix properties, highlighting the need to tailor drying protocols to each macromolecule’s specific sensitivities. Understanding these interactions provides a mechanistic framework for optimizing dry-state preservation, particularly for proteins whose stability is closely linked to both water content and drying conditions.

### 7. Aging enhances glass stability but compromises protein preservation in the dry state

Our findings highlight the complex role of Tg in the long-term preservation of dried biomolecules. Tg reflects the internal structure of the glassy matrix, and changes in Tg can signal shifts in molecular mobility caused by moisture uptake, phase separation, or chemical reactions[73,99,100]. Monitoring Tg therefore provides early insight into potential physicochemical changes that may precede functional losses, making it a useful tool for evaluating storage stability[78,101,102].

We observed prominent Tg changes in matrices dried under mild conditions, reflecting gradual compaction and reduced molecular mobility. In contrast, matrices dried rapidly at higher temperatures and humidity were more stable, showing minimal Tg changes during storage. Although higher Tg is often associated with improved stability, our results show that its effect is biomolecule-specific. DNA and RNA remained stable across a wide Tg range, reflecting their intrinsic structural resilience and minimal dependence on matrix rigidity. In contrast, proteins were more sensitive to matrix properties. Although higher Tg is often protective, excessively rigid matrices can reduce stability by trapping proteins in partially unfolded or misfolded conformations, limiting their ability to refold upon rehydration and diminishing activity, with additional loss occurring due to aging over time[26,28,34]. This effect was most pronounced for LDH, whereas more structurally stable proteins like CS were less affected.

These observations emphasize that Tg alone cannot predict biological preservation during long-term storage. Effective stabilization requires balancing matrix rigidity and molecular flexibility: the glass must be firm enough to protect biomolecules from degradation, yet flexible enough to allow conformational recovery. Achieving this balance depends on careful control of both initial drying conditions and storage environment to ensure that the protective matrix maintains function without imposing irreversible structural constraints.

### 8. Storage-induced changes in glass former fragility reveal selective biomolecular protections

Glass former fragility provides insight into how the mechanical and dynamic properties of a glassy matrix evolve during storage. Unlike Tg, glass former fragility describes how quickly a glass transitions with temperature, with highly fragile glasses undergoing the transition over a short temperature range, whereas stronger, less fragile glasses transition over a broader range[41,42]. During storage, the amorphous glass matrix underwent progressive molecular relaxation, stabilizing into lower-energy arrangements. This is reflected by increasing glass former fragility, indicating structural heterogeneity that can hinder uniform relaxation and affect biomolecular function[103].

In our study, glass former fragility was strongly influenced by the initial drying conditions. Samples dried at 30 °C and 10% humidity showed a rapid increase in glass former fragility before stabilizing, while samples dried under other conditions exhibited slower or more irregular shifts. This suggests that partial relaxation had already occurred during drying, highlighting that both the drying protocol and subsequent storage collectively shape the evolution of the glassy matrix.

Importantly, lower glass former fragility did not universally translate to improved biological protection. DNA and RNA remained stable across a wide range of glass former fragility values, reflecting their inherent resilience and protection by trehalose. Proteins, in contrast, were highly sensitive to glass former fragility, with their stability negatively correlated with glass former fragility and strongly influenced by the early drying conditions that set it. This highlights the critical role of early drying in controlling glass former fragility to preserve structurally sensitive proteins.

### 9. Tg positively correlates with protection during short-term dry storage, but is indicative of physical aging and loss of protection during long-term dry storage

Classically, increased Tg has been considered a desirable glassy trait positively correlated with increased protection in vitrified systems. Here we confirmed this finding, especially for enzymes dried and stored for short periods of time. However, for dry samples stored for longer periods, Tg was inversely correlated with protective capacity of the glassy system. The fact that increased Tg during long-term storage also correlated with higher glass former fragility indicates physical aging of the glass.

Physical aging, the slow, spontaneous, nonequilibrium relaxation of a glass to a lower energy and more compact state is known to lead to increased internal stress within molecules, especially proteins, promoting their deformation, loss of stability, and even aggregation over time. Our study provides a novel paradigm in which Tg is not always positively associated with protection in the glassy state and increased Tg over time is rather indicative of compaction of the vitrified matrix, a tell-tale sign of physical aging.

## Conclusion

This study demonstrates that the stability of biomolecules in the dry state is governed by the interplay between drying method, environmental conditions, and storage duration, which together determine the physical properties of the glassy matrix. While DNA remains largely stable across a range of drying and storage conditions, RNA shows moderate sensitivity, and proteins are severely affected by the conditions established during initial drying. Rapid hotplate drying can increase matrix rigidity and Tg, stabilizing the system, but overly rigid glasses may trap proteins in partially unfolded states, limiting recovery. In contrast, rapid yet mild drying reduces intermediate moisture exposure while preserving matrix flexibility, allowing proteins to retain functional conformations. During storage, gradual water loss leads to shifts in Tg and glass former fragility, which further affect RNA and protein stability in a molecule-specific manner, with the extent and direction of these changes depending on the initial drying conditions. These results highlight that effective room-temperature preservation requires carefully tuning drying and protectant formulations to balance matrix rigidity for stability with enough flexibility to allow functional recovery, particularly for sensitive proteins. Additionally, our study provides a new paradigm for conceptualizing the role of Tg in providing protection to client molecules embedded within trehalose glasses. Initially higher Tg is indicative of protection, but over time increasing Tg indicates the compaction and physical aging of the glass, which leads to internal strain, especially in proteins, and loss of client integrity.

## Materials and Methods

### Sample drying

A 300 mM trehalose solution (MW 378.33 g/mol;Avantor Science Control), a concentration commonly used for preserving complex biological samples[44,46], was tested using two drying methods. For each experiment, 50 μL of trehalose solution was placed in three-well glass slides (1 mm) and dried for 30 min, a short duration chosen to minimize degradation[45].

In the humidified chamber (ESPEC North America, model BTL-433) method, temperature is set according to experimental conditions. For the hotplate (Thermo scientific, model SP88857100) method, the hotplate is placed inside the humidified chamber, which is maintained at room temperature (22 °C), while the hotplate temperature is adjusted as needed. Humidity is controlled using the humidifier for both methods. Drying was performed across all combinations of three temperatures (30°C, 37°C, 44°C) and three relative humidities (10%, 20%, 50%), resulting in nine distinct conditions: 30°C/10%, 30°C/20%, 30°C/50%, 37°C/10%, 37°C/20%, 37°C/50%, 44°C/10%, 44°C/20%, and 44°C/50%. These conditions represent physiologically and practically relevant drying environments: 30 °C (mild stress), 37 °C (normal mammalian temperature), and 44 °C (elevated thermal stress), with relative humidities of 10% (very dry), 20% (moderately dry), and 50% (relatively moist). All nine conditions were tested with both drying methods, with or without trehalose, to assess glass formation and biomolecular stability, immediately after 30 min drying.

### Glass transition temperature (Tg) calculation

Dried samples (3-4 mg) were placed in pre-weighed pans, hermetically sealed, and analyzed by differential scanning calorimetry (DSC). Samples were heated from −40 °C to 250 °C at a rate of 10 °C/min to assess the glassy state of trehalose. Thermograms obtained after the full heating ramp were used to identify degradation, melting, and glass transition peaks. Tg onset, midpoint, and endset were determined using Trios software (TA Instruments TRIOS v5.0.0.44608), and all reported values represent the average of triplicate measurements.

### Glass former fragility (m-index) calculation

The glass former fragility (m-index) was calculated following the method originally described by Crowley and Zografi[104] and more recently adapted for desiccation by others[28,29,34]. It was determined using equations 10 and 14, which incorporate the activation enthalpies of structural relaxation at Tg (ΔEₜg) and for viscosity (ΔEη), the gas constant (R), and the experimental glass transition onset (Tg) and endset (Tg_off) temperatures. The mean of each triplicate set was calculated to ensure consistency.

### Calculation of water content in dried samples

This analysis aimed to assess thermal stability and the proportion of volatile components by tracking weight loss as temperature increases. Platinum crucibles (TA 952018.906) were tared before use. Dried samples (10-11 mg) were placed on thermogravimetric analyser (TGA) crucibles and heated from 30°C to 250°C at a rate of 10°C/min. Water loss was used to determine residual water content, and mass changes from ∼100°C to 180°C were analyzed in Trios “Smart Analysis” to identify inflection points and compare initial and plateau masses. Triplicates were analyzed, and average values reported.

### DNA Integrity Assay

Mouse genomic DNA (228 µg/mL; Promega; G309A) was used to assess DNA integrity using the Agilent TapeStation system. DNA (10 µL) was mixed with 90 µL of 300 mM trehalose in Tris-EDTA (Research Products International, pH 8.0) and split into two sets: one dried under nine temperature-humidity combinations using hotplate or humidified chamber, the other stored at 4°C for 30 min as control. Dried samples were rehydrated with 50 µL molecular-grade water (Sigma-Aldrich) and prepared for TapeStation analysis. For each run, 1 µL of sample was mixed with 10 µL genomic DNA sample buffer (Agilent) and loaded into TapeStation along with genomic DNA ScreenTape (Agilent)[105]. The system generated electropherograms and gel-like images, automatically calculating the DNA Integrity Number (DIN) from 1 (highly degraded) to 10 (intact DNA), with higher scores indicating better preservation. A genomic DNA ladder (1 µL) with 10 µL buffer was used for size reference; all conditions were tested with and without trehalose in triplicate.

### RNA Integrity Assay

Commercially available mouse liver total RNA (1.86 µg/µL; BioChain, R133414950) was used to assess RNA integrity with the Agilent TapeStation. RNA was diluted 1:10 in molecular-grade water, and each 10 µL sample was mixed with 90 µL of 300 mM trehalose or molecular-grade water and split into two sets: one dried under nine temperature-humidity combinations using hotplate and humidified chamber methods, the other stored at 4°C for 30 min as control. Dried samples were rehydrated with 50 µL molecular-grade water for TapeStation analysis. For each sample, 2 µL RNA was mixed with 1 µL high sensitivity RNA sample buffer (Agilent) and loaded onto the TapeStation with high sensitivity RNA ScreenTape (Agilent)[106]. The system generated electropherograms and gel-like images, automatically calculating the RNA Integrity Number (RIN) on a scale from 1 (highly degraded) to 10 (intact RNA), with higher scores indicating better-preservation. A 2 µL high-sensitivity RNA ladder (Agilent) with 1 µL buffer served as size reference; all conditions were measured in triplicate.

### Lactate dehydrogenase (LDH) enzyme protection assay

LDH activity assays were performed as described previously[19,28,34,68,84,107]. Stock solutions of 25 mM Tris-HCl (pH 7.0; MW 157.6 g/mol; US Biological), 100 mM sodium phosphate (MW 137.99 g/mol; Biomatik), 2 mM sodium pyruvate (MW 110.04 g/mol; TCI), and 10 mM NADH (MW 709.4 g/mol; Roche) were prepared and stored at 4°C. L-lactate dehydrogenase (LDH) from rabbit muscle (MW ∼140,000 g/mol; Sigma-Aldrich; 10127230001) was diluted to 1 g/L in 25 mM Tris-HCl. Trehalose samples were prepared by mixing LDH with trehalose at a 1:10 ratio. Each 100 µL sample (90 µL mixture + 10 µL LDH) was split into a 4 °C control and dried for 30 min under nine drying combinations using a hotplate or humidified chamber. After drying, control and experimental samples were rehydrated with molecular-grade water to a final volume of 250 µL. Absorbance was measured using a Nanodrop One^c^ spectrophotometer (Thermo Scientific) with quartz cuvettes (Hellma Analytics). Before measurement, the spectrophotometer was blanked with 980 μL of 100 mM sodium phosphate buffer containing 2 mM sodium pyruvate. Absorbance at 340 nm (A340) was recorded every 2 s for 60 s. LDH activity was determined by adding 10 μL of rehydrated sample and 10 μL of NADH to 980 μL of the buffer, and monitoring NADH oxidation at 340 nm. Protection efficiency (%) was calculated as (rehydrated/control reaction rate) × 100 and averaged from triplicates with and without trehalose.

### Citrate synthase (CS) enzyme protection assay

The CS enzyme assay was conducted as outlined previously[69]. Five milliliters of 5x CS assay buffer (Sigma-Aldrich) was diluted to 25 mL with molecular-grade water to make 1x buffer. Dilute the commercially available citrate synthase (CS) from porcine heart (MW 3486 g/mol; Biosynth; JAA02796) 1:5 in 1x CS buffer, then mix 10 µL of this diluted enzyme with either 90 µL of 300 mM trehalose or 90 µL of 1x CS buffer (without trehalose). Each 100 µL sample was split into a 4 °C control and a set dried for 30 min under nine temperature and humidity combinations using humidified chamber or hotplate methods. Dried samples were rehydrated with 50 µL molecular-grade water and assayed in triplicate to assess protection efficiency. While drying, prepare the necessary reagents by dissolving 5,5′-Dithiobis (2-nitrobenzoic acid) (DTNB, Sigma-Aldrich) in 1 mL of 100% ethanol and Acetyl CoA (Millipore) in 1 mL of molecular-grade water. Aliquot 160 µL into six tubes and store them at −20°C. When needed, thaw one aliquot of each, mix with 12.88 mL of 1x CS assay buffer, and store the solution at 4°C. Prepare the oxaloacetic acid (OAA) solution by dissolving 3.9 mg of dry OAA (Sigma-Aldrich) in 2 mL of 1x assay buffer, then dilute to a final volume of 3 mL and store at 4°C. A total of 2 µL of rehydrated sample was mixed with 188 µL of buffer, followed by the addition of 10 µL of OAA. Protection efficiency (%) was calculated as the ratio of experimental to control absorbance at 412 nm (A412) × 100, averaged from triplicates with or without trehalose.

### Evaluation of protection and glassy properties during long-term storage

Samples dried for 30 min were stored in a sealed lithium chloride (LiCl; MW 42.4 g/mol; Research Products International) desiccator at room temperature[44]. LiCl maintained a constant humidity equal to the final water content of the dried samples, as determined by TGA. Samples of DNA, RNA, LDH, and CS, with or without 300 mM trehalose, were stored for 1, 2, 4, 6, and 8 weeks, then rehydrated for analysis of nucleic acid integrity and enzyme activity relative to fresh controls. For glassy property measurements, 300 mM, 30-min dried trehalose samples were stored under the same conditions, with Tg, glass former fragility, and water content recorded at Day 0 and subsequent time points. All measurements were performed in triplicate.

### Statistical Analysis

Statistical analyses were performed using GraphPad Prism 10 and RStudio (R 4.2.2) to assess the effects of drying and storage conditions on biomolecular stability and glassy properties. One-way ANOVA with Tukey’s post hoc test was performed in RStudio to compare formulations and drying conditions at each time and to assess storage effects within each drying method. For the storage (aging) experiments with multiple time points and drying conditions, two-way ANOVA was performed in GraphPad Prism to assess the effects of drying condition, storage time, and their interaction on biomolecular integrity and physical properties. Correlation analyses were conducted to assess relationships between glassy properties and biomolecular stability. Depending on data distribution, Pearson or Spearman correlation tests were performed using GraphPad Prism.

All experiments were performed in triplicate, with significance set at p < 0.05 (*p < 0.05; **p < 0.01; ***p < 0.001). Error bars in bar graphs represent the standard error (SE), while correlation graphs display 95% confidence intervals (CI).

## List of Abbreviations

LDH: Lactate dehydrogenase
CS: Citrate synthase
RH: Relative humidity
DSC: Differential scanning calorimetry
TGA: Thermogravimetric analysis
DIN: DNA integrity number
RIN: RNA integrity number
Tg: Glass transition temperature
DTNB: 5,5’-Dithiobis(2-nitrobenzoic acid)
OAA: Oxaloacetic acid
EDTA: Ethylenediaminetetraacetic acid
NADH: Nicotinamide adenine dinucleotide
ANOVA: Analysis of variance
SE: Standard error
CI: Confidence interval

## Acknowledgments

The authors are thankful to all the members of the Boothby Lab for discussions and reading of this manuscript. This work was primarily supported by NSF DBI grant # 2213983 and the Institutional Development Award (IDeA) from the National Institute of General Medical Sciences of the National institute of Health (Grant #2P20GM103432).

## Author contributions

**U.G.V.S.S.K:** Conceptualization, Methodology, Investigation, Formal analysis, Data analysis and curation, Writing - Original Draft, Writing - Review and Editing, Visualization. **T.C.B:** Conceptualization, Methodology, Writing - Original Draft, Writing - Review and Editing, Supervision, Visualization, Project administration, Funding acquisition.

## Data availability statement

All raw data and supplementary figures used in this manuscript will be made public upon acceptance/publication.

## Additional information

The authors declare no competing interests.

---

**Supplementary Figure S1.**
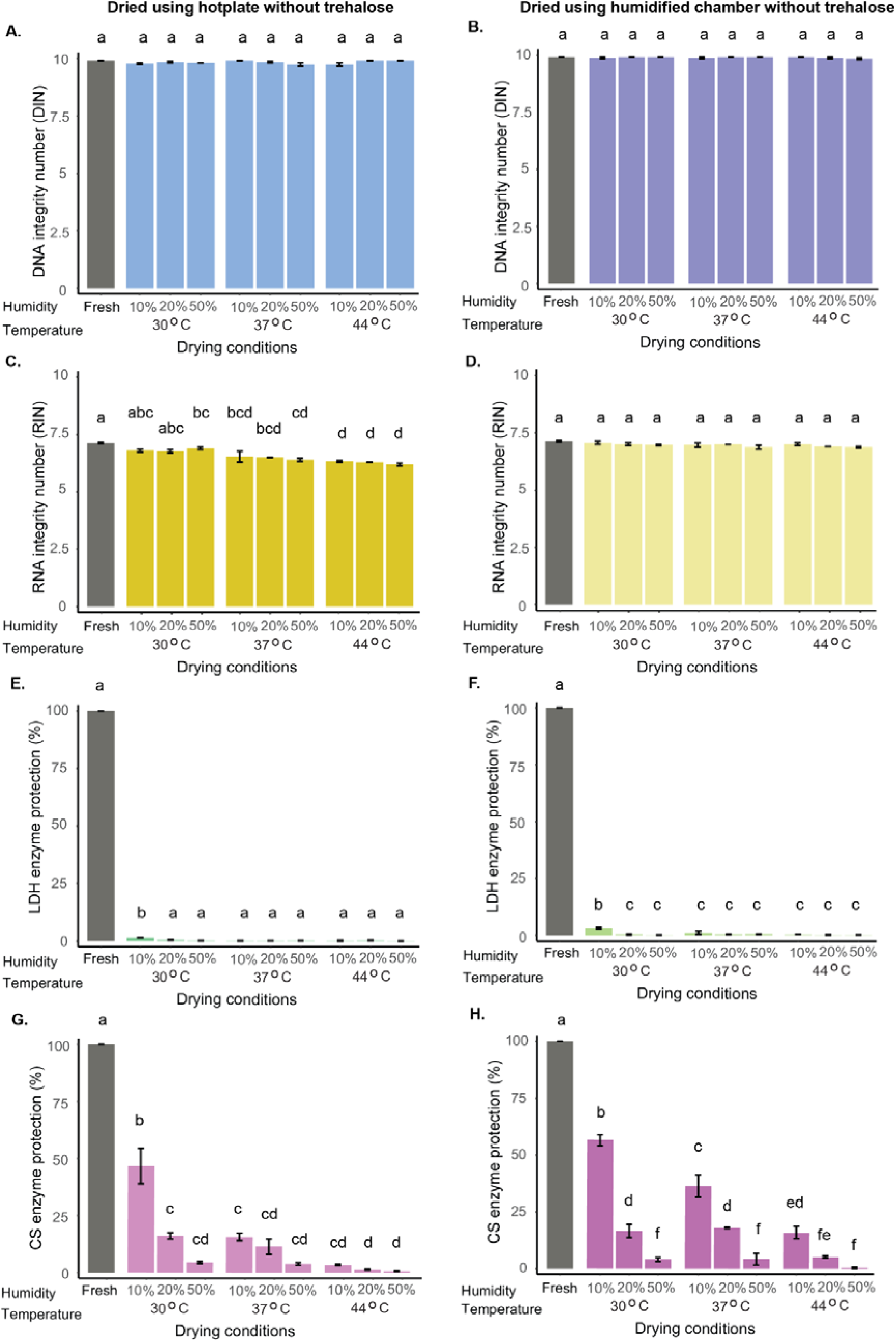
Differential stability of nucleic acids and proteins under various drying conditions in the absence of trehalose. DNA (A,B), RNA (C,D), lactate dehydrogenase (LDH; E,F), and citrate synthase (CS; G,H) samples were dried using two methods-hotplate (A,C,E,G) and humidified chamber (B,D,F,H)-across nine temperature-humidity combinations, all in the absence of 300 mM trehalose. DNA and RNA integrity were assessed using DNA Integrity Number (DIN) and RNA Integrity Number (RIN), respectively, while enzyme activity retention was used to evaluate protein function. Different letters indicate statistically significant differences among drying conditions within each panel, as determined by one-way ANOVA followed by Tukey’s post hoc test (α = 0.05). Data represent mean ± SE from three independent replicates per condition.

**Supplementary Figure S2.**
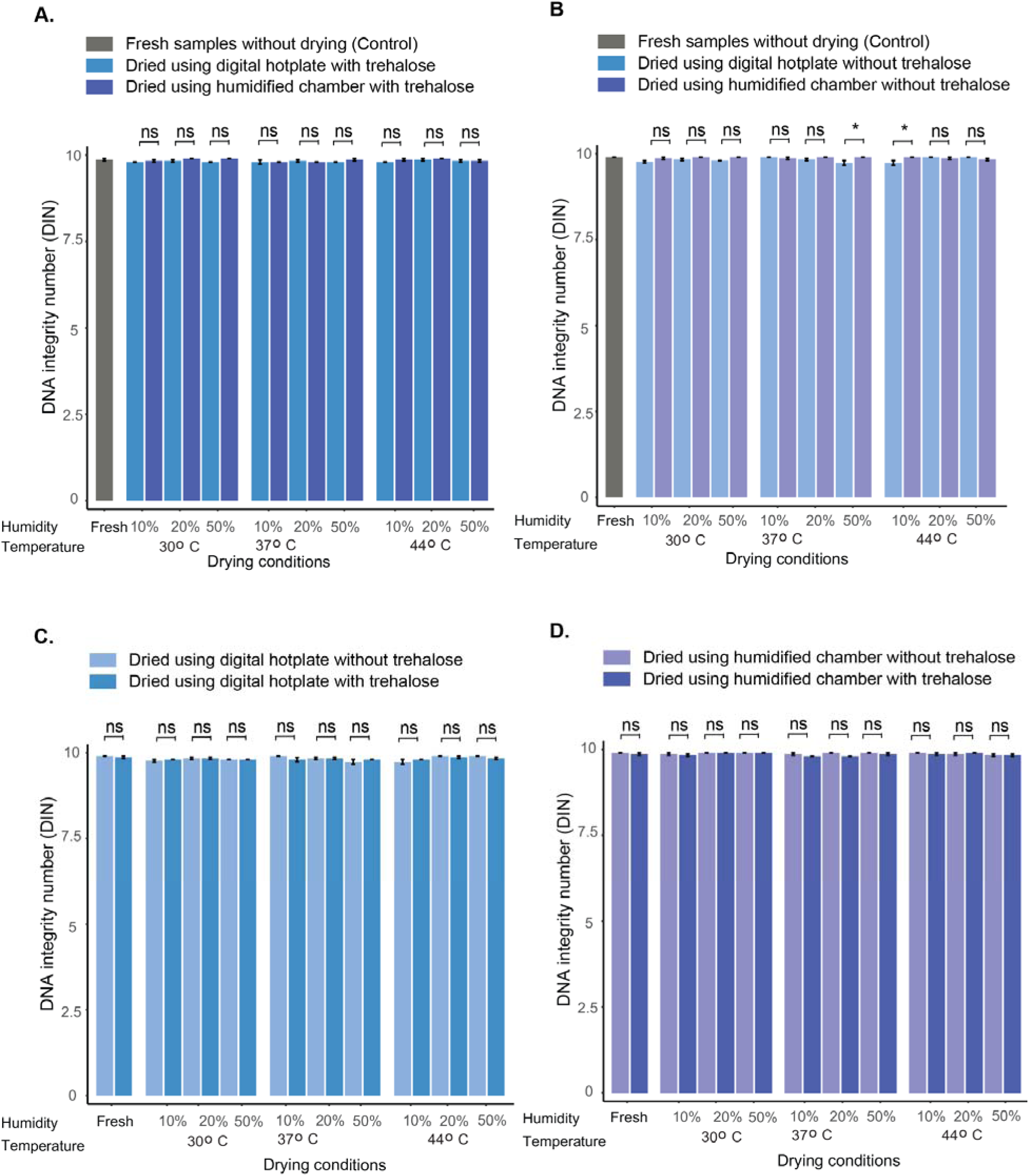
Effect of drying method and trehalose on DNA integrity. Comparison of drying methods with trehalose (A). Comparison of drying methods without trehalose (B). Hotplate drying with and without trehalose (C). Humidified chamber drying with and without trehalose (D). DNA integrity was assessed using the DNA Integrity Number (DIN). Statistical comparisons were made only between treatments within the same drying condition, as determined by one-way ANOVA followed by Tukey’s post hoc test (α = 0.05). Data represent mean ± SE from three independent replicates per condition.

**Supplementary Figure S3.**
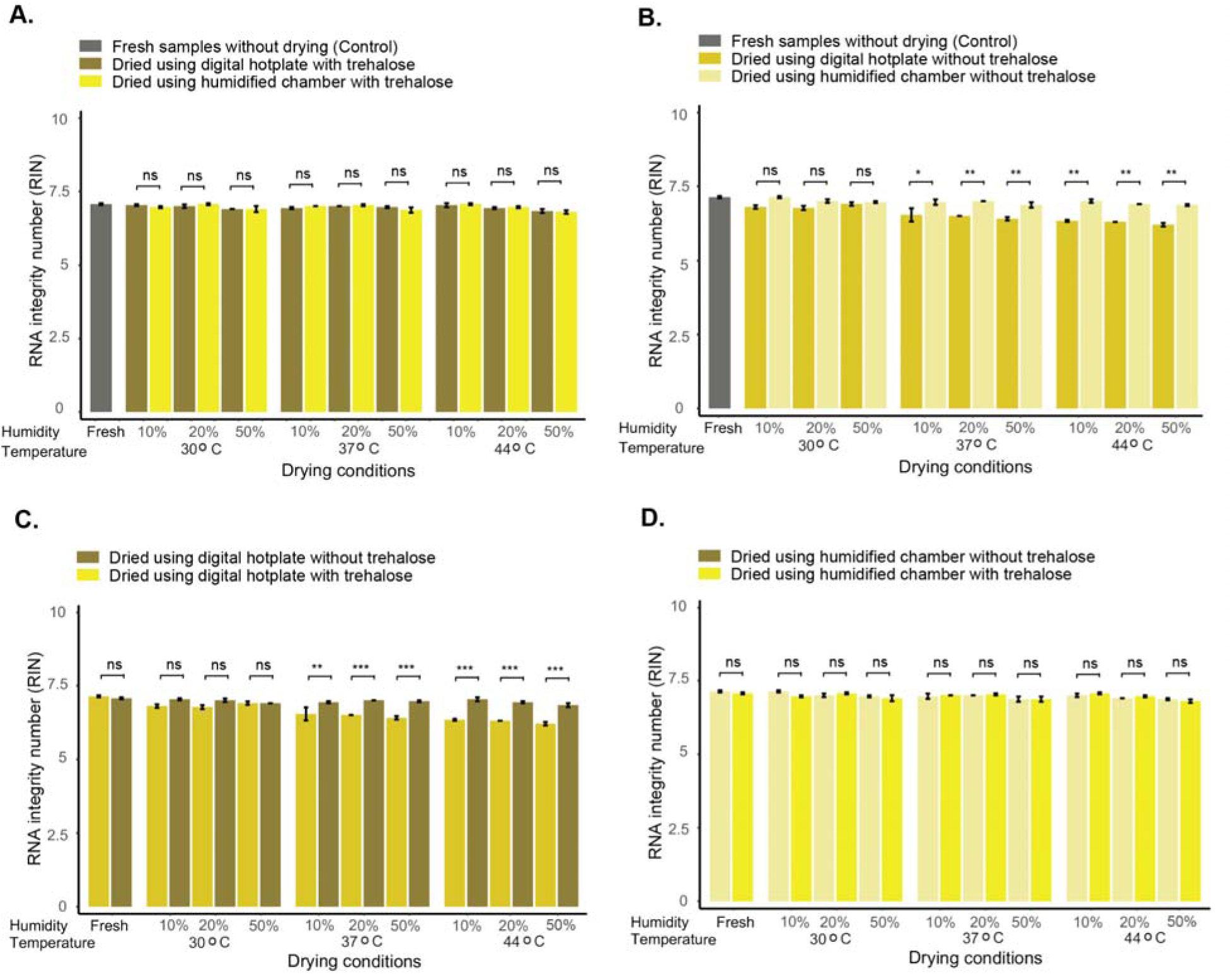
Effect of drying method and trehalose on RNA integrity. Comparison of drying methods with trehalose (A). Comparison of drying methods without trehalose (B). Hotplate drying with and without trehalose (C). Humidified chamber drying with and without trehalose (D). RNA integrity was assessed using the RNA Integrity Number (RIN). Statistical comparisons were made only between treatments within the same drying condition, as determined by one-way ANOVA followed by Tukey’s post hoc test (α = 0.05). Data represent mean ± SE from three independent replicates per condition.

**Supplementary Figure S4.**
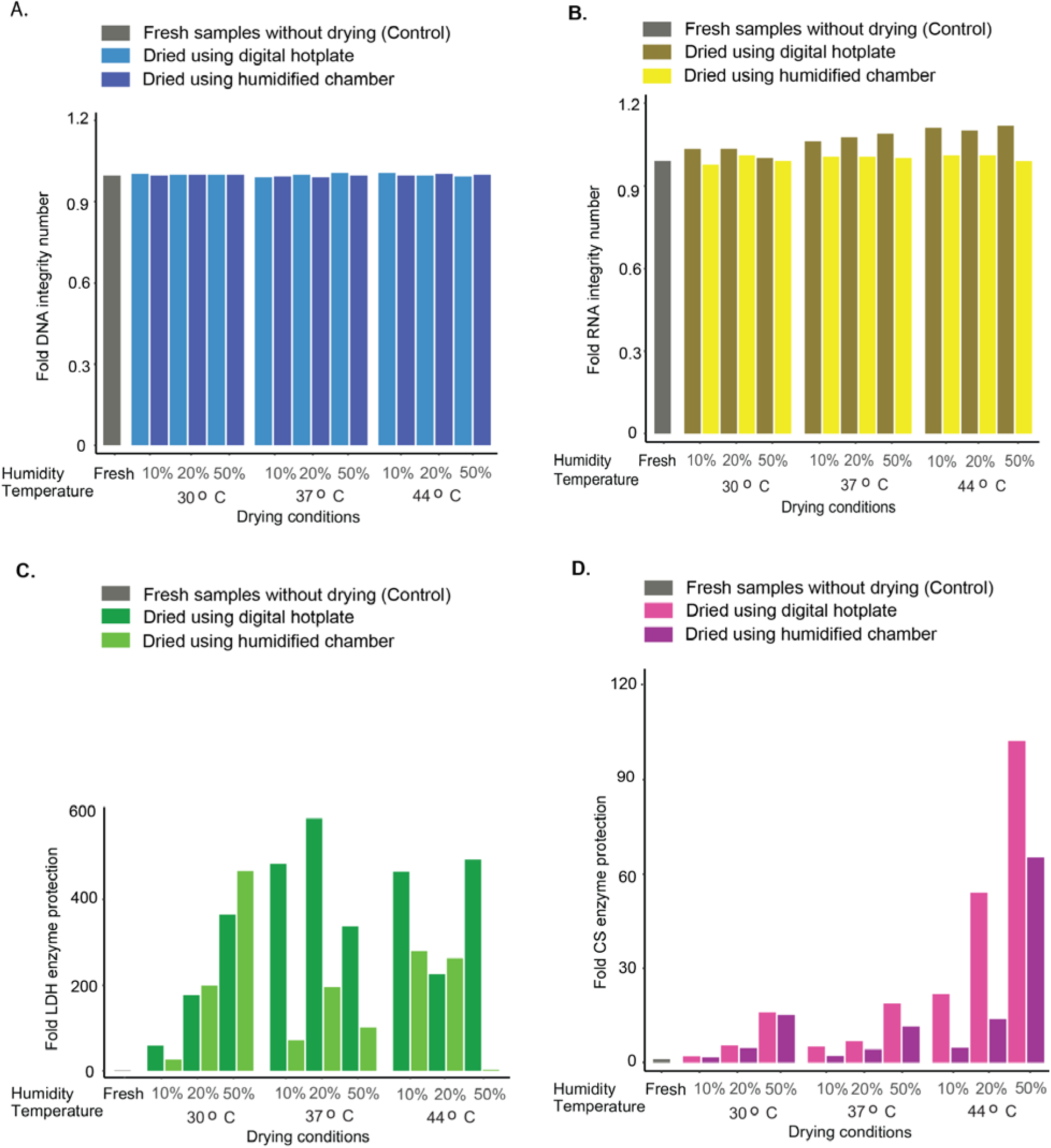
Fold protection of nucleic acids and proteins under different drying conditions. Fold protection of DNA (A), RNA (B), lactate dehydrogenase (LDH; C), and citrate synthase (CS; D) in both drying methods: hotplate and humidified chamber. Fold protection was calculated as the ratio of average biomolecular integrity/activity in samples dried with trehalose to that in samples dried without.

**Supplementary Figure S5.**
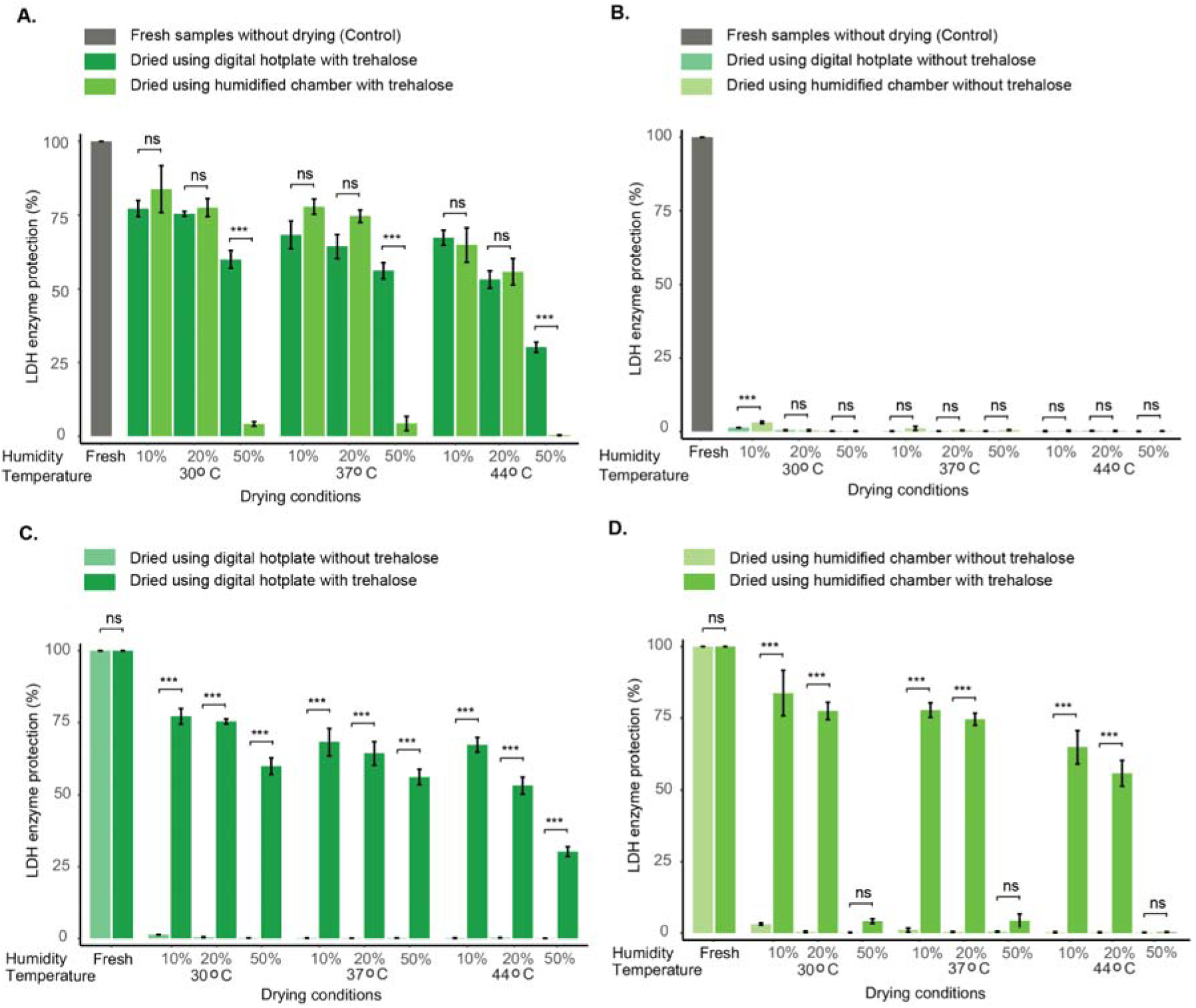
Effect of drying method and trehalose on LDH activity. Comparison of drying methods with trehalose (A). Comparison of drying methods without trehalose (B). Hotplate drying with and without trehalose (C). Humidified chamber drying with and without trehalose (D). Lactate dehydrogenase (LDH) activity was assessed using the lactate dehydrogenase enzyme protection assay. Statistical comparisons were made only between treatments within the same drying condition, as determined by one-way ANOVA followed by Tukey’s post hoc test (α = 0.05). Data represent mean ± SE from three independent replicates per condition.

**Supplementary Figure S6.**
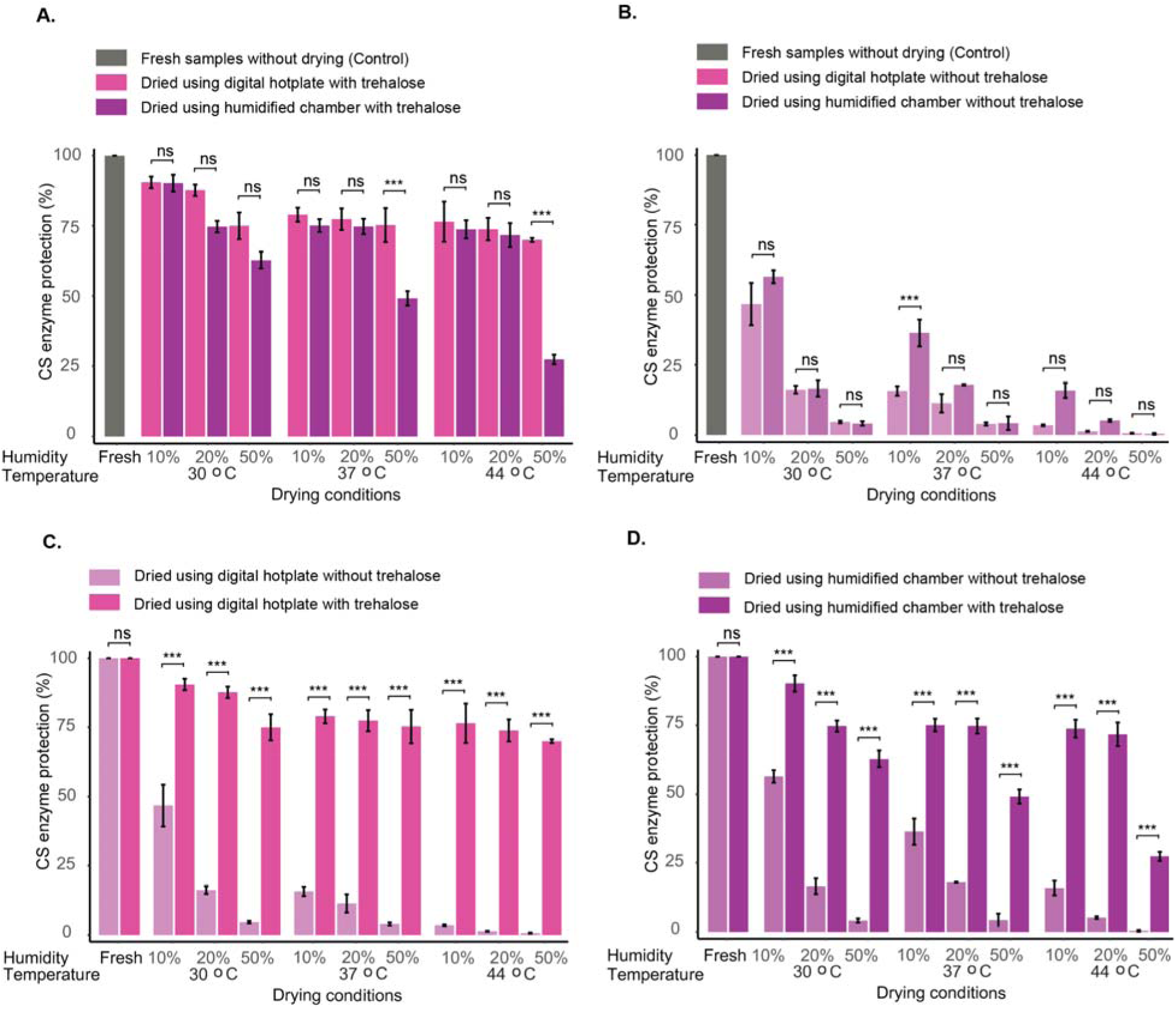
Effect of drying method and trehalose on CS activity. Comparison of drying methods with trehalose (A). Comparison of drying methods without trehalose (B). Hotplate drying with and without trehalose (C). Humidified chamber drying with and without trehalose (D). Citrate synthase (CS) activity was assessed using the citrate synthase enzyme protection assay. Statistical comparisons were made only between treatments within the same drying condition, as determined by one-way ANOVA followed by Tukey’s post hoc test (α = 0.05). Data represent mean ± SE from three independent replicates per condition.

**Supplementary Figure S7.**
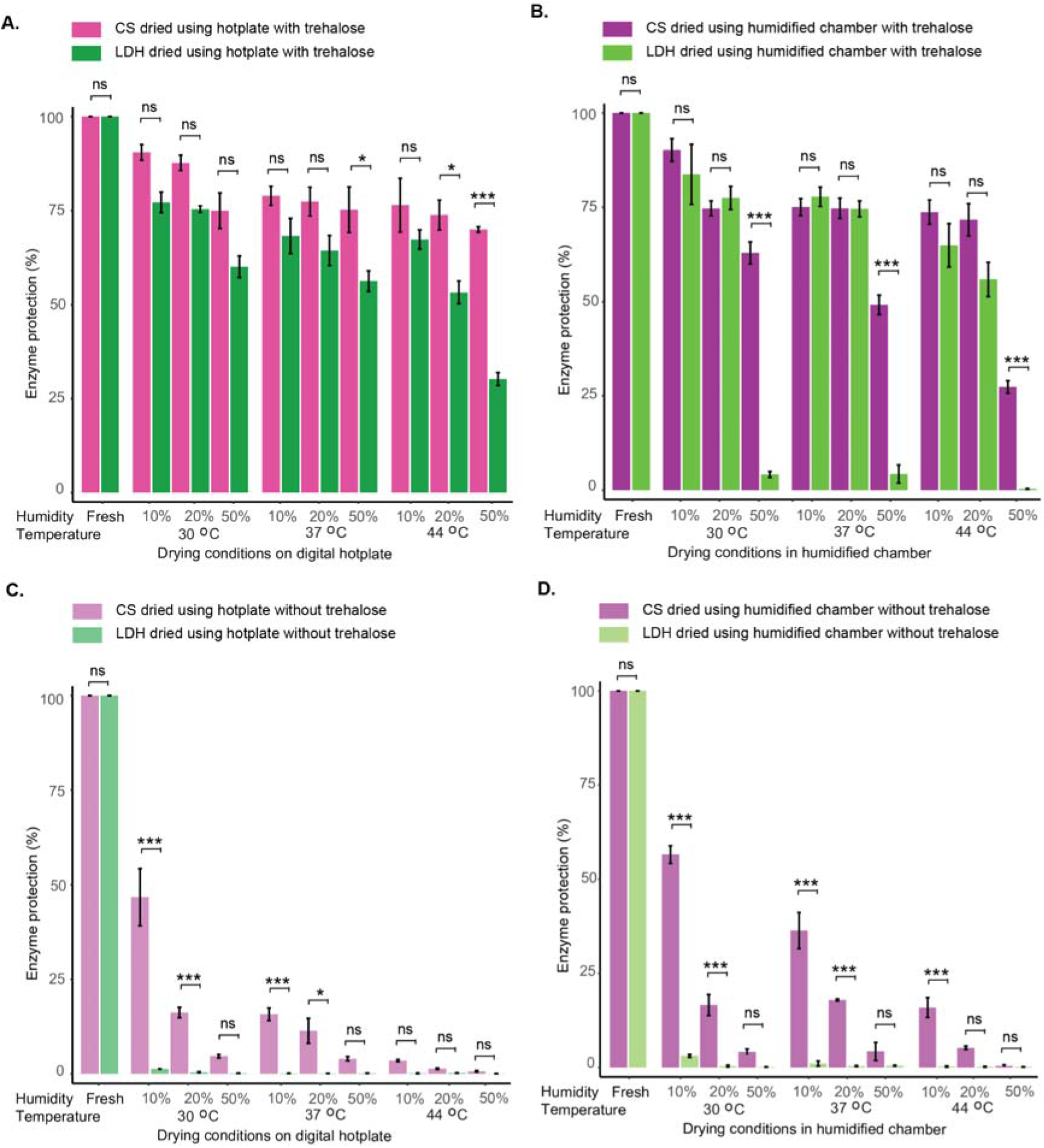
Effect of drying method and trehalose on CS and LDH activity. Citrate synthase (CS) and lactate dehydrogenase (LDH) activity in hotplate-dried samples with trehalose (A), humidified chamber-dried samples with trehalose (B), hotplate-dried samples without trehalose (C), and humidified chamber-dried samples without trehalose (D). Activity of CS and LDH was assessed using the citrate synthase and lactate dehydrogenase enzyme protection assays, respectively. Statistical comparisons were made only between treatments within the same drying method, as determined by one-way ANOVA followed by Tukey’s post hoc test (α = 0.05). Data represent mean ± SE from three independent replicates per condition.

**Supplementary Figure S8.**
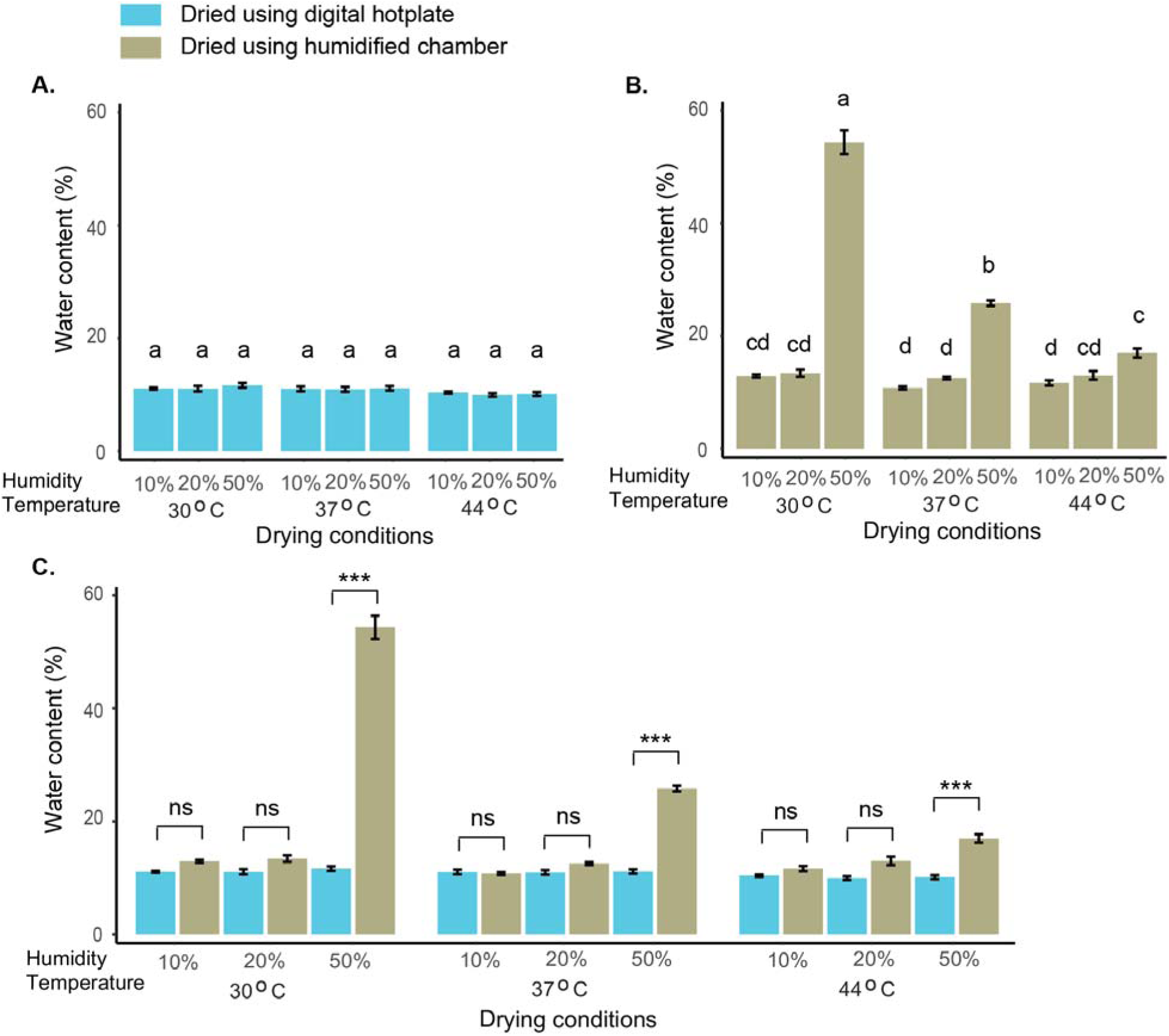
Effect of drying methods and conditions on water content in dried samples. Water content in hotplate-dried samples (A). Water content in humidified chamber-dried samples (B). Comparison of water content between hotplate and humidified chamber drying at each temperature and relative humidity (C). Statistical comparisons in (A) and (B) were determined using one-way ANOVA followed by Tukey’s post hoc test (α = 0.05); different letters indicate statistically significant differences among drying conditions within each panel. For (C), statistical comparisons were made only between treatments within the same drying condition using one-way ANOVA followed by Tukey’s post hoc test (α = 0.05). Data represent mean ± SE from three independent replicates per condition.

**Supplementary Figure S9.**
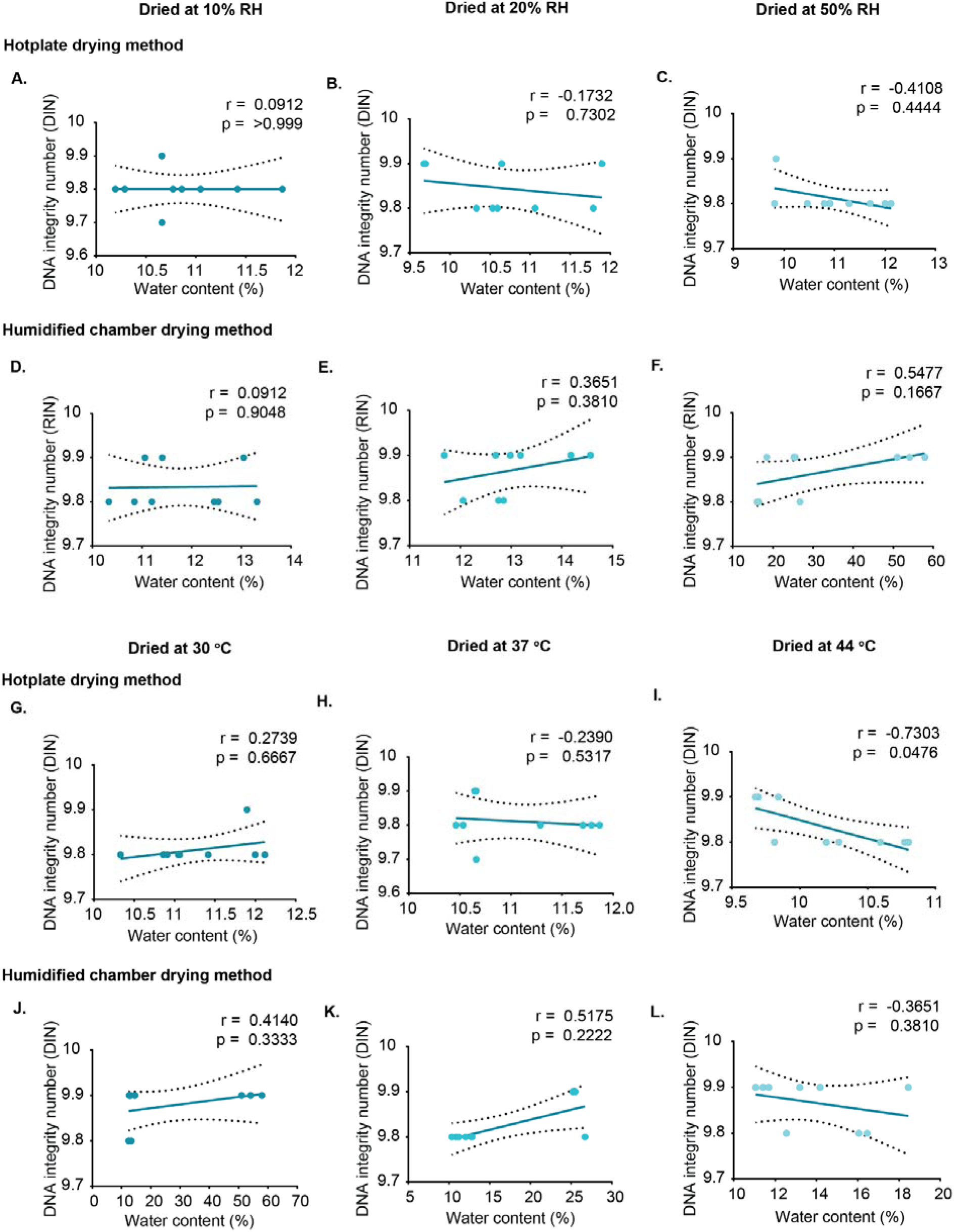
Correlation between residual water content and DNA integrity in trehalose-containing samples under constant humidity and constant temperature conditions using two drying methods. DNA integrity was assessed using the DNA Integrity Number (DIN). A-C show hotplate-dried samples at constant humidity, and D-F show humidified chamber-dried samples at constant humidity. G-I show hotplate-dried samples at constant temperature, and panels J-L show humidified chamber-dried samples at constant temperature. Correlation coefficients (r) and significance values (p) were calculated using Pearson correlation for normally distributed data and Spearman correlation for non-normally distributed data. Each data point represents an individual replicate. Dashed lines indicate 95% confidence interval (CI).

**Supplementary Figure S10.**
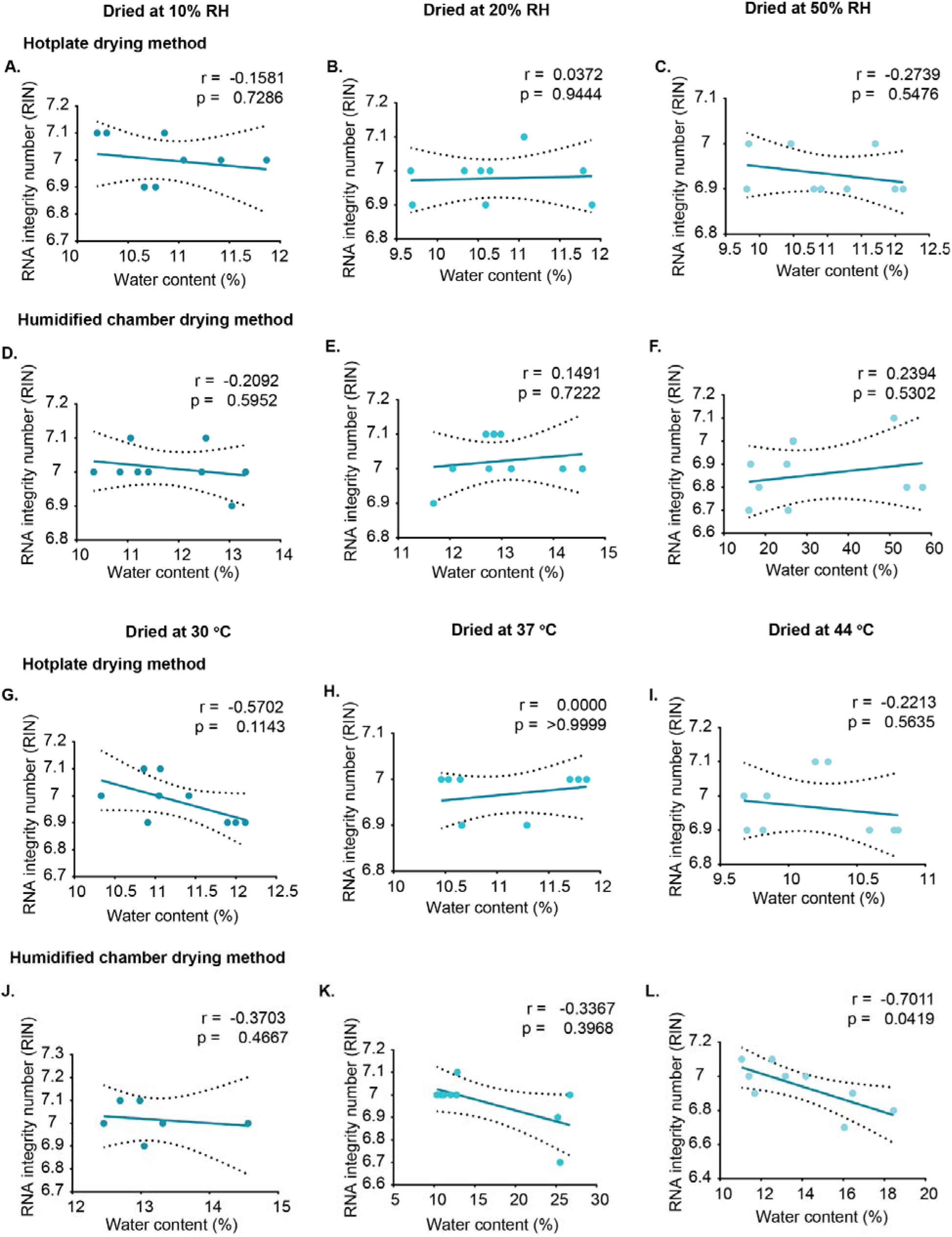
Correlation between residual water content and RNA integrity in trehalose-containing samples under constant humidity and constant temperature conditions using two drying methods. RNA integrity was assessed using the RNA Integrity Number (RIN). A-C show hotplate-dried samples at constant humidity, and D-F show humidified chamber-dried samples at constant humidity. G-I show hotplate-dried samples at constant temperature, and J-L show humidified chamber-dried samples at constant temperature. Correlation coefficients (r) and significance values (p) were calculated using Pearson correlation for normally distributed data and Spearman correlation for non-normally distributed data. Each data point represents an individual replicate. Dashed lines indicate 95% confidence interval (CI).

**Supplementary Figure S11.**
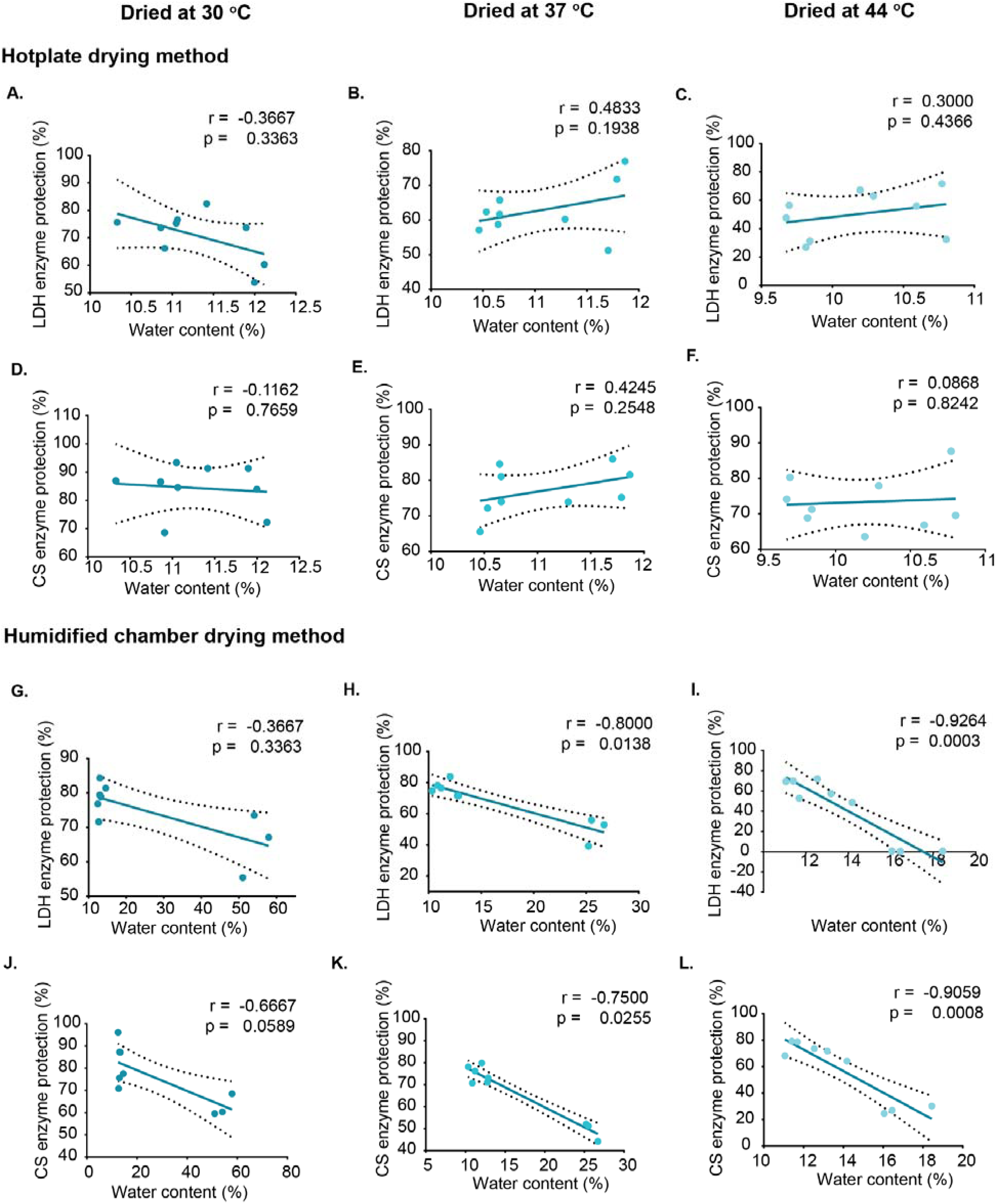
Correlation between residual water content and enzyme activity of LDH and CS under constant temperature and varying humidity conditions using two drying methods. Enzymatic activity of lactate dehydrogenase (LDH; A-C, G-I) and citrate synthase (CS; D-F, J-L) was assessed after drying under three different temperatures at each of the following constant relative humidity levels: 10%, 20%, and 50%. Drying was performed using either the hotplate method (A-F) or the humidified chamber method (G-L). Correlation coefficients (r) and significance values (p) were calculated using Pearson correlation for normally distributed data and Spearman correlation for non-normally distributed data. Each data point represents an individual replicate. Dashed lines indicate 95% confidence interval (CI).

**Supplementary Figure S12.**
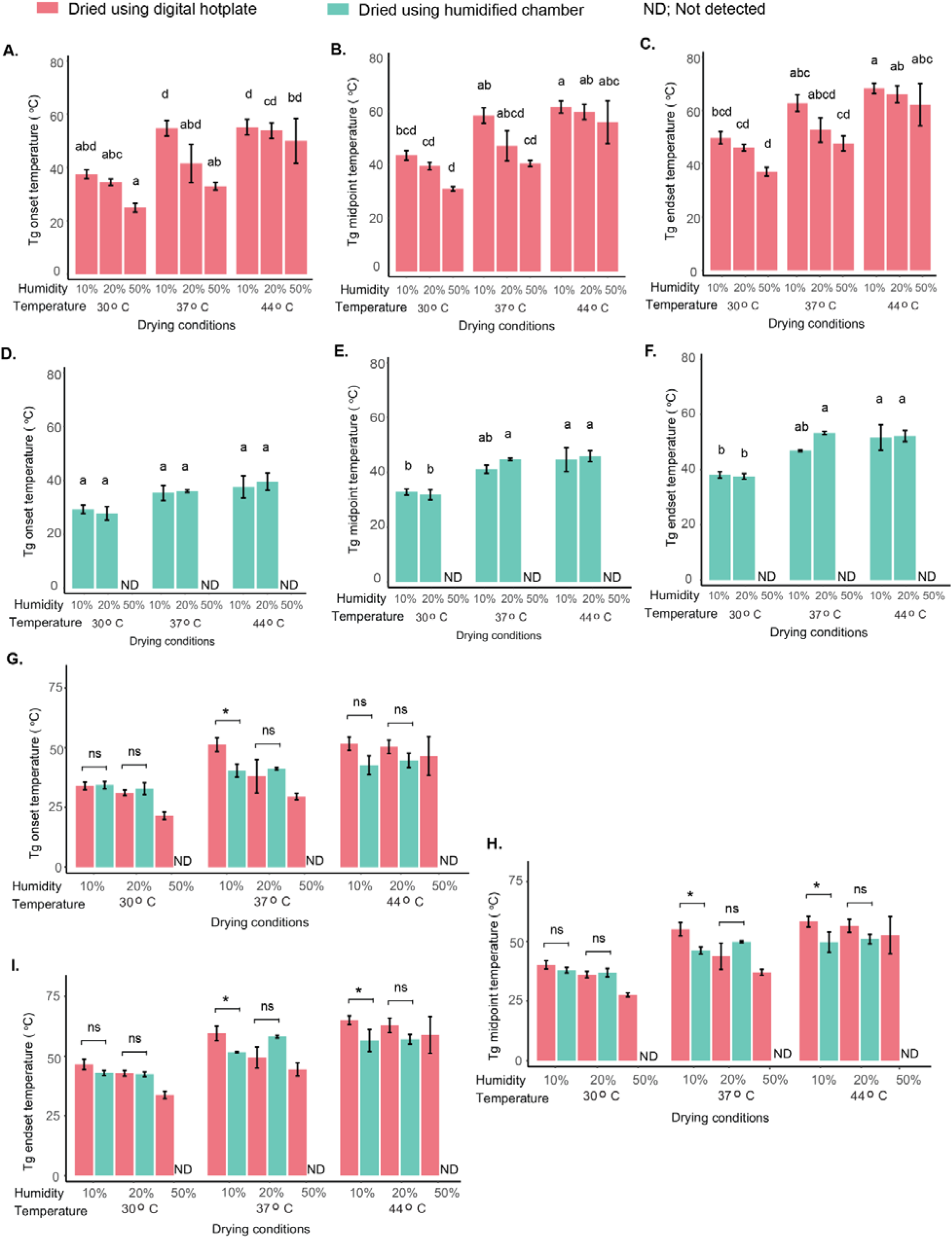
Glass transition temperature (Tg) of trehalose-containing samples under different drying methods and environmental conditions. Tg onset (A, D, G), midpoint (B, E, H), and endset (C, F, I) were measured for samples dried using the hotplate method (A-C) or the humidified chamber method (panels D-F) across a range of temperatures and relative humidities. G-I show direct comparisons of Tg values between hotplate and humidified chamber drying at each temperature and relative humidity. Statistical comparisons in A-F were determined using one-way ANOVA followed by Tukey’s post hoc test (α = 0.05); different letters indicate statistically significant differences among drying conditions within each panel. For G-I, statistical comparisons were made only between treatments within the same drying condition. Data represent mean ± SE from three independent replicates per condition.

**Supplementary Figure S13.**
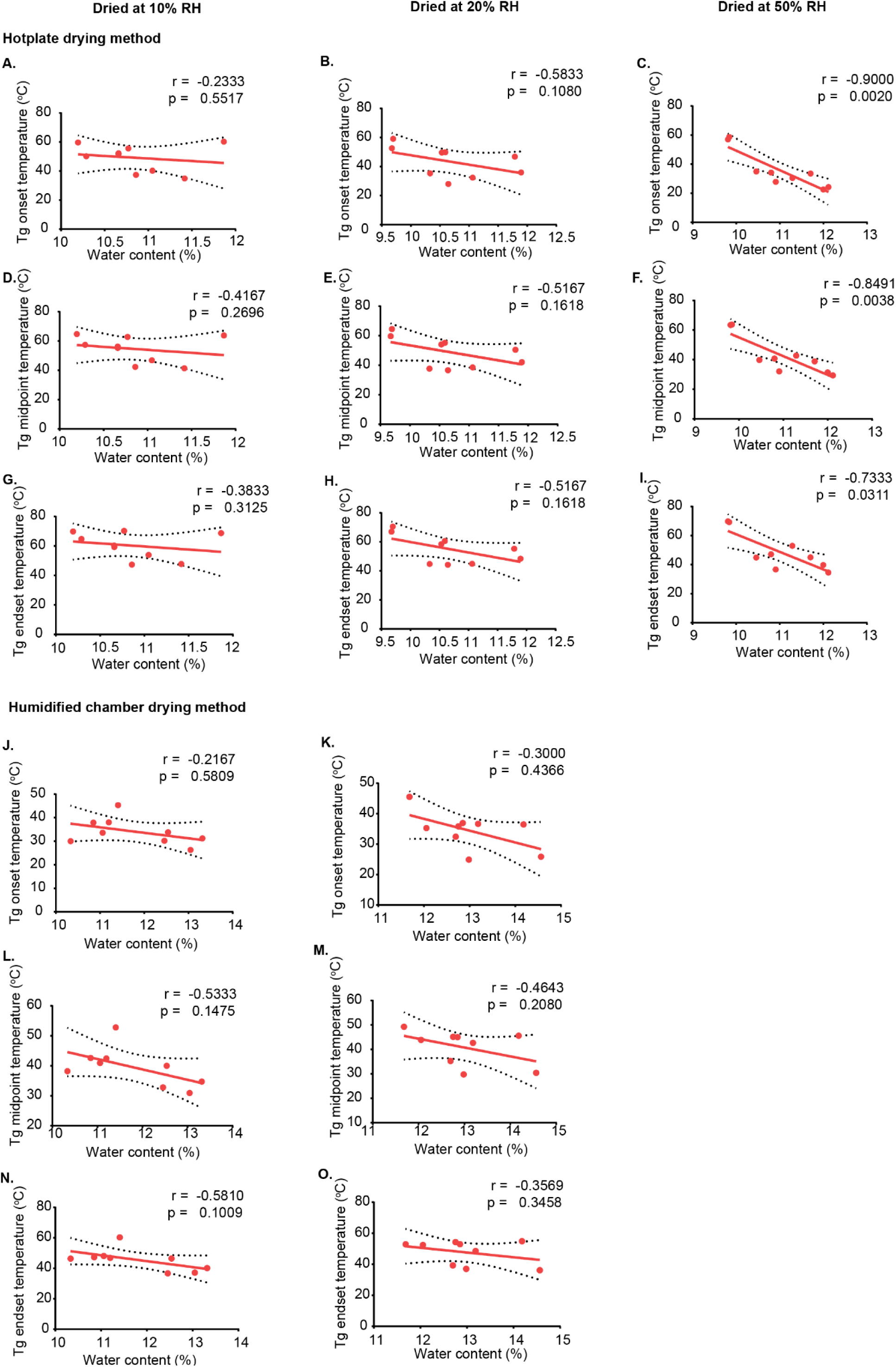
Correlation between residual water content and glass transition temperature (Tg) in trehalose-containing samples. Tg onset (A-C, J-K), midpoint (D-F, L-M), and endset (G-I, N-O) were assessed for samples dried under constant humidity with varying temperatures using either the hotplate method (A-I) or the humidified chamber method (J-O). Correlation coefficients (r) and significance values (p) were calculated using Pearson correlation for normally distributed data and Spearman correlation for non-normally distributed data. Each data point represents an individual replicate. Dashed lines indicate 95% confidence interval (CI).

**Supplementary Figure S14.**
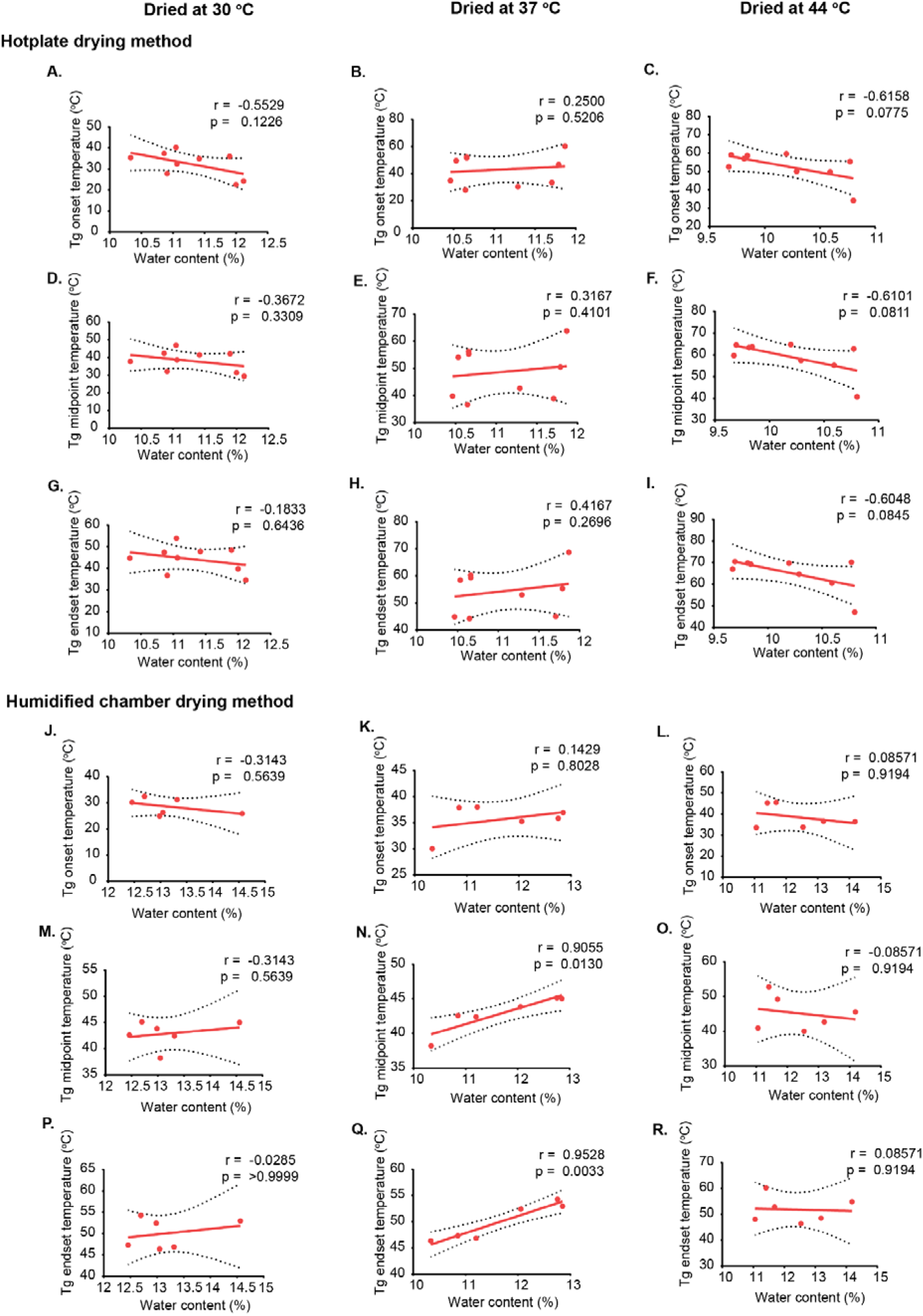
Correlation between residual water content and glass transition temperature (Tg) in trehalose-containing samples. Tg onset (A-C, J-L), midpoint (D-F, M-O), and endset (G-I, P-R) were assessed for samples dried under constant temperature with varying relative humidities using either the hotplate method (A-I) or the humidified chamber method (J-R). Correlation coefficients (r) and significance values (p) were calculated using Pearson correlation for normally distributed data and Spearman correlation for non-normally distributed data. Each data point represents an individual replicate. Dashed lines indicate 95% confidence interval (CI).

**Supplementary Figure S15.**
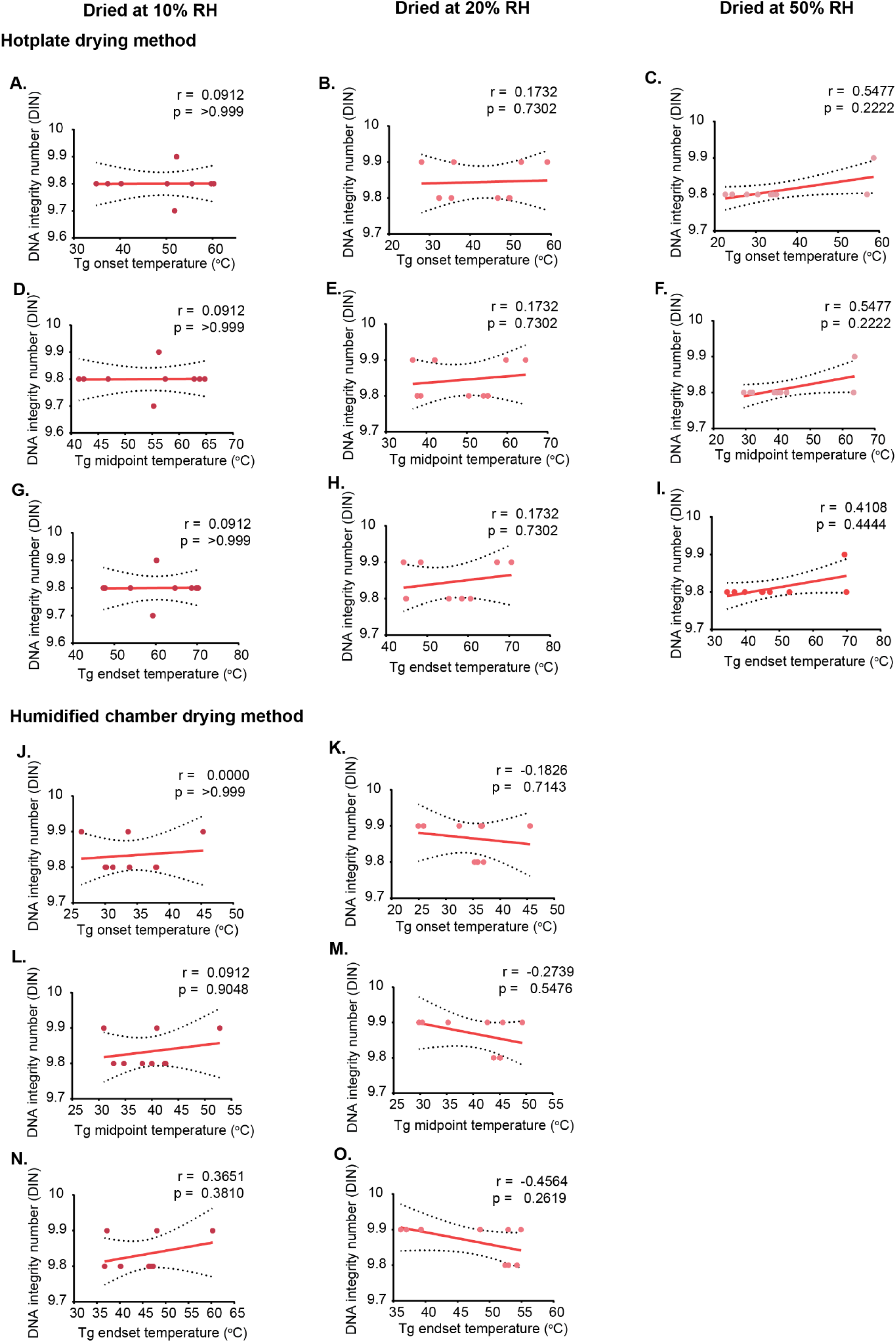
Correlation between glass transition temperature (Tg) and DNA integrity in samples dried using the hotplate and humidified chamber methods under constant humidity conditions. DNA integrity was assessed using the DNA Integrity Number (DIN). A-I show correlations between Tg values (onset, midpoint, and endset) and DNA integrity for hotplate-dried samples at 10%, 20%, and 50% relative humidity and humidified chamber-dried samples at 10% and 20% RH (50% RH not included for humidified chamber due to the absence of distinct Tg). Correlation coefficients (r) and significance values (p) were calculated using Pearson correlation for normally distributed data and Spearman correlation for non-normally distributed data. Each data point represents an individual replicate. Dashed lines indicate 95% confidence interval (CI).

**Supplementary Figure S16.**
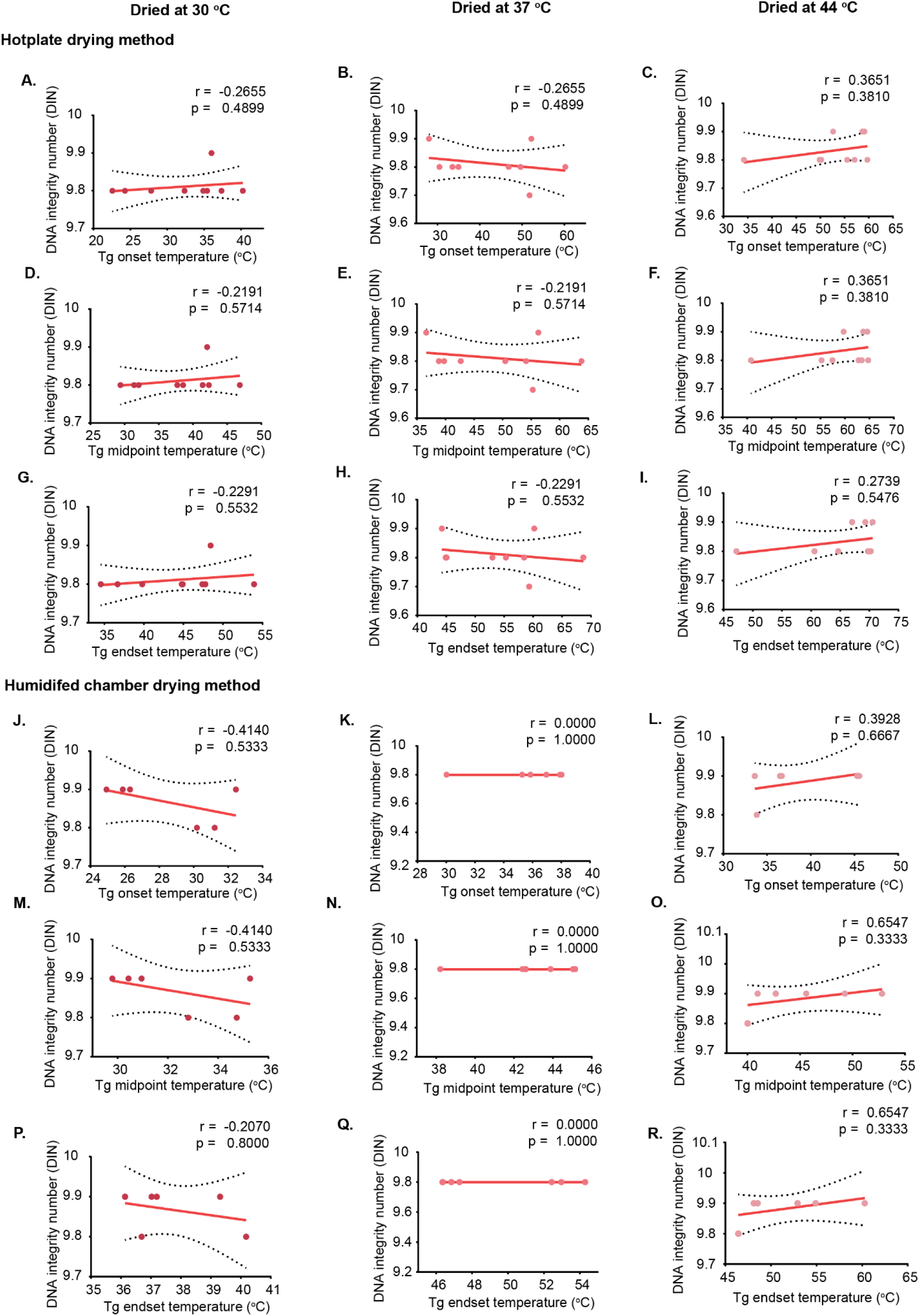
Correlation between glass transition temperature (Tg) and DNA integrity in samples dried using the hotplate and humidified chamber methods under constant temperature conditions. DNA integrity was assessed using the DNA Integrity Number (DIN). A-I show correlations between Tg values (onset, midpoint, and endset) and DNA integrity for hotplate-dried samples at 30 °C, 37 °C, and 44 °C, while J-R show correlations for humidified chamber-dried samples at the same temperatures. Correlation coefficients (r) and significance values (p) were calculated using Pearson correlation for normally distributed data and Spearman correlation for non-normally distributed data. Each data point represents an individual replicate. Dashed lines indicate 95% confidence interval (CI).

**Supplementary Figure S17.**
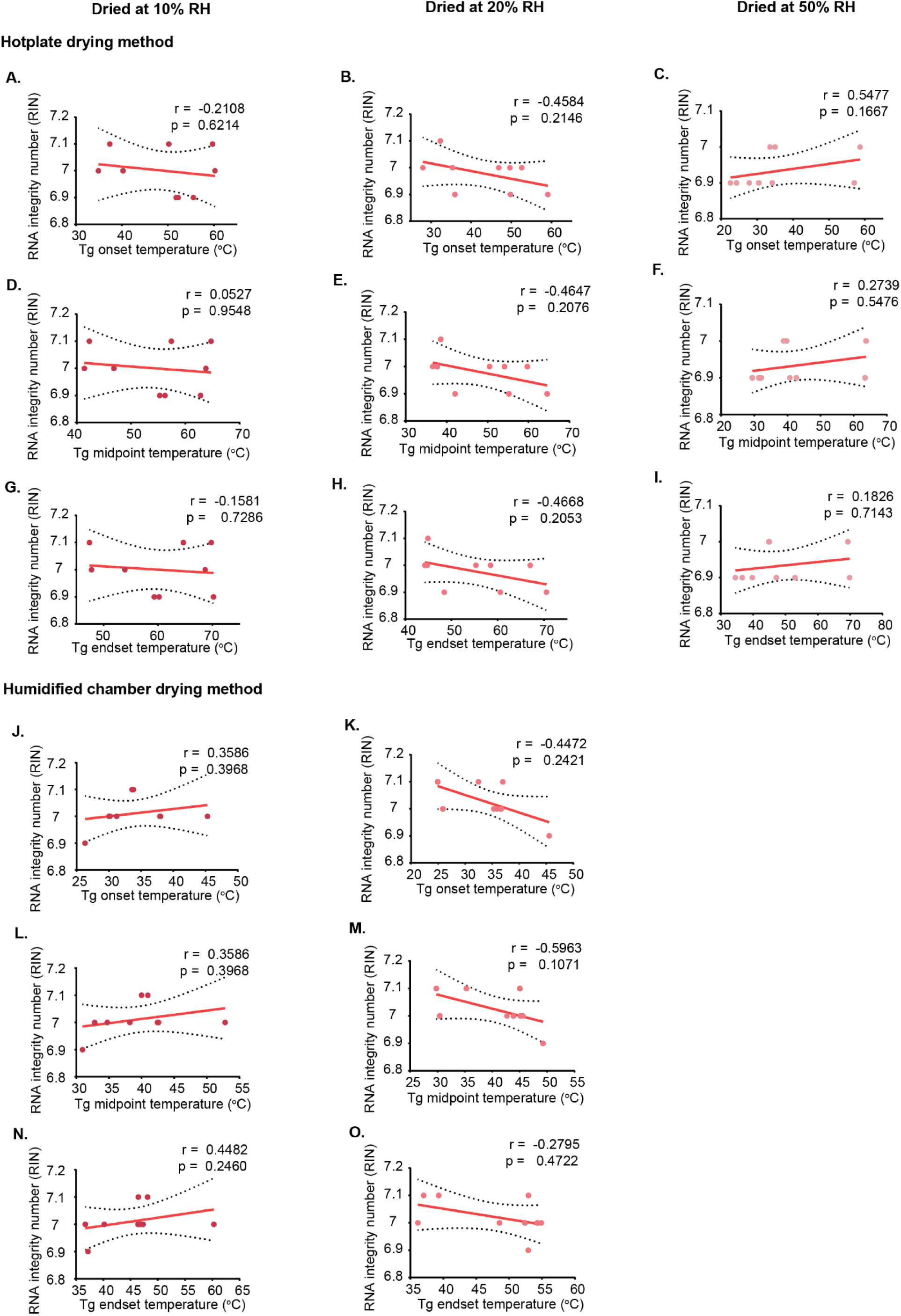
Correlation between glass transition temperature (Tg) and RNA integrity in samples dried using the hotplate and humidified chamber methods under constant humidity conditions. RNA integrity was assessed using the RNA Integrity Number (RIN). A-I show correlations between Tg values (onset, midpoint, and endset) and RNA integrity for hotplate-dried samples at 10%, 20%, and 50% relative humidity and humidified chamber-dried samples at 10% and 20% RH (50% RH not included for humidified chamber due to the absence of distinct Tg). Correlation coefficients (r) and significance values (p) were calculated using Pearson correlation for normally distributed data and Spearman correlation for non-normally distributed data. Each data point represents an individual replicate. Dashed lines indicate 95% confidence interval (CI).

**Supplementary Figure S18.**
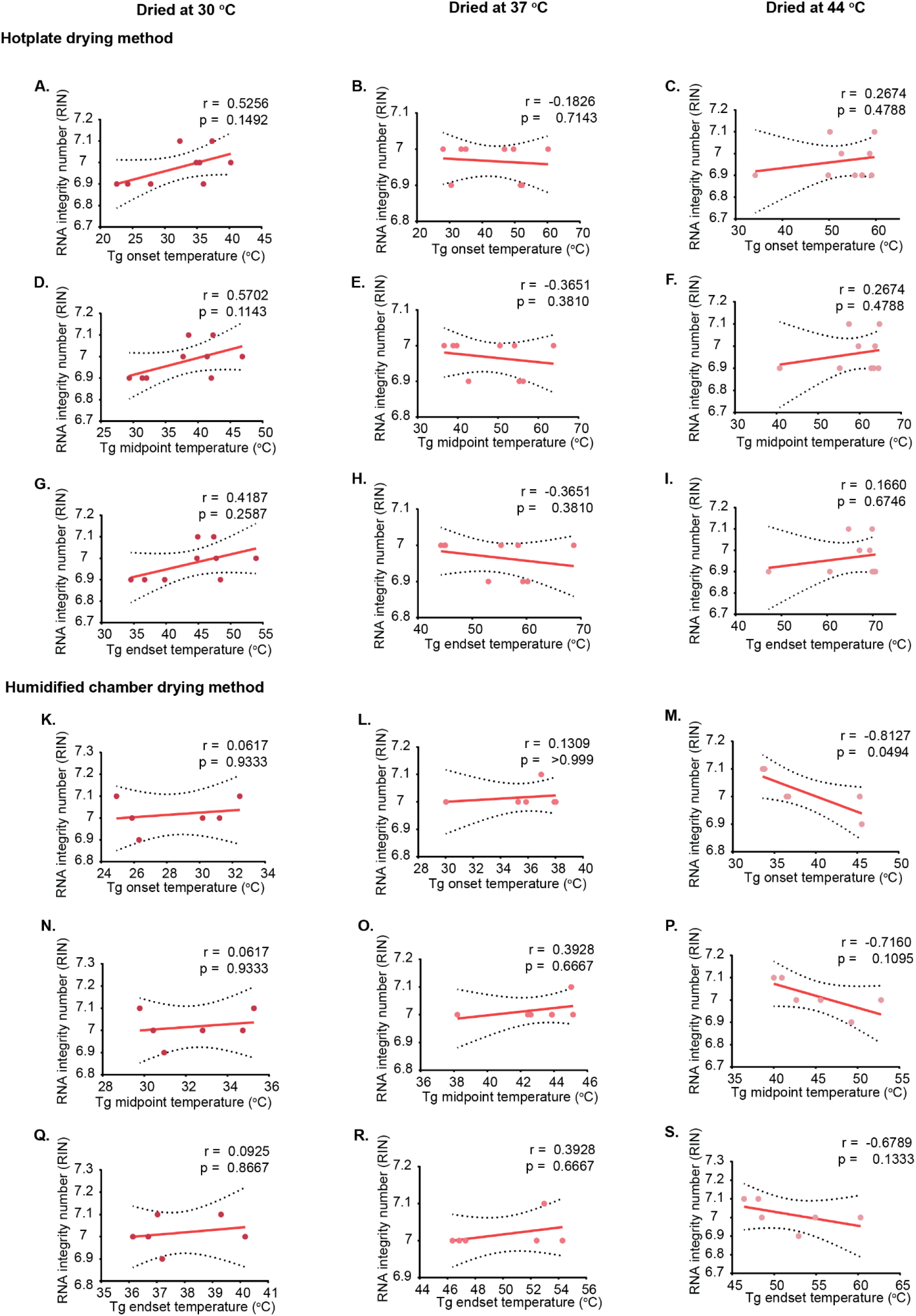
Correlation between glass transition temperature (Tg) and RNA integrity in samples dried using the hotplate and humidified chamber methods under constant temperature conditions. RNA integrity was assessed using the RNA Integrity Number (RIN). A-I show correlations between Tg values (onset, midpoint, and endset) and RNA integrity for hotplate-dried samples at 30 °C, 37 °C, and 44 °C, while J-R show correlations for humidified chamber-dried samples at the same temperatures. Correlation coefficients (r) and significance values (p) were calculated using Pearson correlation for normally distributed data and Spearman correlation for non-normally distributed data. Each data point represents an individual replicate. Dashed lines indicate 95% confidence interval (CI).

**Supplementary Figure S19.**
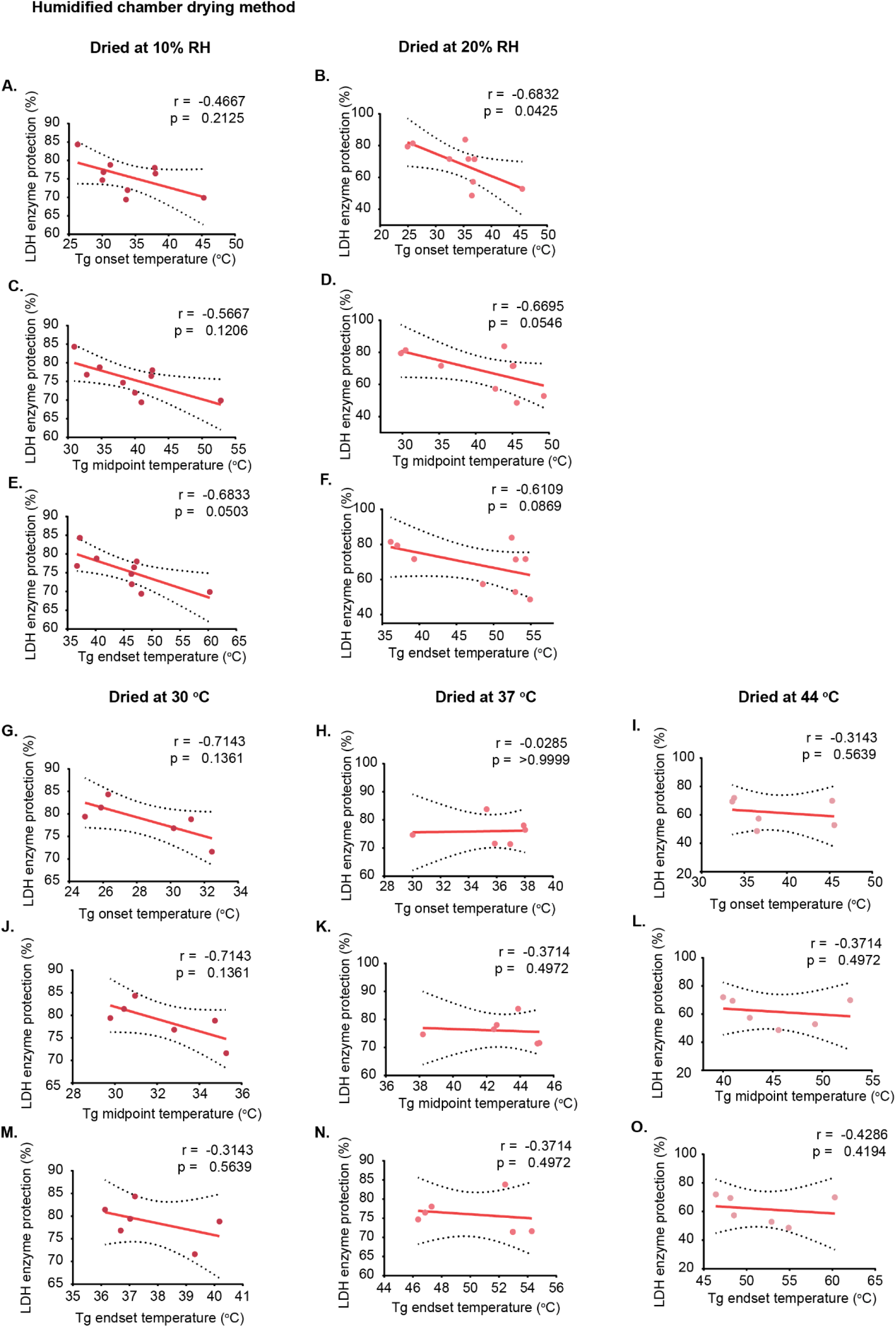
Correlation between glass transition temperature (Tg) and LDH activity in samples dried using the humidified chamber method under constant humidity and temperature conditions. A-F show correlations between Tg values (onset, midpoint, and endset) and lactate dehydrogenase (LDH) activity under constant humidity conditions (10% and 20% RH; 50% RH not included due to absence of measurable Tg), while panels G-O show correlations under constant temperature conditions (30 °C, 37 °C, and 44 °C). Correlation coefficients (r) and significance values (p) were calculated using Pearson correlation for normally distributed data and Spearman correlation for non-normally distributed data. Each data point represents an individual replicate. Dashed lines indicate 95% confidence interval (CI).

**Supplementary Figure S20.**
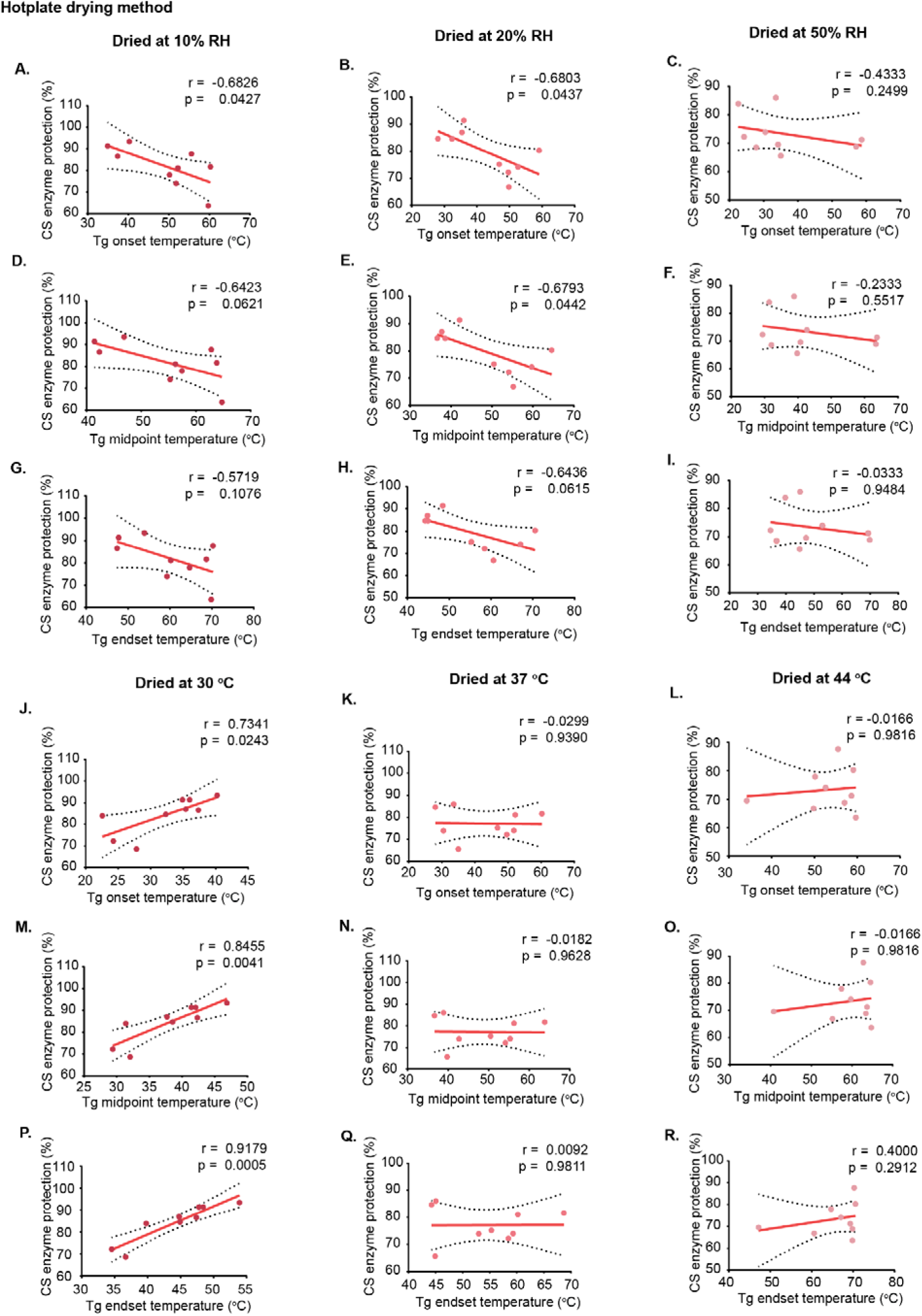
Correlation between glass transition temperature (Tg) and CS activity in samples dried using the hotplate method under constant humidity and temperature conditions. Citrate synthase (CS) activity was assessed in samples dried using the hotplate method. A-I show correlations between Tg values (onset, midpoint, and endset) and CS activity under constant humidity conditions (10%, 20%, and 50% RH), while J-R show correlations under constant temperature conditions (30 °C, 37 °C, and 44 °C). Correlation coefficients (r) and significance values (p) were calculated using Pearson correlation for normally distributed data and Spearman correlation for non-normally distributed data. Each data point represents an individual replicate. Dashed lines indicate 95% confidence interval (CI).

**Supplementary Figure S21.**
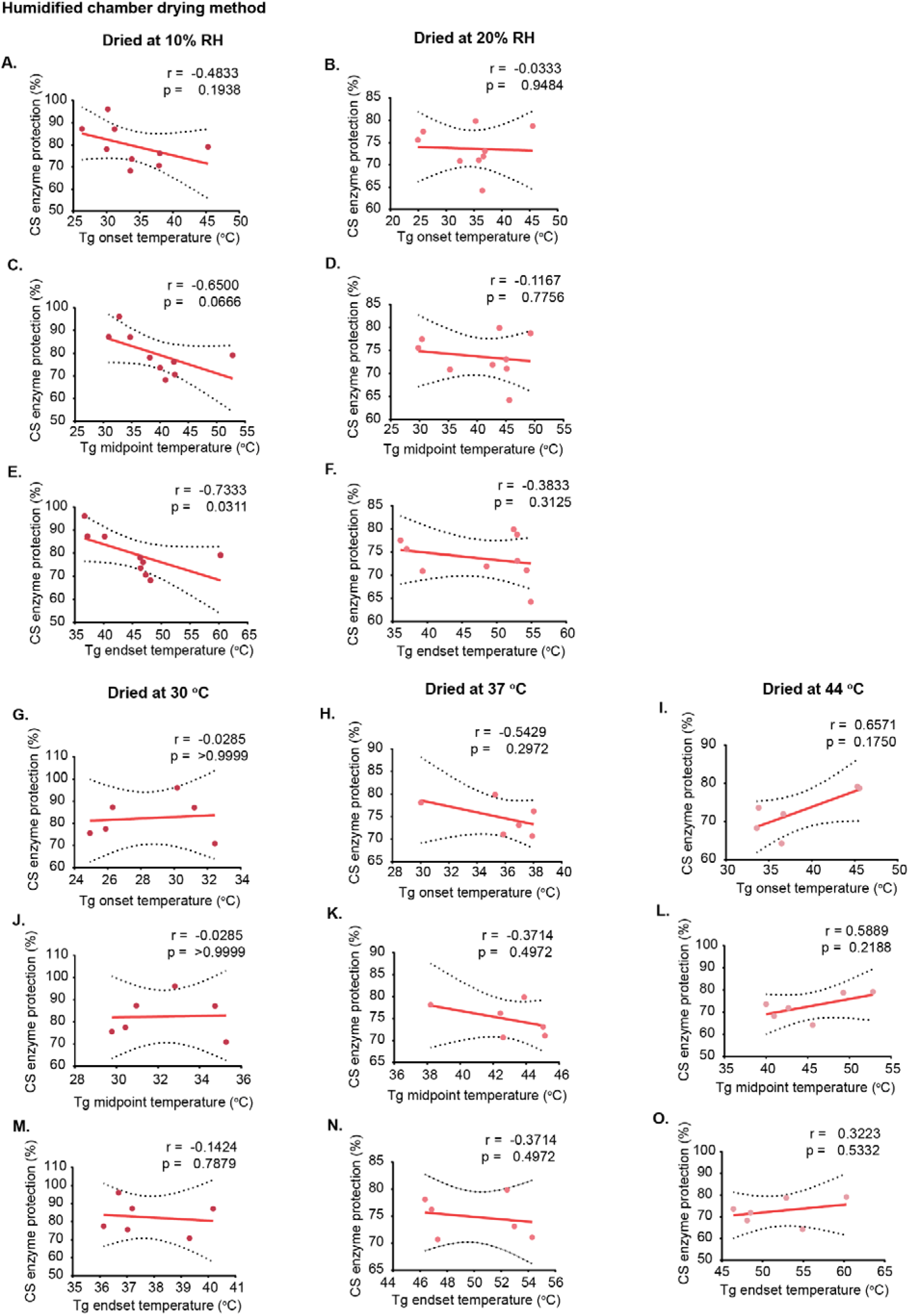
Correlation between glass transition temperature (Tg) and CS activity in samples dried using the humidified chamber method under constant humidity and temperature conditions. A-F show correlations between Tg values (onset, midpoint, and endset) and CS activity under constant humidity conditions (10% and 20% RH; 50% RH not included due to absence of measurable Tg), while G-O show correlations under constant temperature conditions (30 °C, 37 °C, and 44 °C). Correlation coefficients (r) and significance values (p) were calculated using Pearson correlation for normally distributed data and Spearman correlation for non-normally distributed data. Each data point represents an individual replicate. Dashed lines indicate 95% confidence interval (CI).

**Supplementary Figure S22.**
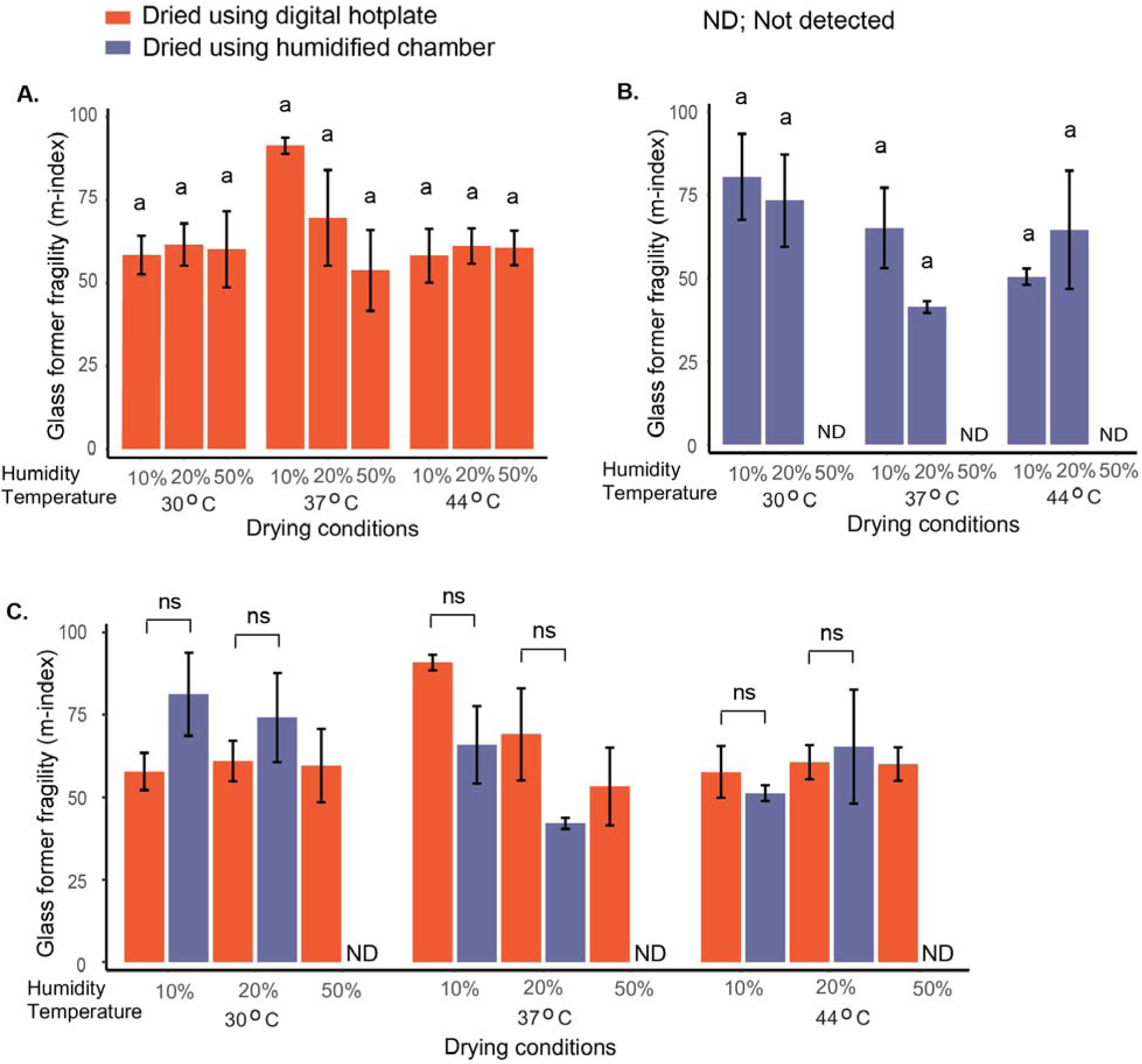
Effect of drying methods and conditions on glass former fragility (m-index) in dried samples. Glass former fragility (m-index) in hotplate-dried samples (A) and humidified chamber-dried samples (B). Comparison of fragility between hotplate and humidified chamber drying at each temperature and relative humidity (C). Statistical comparisons in panels A and B were performed using one-way ANOVA followed by Tukey’s post hoc test (α = 0.05); different letters indicate statistically significant differences among drying conditions within each panel. For panel C, comparisons were made only between treatments within the same drying condition using one-way ANOVA followed by Tukey’s post hoc test (α = 0.05). Data represent mean ± SE from three independent replicates per condition.

**Supplementary Figure S23.**
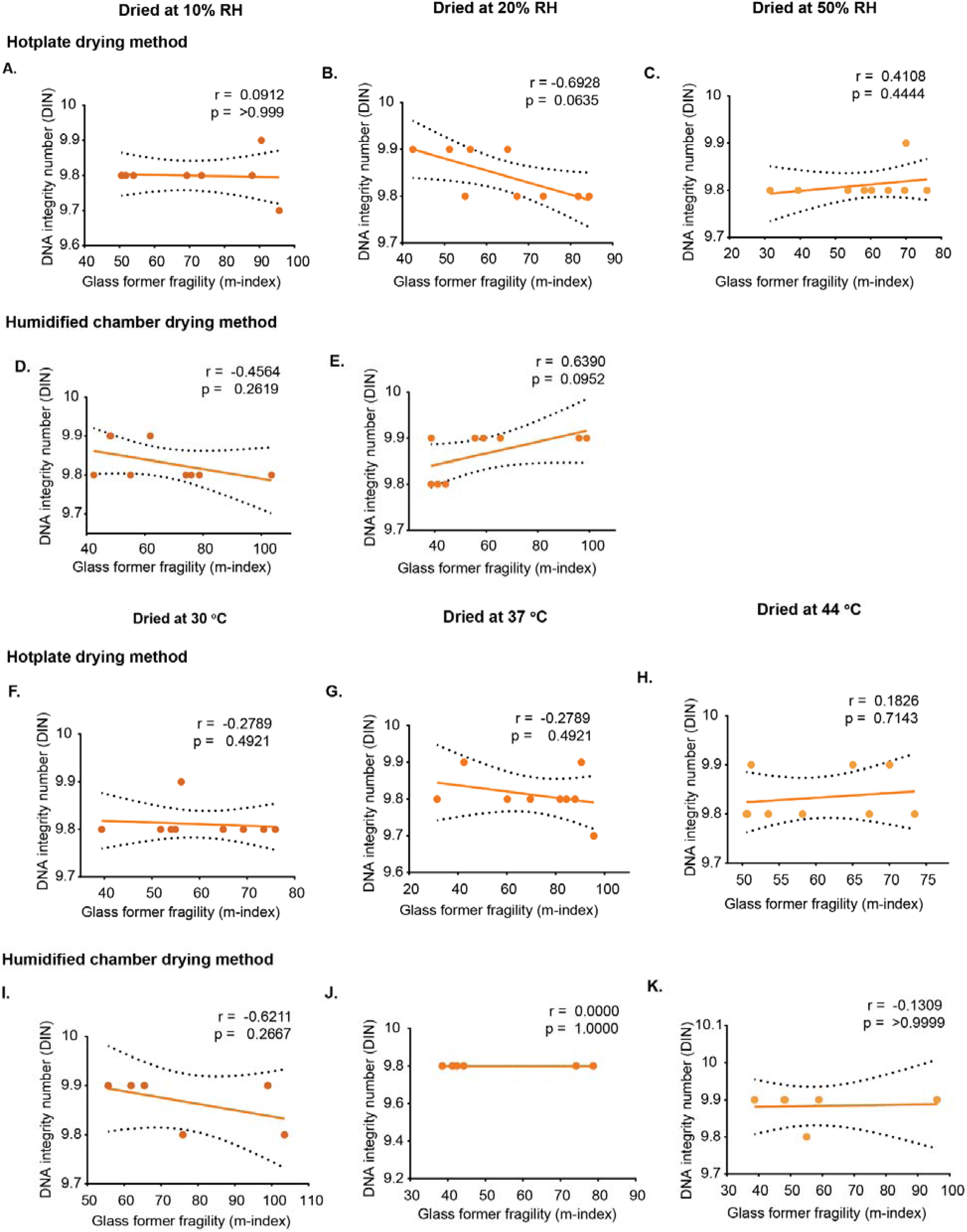
Correlation between glass former fragility (m-index) and DNA integrity in samples dried using the hotplate and humidified chamber methods under constant humidity and constant temperature conditions. DNA integrity was assessed using the DNA Integrity Number (DIN). A-C show correlations for hotplate-dried samples at constant humidity (10%, 20%, and 50% RH), while D,E show correlations for humidified chamber-dried samples at constant humidity (10% and 20% RH; 50% RH not included as no fragility value was obtained). F-I show correlations for hotplate-dried samples at constant temperature (30 °C, 37 °C, and 44 °C), and I-K show correlations for humidified chamber-dried samples at constant temperature (30 °C, 37 °C, and 44 °C). Correlation coefficients (r) and significance values (p) were calculated using Pearson correlation for normally distributed data and Spearman correlation for non-normally distributed data. Each data point represents an individual replicate. Dashed lines indicate 95% confidence interval (CI).

**Supplementary Figure S24.**
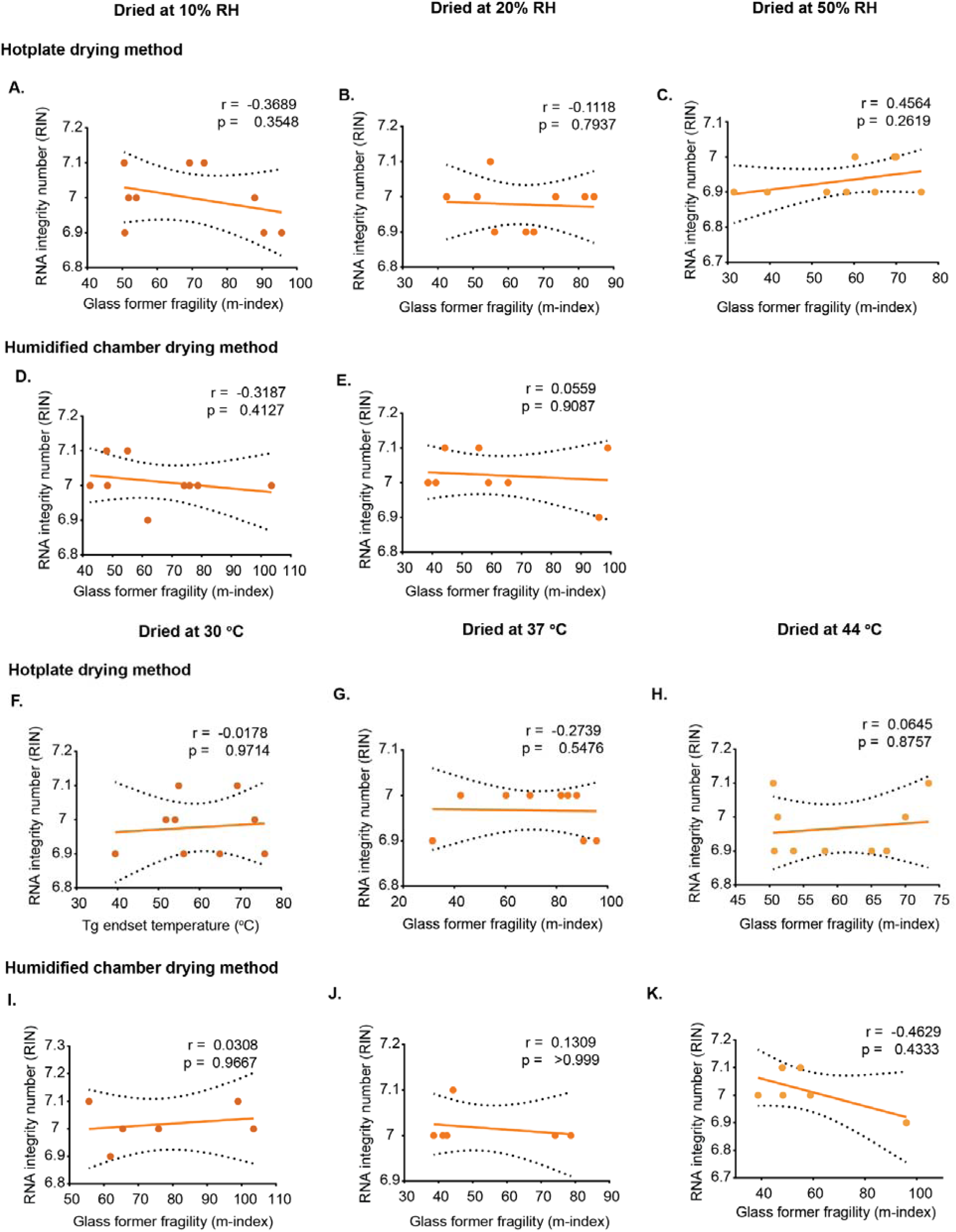
Correlation between glass former fragility (m-index) and RNA integrity in samples dried using the hotplate and humidified chamber methods under constant humidity and constant temperature conditions. RNA integrity was assessed using the RNA Integrity Number (RIN). A-C show correlations for hotplate-dried samples at constant humidity (10%, 20%, and 50% RH), while D,E show correlations for humidified chamber-dried samples at constant humidity (10% and 20% RH; 50% RH not included as no fragility value was obtained). F-I show correlations for hotplate-dried samples at constant temperature (30 °C, 37 °C, and 44 °C), and I-K show correlations for humidified chamber-dried samples at constant temperature (30 °C, 37 °C, and 44 °C). Correlation coefficients (r) and significance values (p) were calculated using Pearson correlation for normally distributed data and Spearman correlation for non-normally distributed data. Each data point represents an individual replicate. Dashed lines indicate 95% confidence interval (CI).

**Supplementary Figure S25.**
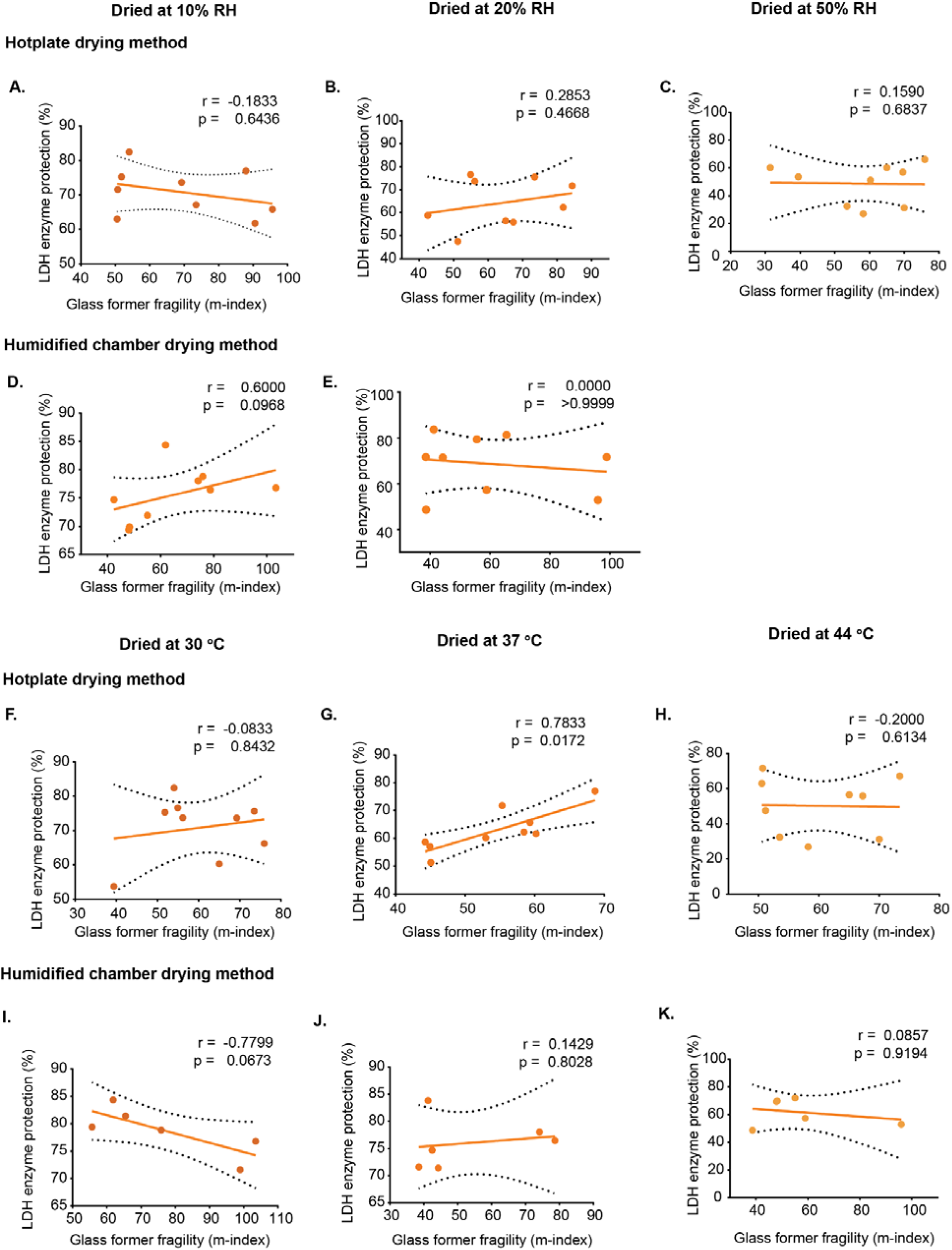
Correlation between glass former fragility (m-index) and LDH activity in samples dried using the hotplate and humidified chamber methods under constant humidity and constant temperature conditions. Lactate dehydrogenase (LDH) activity was expressed as the percentage of activity retained relative to the undried control. A-C show correlations for hotplate-dried samples at constant humidity (10%, 20%, and 50% RH), while D,E show correlations for humidified chamber-dried samples at constant humidity (10% and 20% RH; 50% RH not included as no fragility value was obtained). F-H show correlations for hotplate-dried samples at constant temperature (30 °C, 37 °C, and 44 °C), and I-K show correlations for humidified chamber-dried samples at constant temperature (30 °C, 37 °C, and 44 °C). Correlation coefficients (r) and significance values (p) were calculated using Pearson correlation for normally distributed data and Spearman correlation for non-normally distributed data. Each data point represents an individual replicate. Dashed lines indicate 95% confidence interval (CI).

**Supplementary Figure S26.**
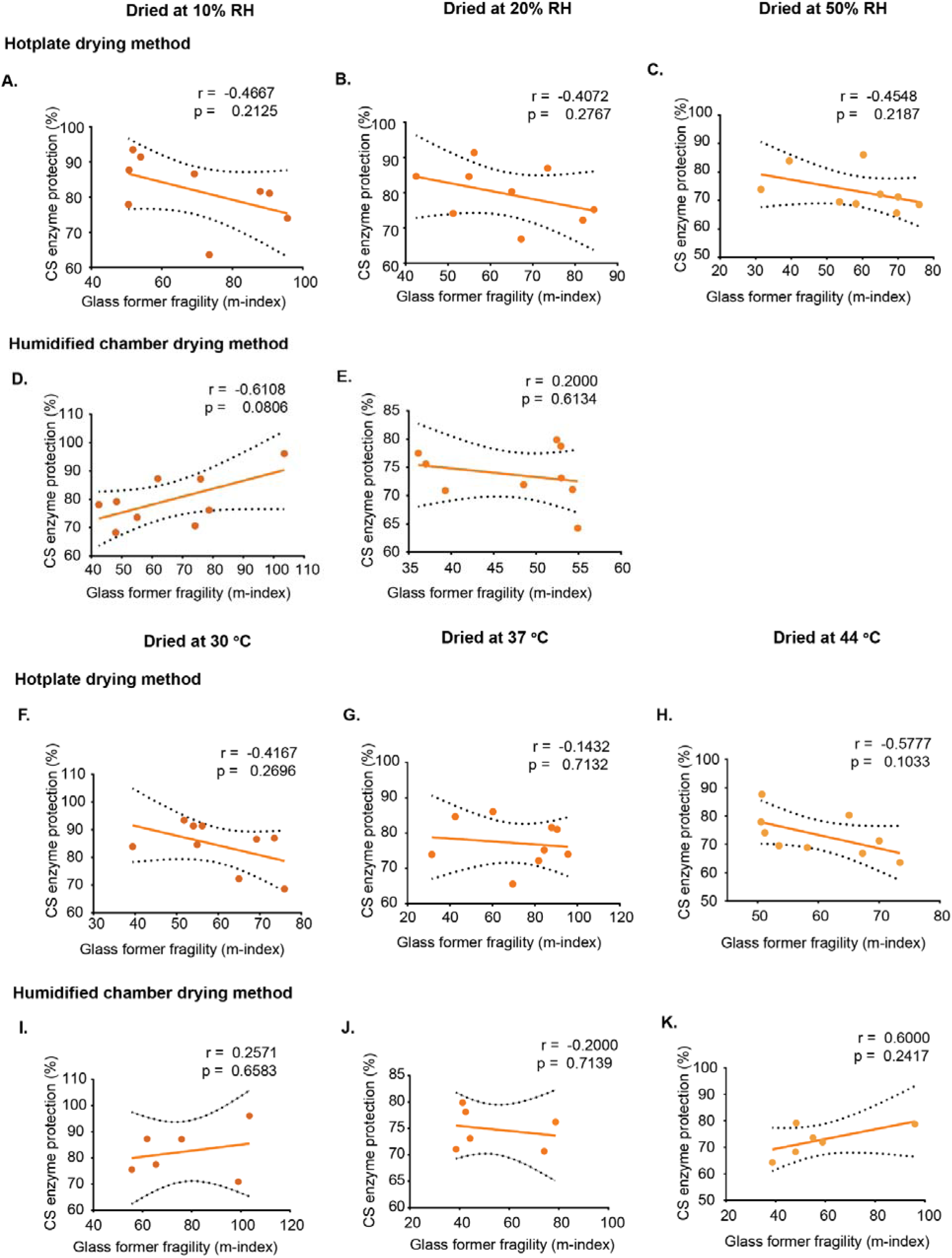
Correlation between glass former fragility (m-index) and CS activity in samples dried using the hotplate and humidified chamber methods under constant humidity and constant temperature conditions. A-C show correlations for hotplate-dried samples at constant humidity (10%, 20%, and 50% RH), while D,E show correlations for humidified chamber-dried samples at constant humidity (10% and 20% RH; 50% RH not included as no fragility value was obtained). F-H show correlations for hotplate-dried samples at constant temperature (30 °C, 37 °C, and 44 °C), and I-K show correlations for humidified chamber-dried samples at constant temperature (30 °C, 37 °C, and 44 °C). Correlation coefficients (r) and significance values (p) were calculated using Pearson correlation for normally distributed data and Spearman correlation for non-normally distributed data. Each data point represents an individual replicate. Dashed lines indicate 95% confidence interval (CI).

**Supplementary Figure 27.**
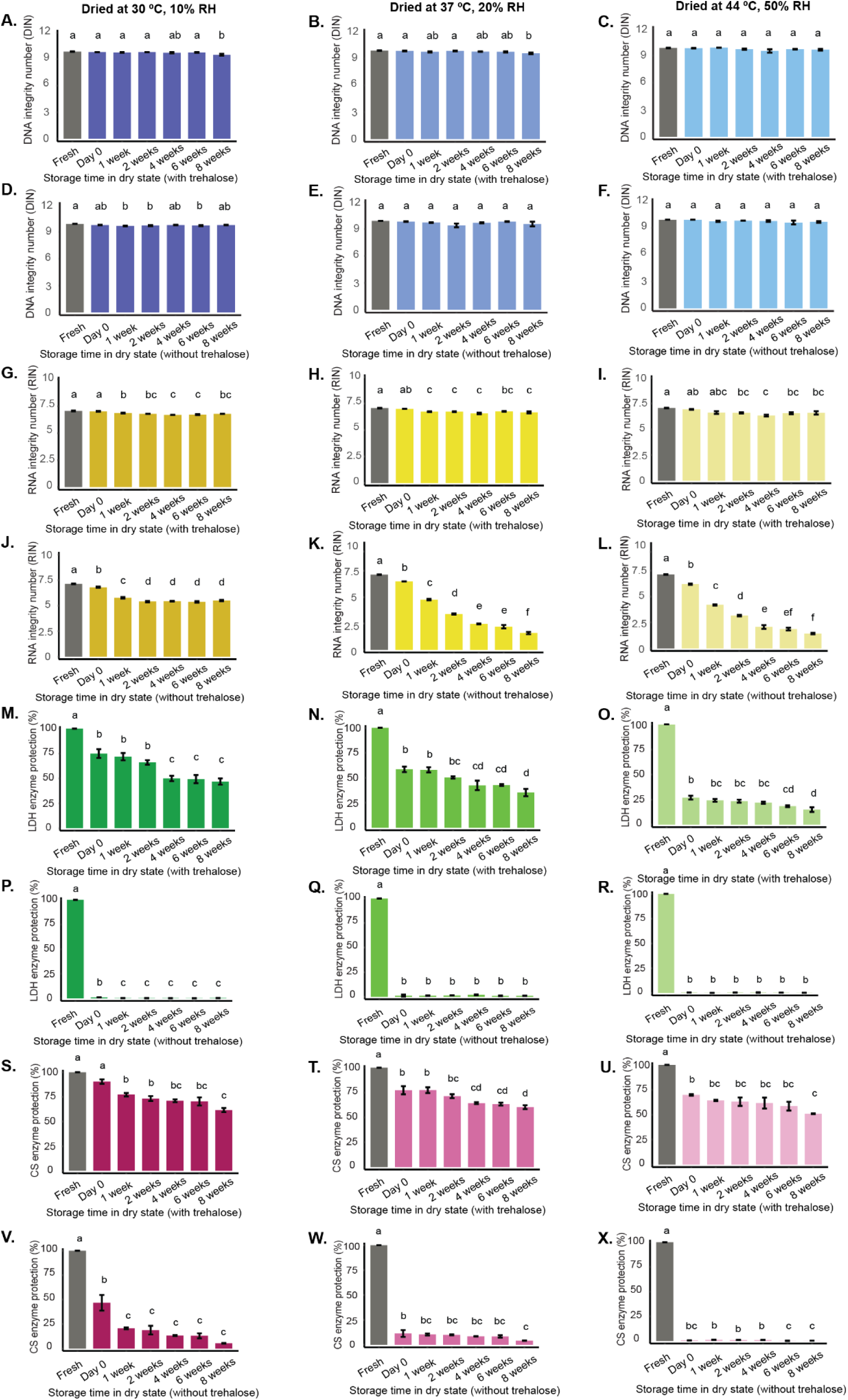
Molecular stability during storage under different drying conditions with and without trehalose. Samples of DNA (A–F), RNA (G–L), lactate dehydrogenase (LDH; M-R), and citrate synthase (CS; S-X) were dried using the hotplate method under three conditions: 30 °C at 10% RH (A, D, G, J, M, P, S, V), 37 °C at 20% RH (B, E, H, K, N, Q, T, W), and 44 °C at 50% RH (C, F, I, L, O, R, U, X). For each molecule, samples with trehalose are shown in the first three panels of each group (A-C, G-I, M-O, S-U), and samples without trehalose are shown in the next three panels (D-F, J-L, P-R, V-X). Samples were stored at room temperature in a 10% RH LiCl jar, and molecular stability was evaluated at Day 0, Week 1, Week 2, Week 4, and Week 8. Nucleic acid integrity and enzyme activity were assessed at each time point. Data represent mean ± SE from three independent experiments. Statistical comparisons were performed using one-way ANOVA followed by Tukey’s post hoc test (α = 0.05); different letters indicate statistically significant differences among drying conditions within each panel.

**Supplementary Figure 28.**
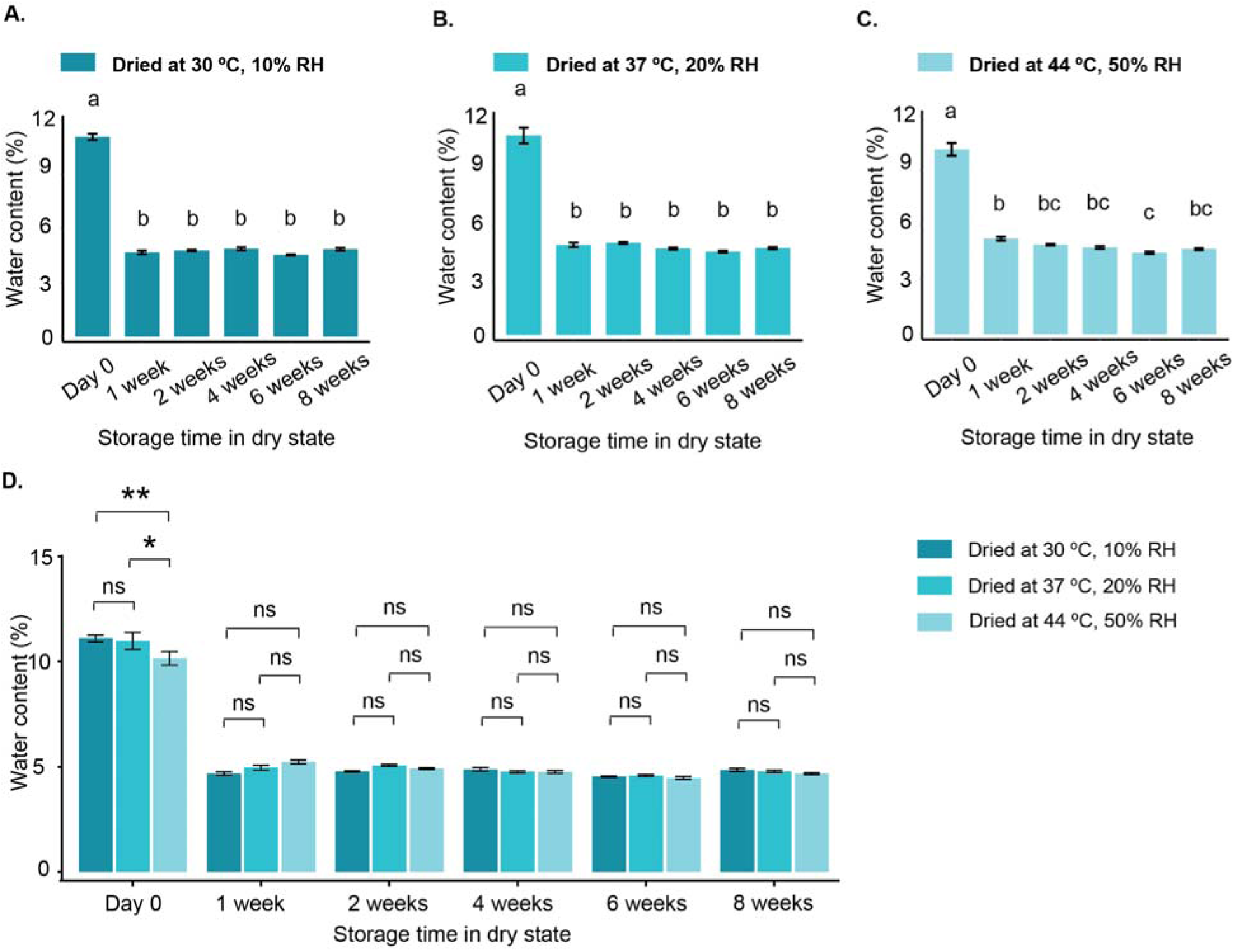
Water content of dried samples during storage under different drying conditions. Water content of samples dried under three conditions (A-C): 30 °C, 10% humidity (A); 37 °C, 20% humidity (B); and 44 °C, 50% humidity (C). Direct comparison of water content across the three drying conditions is shown in (D). Data are presented as mean ± SE. In panels A-C, different letters indicate statistical significance determined by one-way ANOVA followed by Tukey’s test. In panel D, statistical analysis was performed using two-way ANOVA with multiple comparisons.

**Supplementary Figure 29.**
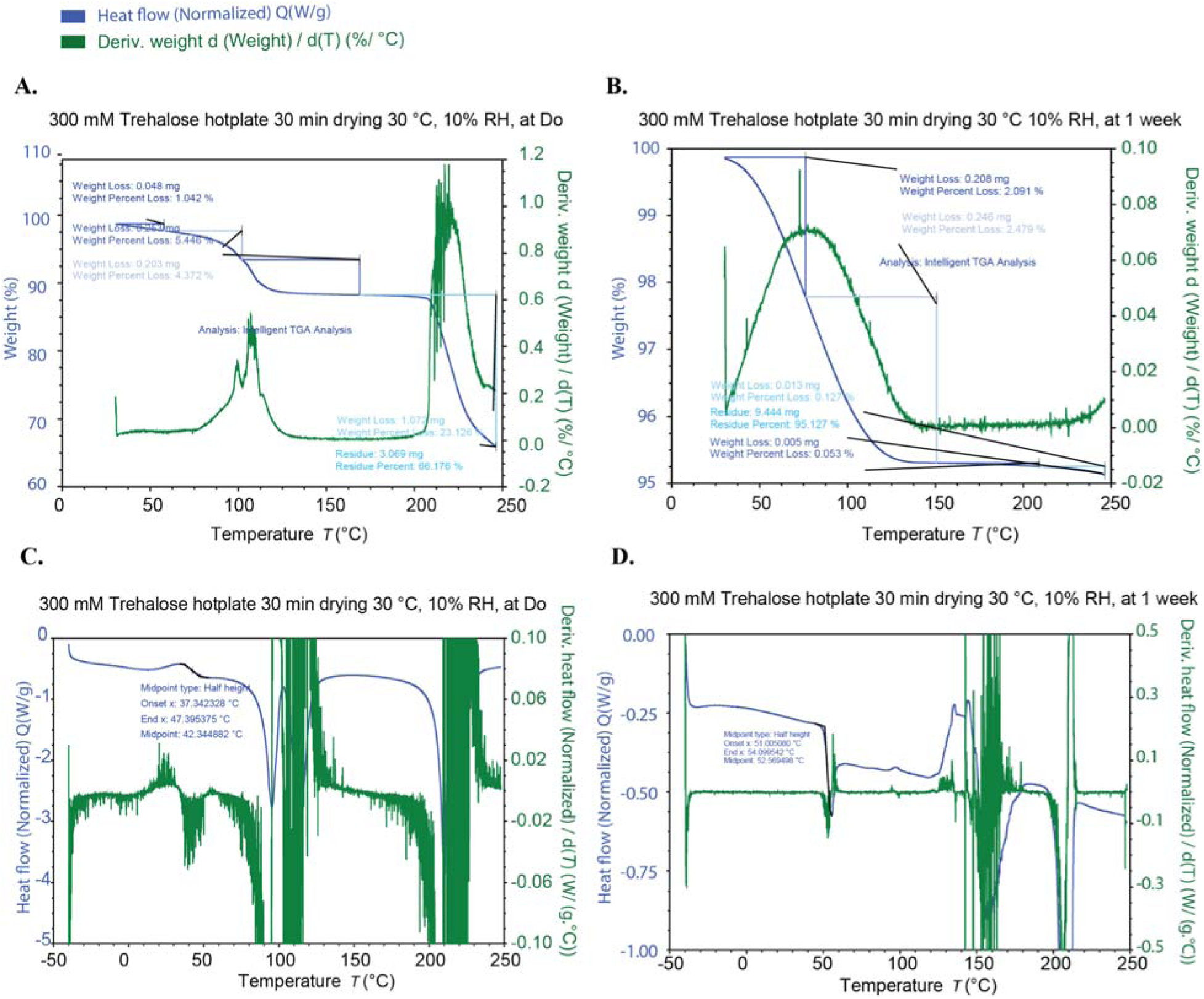
Thermogravimetric analysis (TGA) and differential scanning calorimetry (DSC) thermographs of samples dried at 30 °C, 10% humidity. A and B show TGA thermographs immediately after drying (A) and after 1 week of storage (B). C and D show DSC thermographs immediately after drying (C) and after 1 week of storage (D).

**Supplementary Figure 30.**
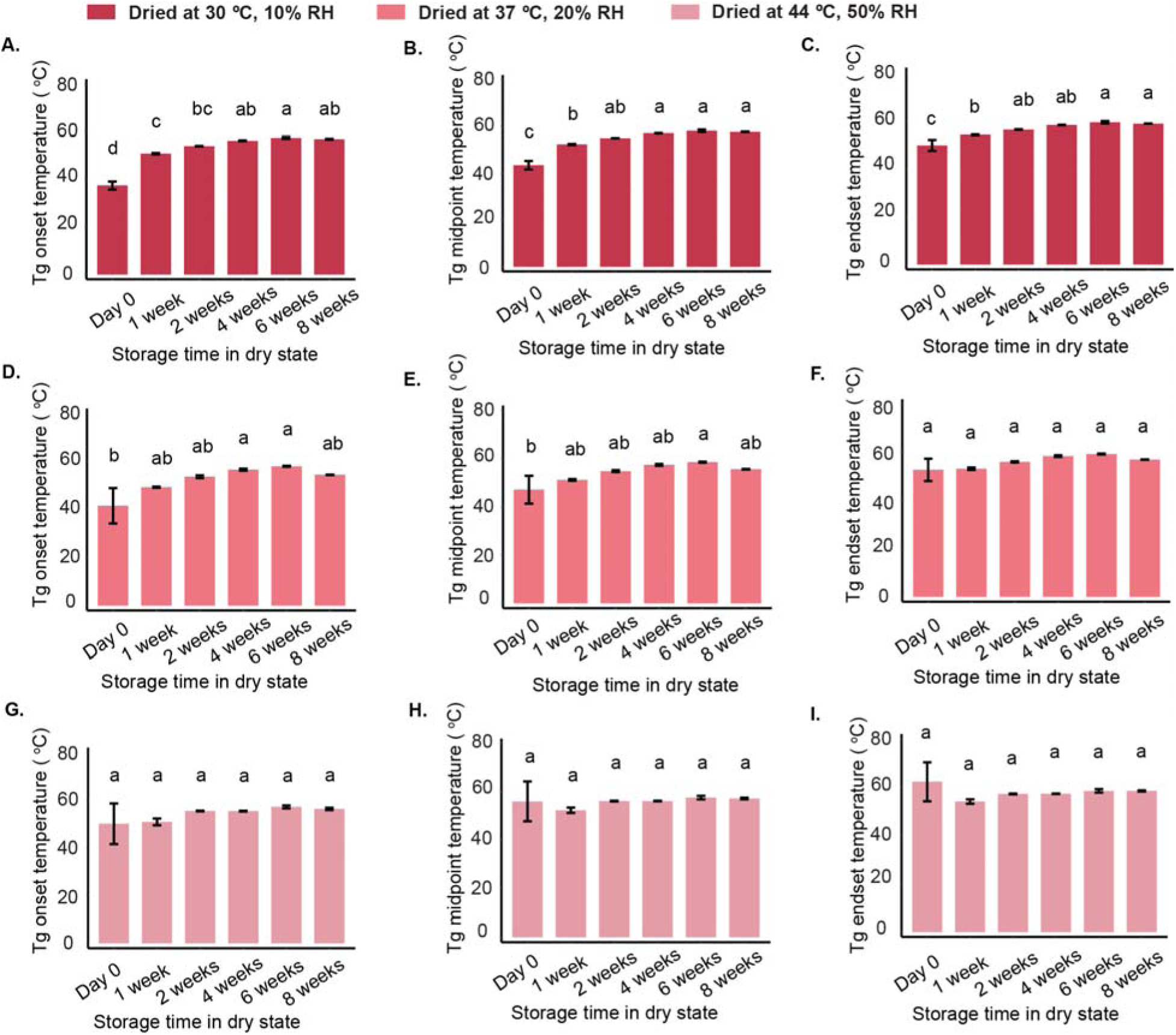
Glass transition temperature (Tg) of dried samples during storage under different drying conditions. Tg onset (A,D,G), midpoint (B,E,H), and endset (C,F,I) values of samples dried under three conditions are shown: 30 °C, 10% humidity (A-C); 37 °C, 20% humidity (D-F); and 44 °C, 50% humidity (G-I). Data are presented as mean ± SE. In each set of panels, different letters indicate statistical significance determined by one-way ANOVA followed by Tukey’s test.

**Supplementary Figure 31.**
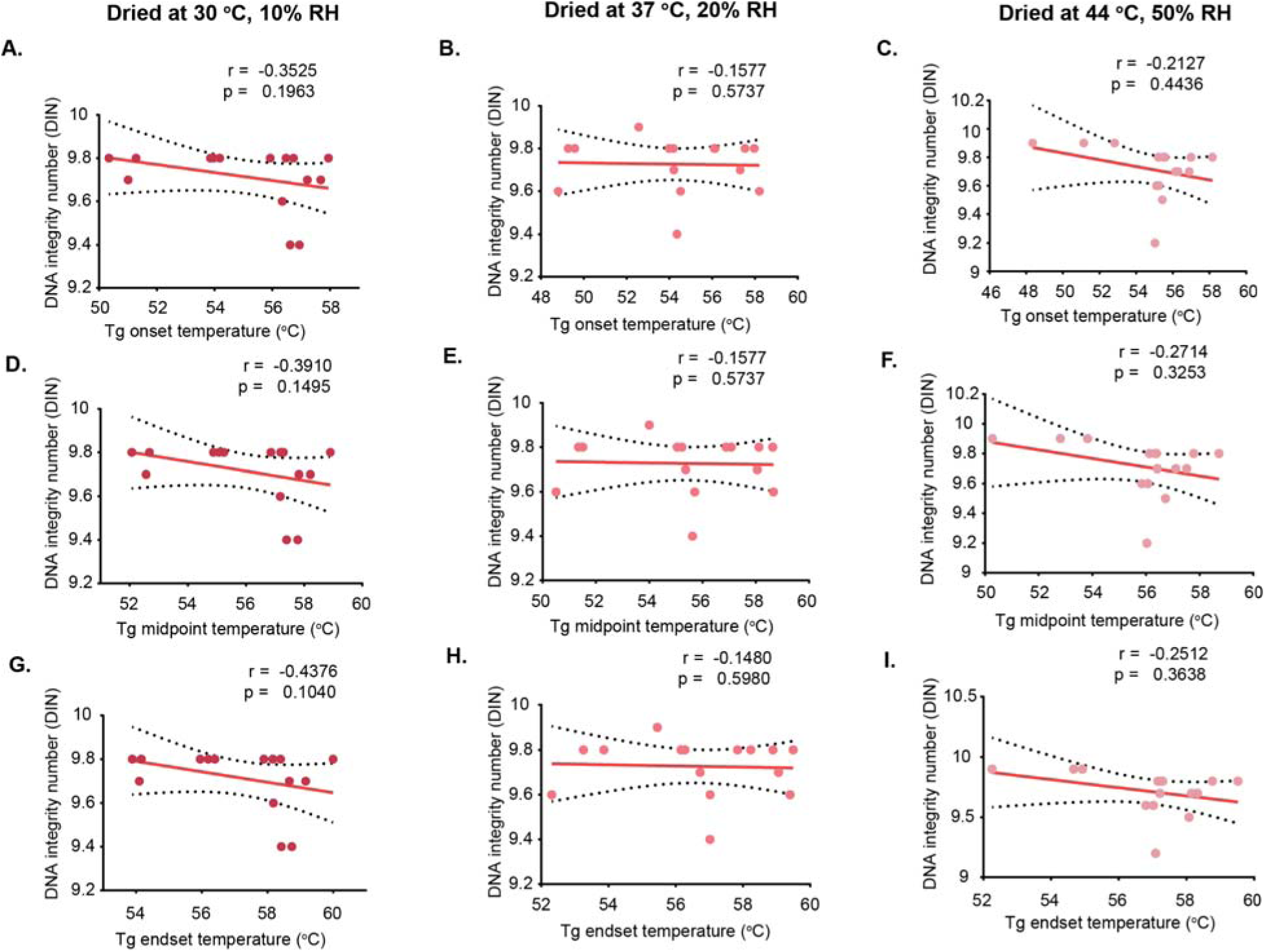
Correlation between DNA integrity and glass transition temperature (Tg) under different drying conditions during storage. Correlations between DNA integrity, assessed using the DNA Integrity Number (DIN), and Tg values are shown for three drying conditions: 30 °C, 10% humidity (A,D,G); 37 °C, 20% humidity (B,E,H); and 44 °C, 50% humidity (C,F,I). A-C show correlations with Tg onset, D-F show correlations with Tg midpoint, and G-I show correlations with Tg endset. Correlation coefficients (r) and significance values (p) were calculated using Pearson correlation for normally distributed data and Spearman correlation for non-normally distributed data. Each data point represents an individual replicate. Dashed lines indicate 95% confidence interval (CI).

**Supplementary Figure 32.**
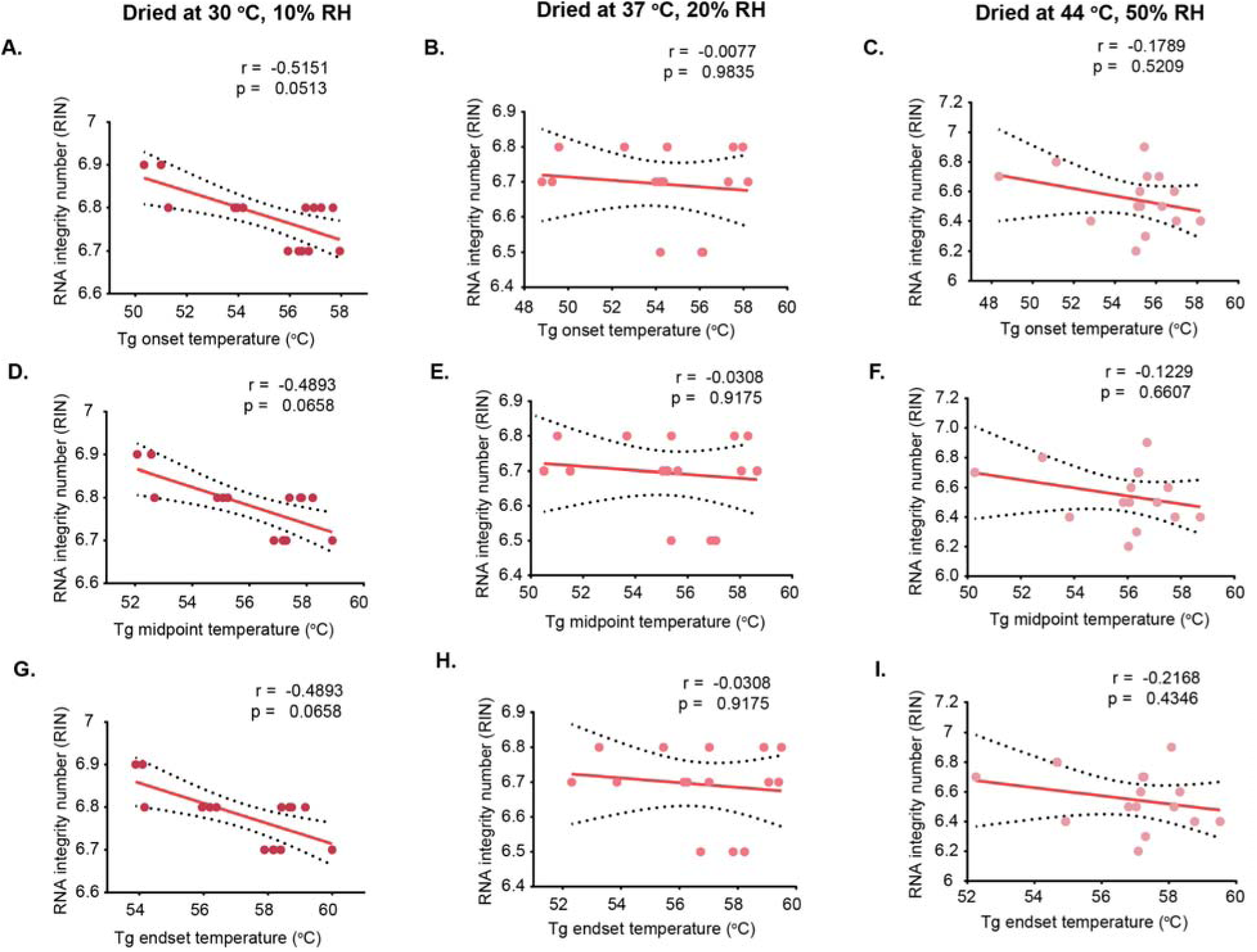
Correlation between RNA integrity and glass transition temperature (Tg) under different drying conditions during storage. Correlations between RNA integrity, assessed using the RNA Integrity Number (RIN), and Tg values are shown for three drying conditions: 30 °C, 10% humidity (A,D,G); 37 °C, 20% humidity (B,E,H); and 44 °C, 50% humidity (C,F,I). A-C show correlations with Tg onset, D-F show correlations with Tg midpoint, and G-I show correlations with Tg endset. Correlation coefficients (r) and significance values (p) were calculated using Pearson correlation for normally distributed data and Spearman correlation for non-normally distributed data. Each data point represents an individual replicate. Dashed lines indicate 95% confidence interval (CI).

**Supplementary Figure 33.**
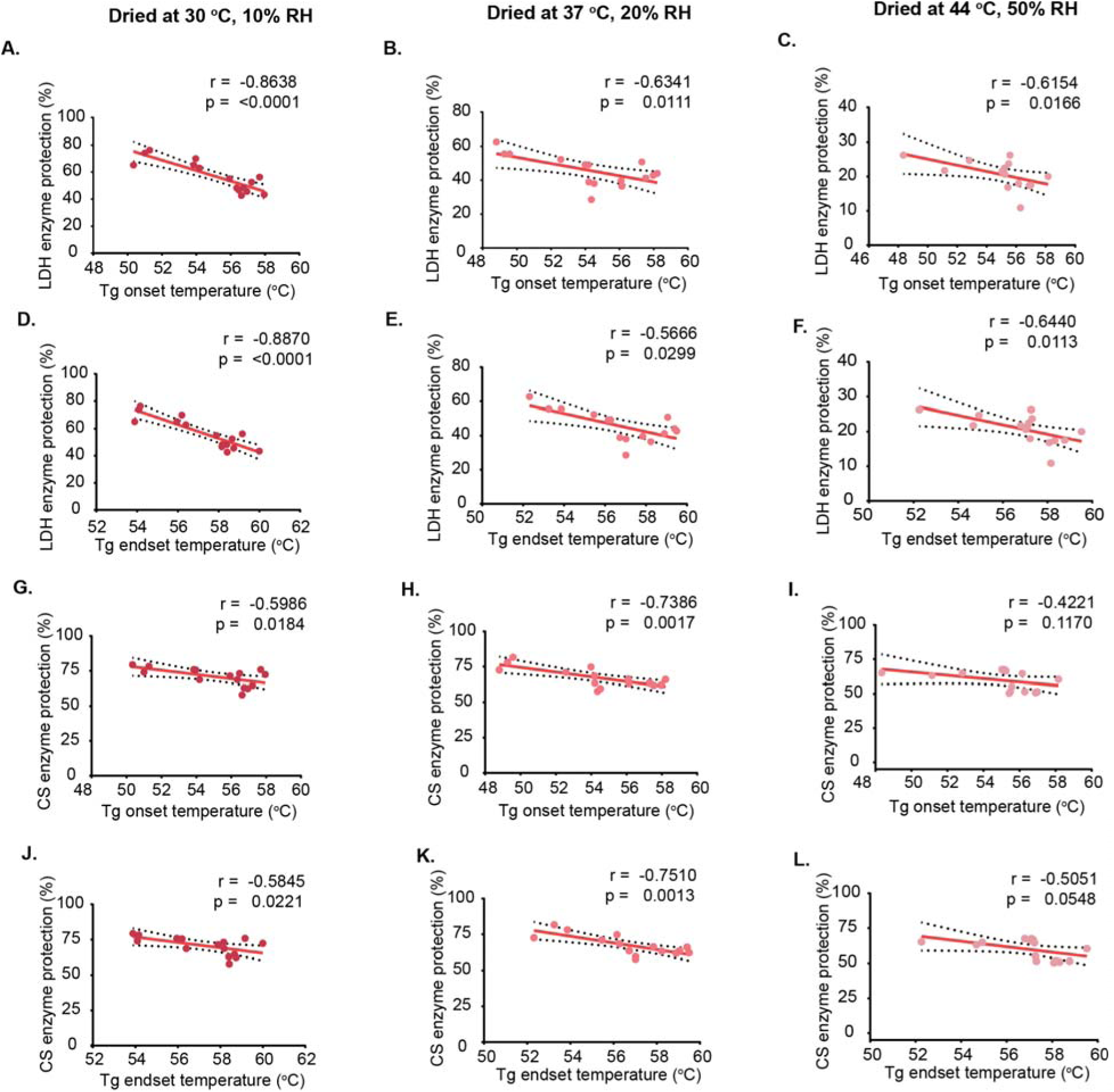
Correlation between glass transition onset and endset temperatures (Tg) and enzyme activity (LDH and CS) under different drying conditions over storage time. Samples were dried under three conditions: 30 °C, 10% humidity (A, D, G, J); 37 °C, 20% humidity (B, E, H, K); and 44 °C, 50% humidity (C, F, I, L). Correlations between Tg and enzyme activity are shown for lactate dehydrogenase (LDH; A-F) and citrate synthase (CS; G-L). Pearson correlation was used for normally distributed data, and Spearman correlation was used for non-normally distributed data. Each data point represents an individual replicate. Dashed lines indicate 95% confidence interval (CI).

**Supplementary Figure 34.**
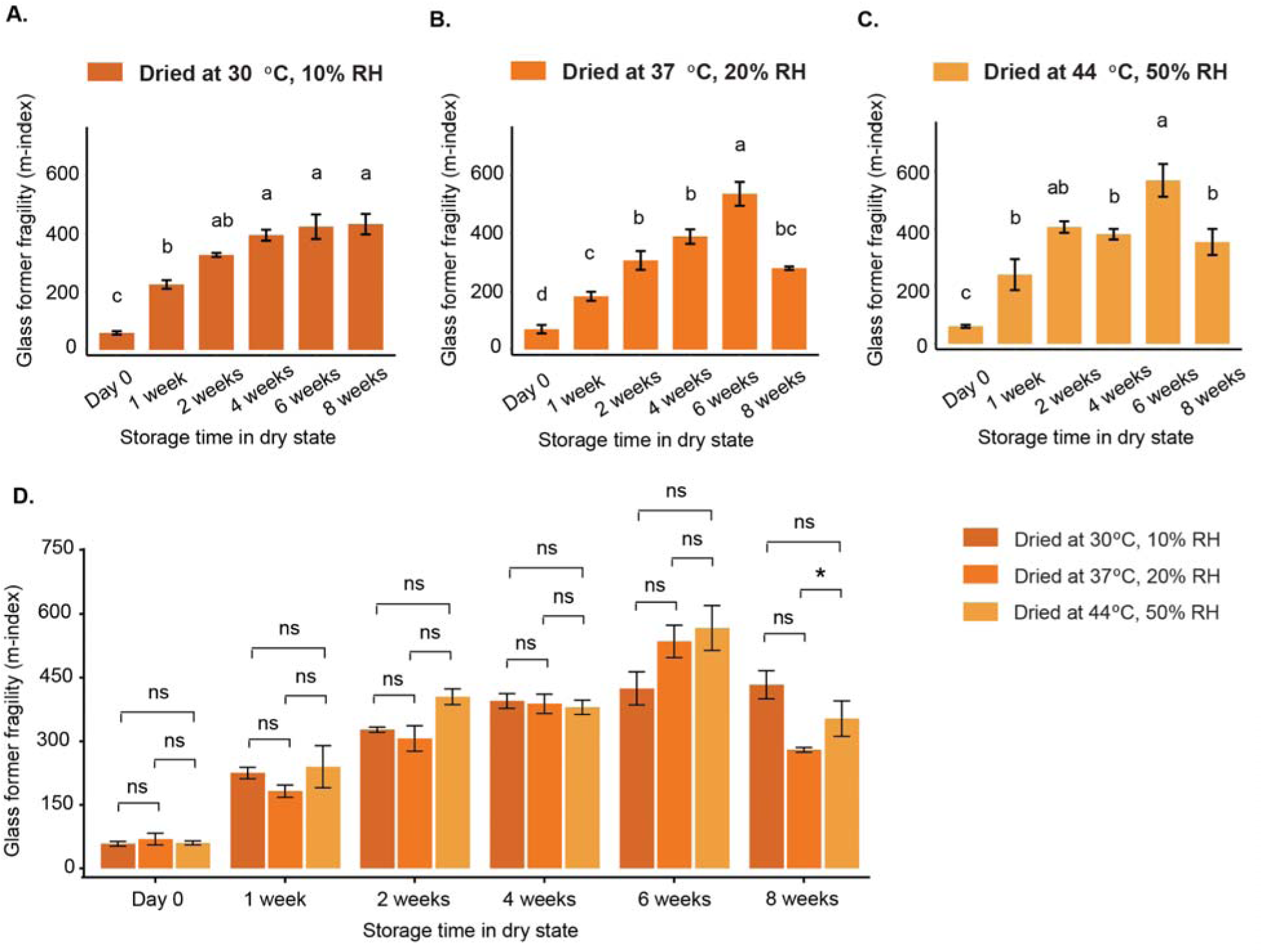
Glass former fragility (m-index) of dried samples during storage under different drying conditions. Fragility values of samples dried under three conditions (A-C): 30 °C, 10% humidity (A); 37 °C, 20% humidity (B); and 44 °C, 50% humidity (C). Direct comparison of fragility across the three drying conditions is shown in (D). Data are presented as mean ± SE. In panels A-C, different letters indicate statistical significance determined by one-way ANOVA followed by Tukey’s test. In panel D, statistical analysis was performed using two-way ANOVA with multiple comparisons.

**Supplementary Figure 35.**
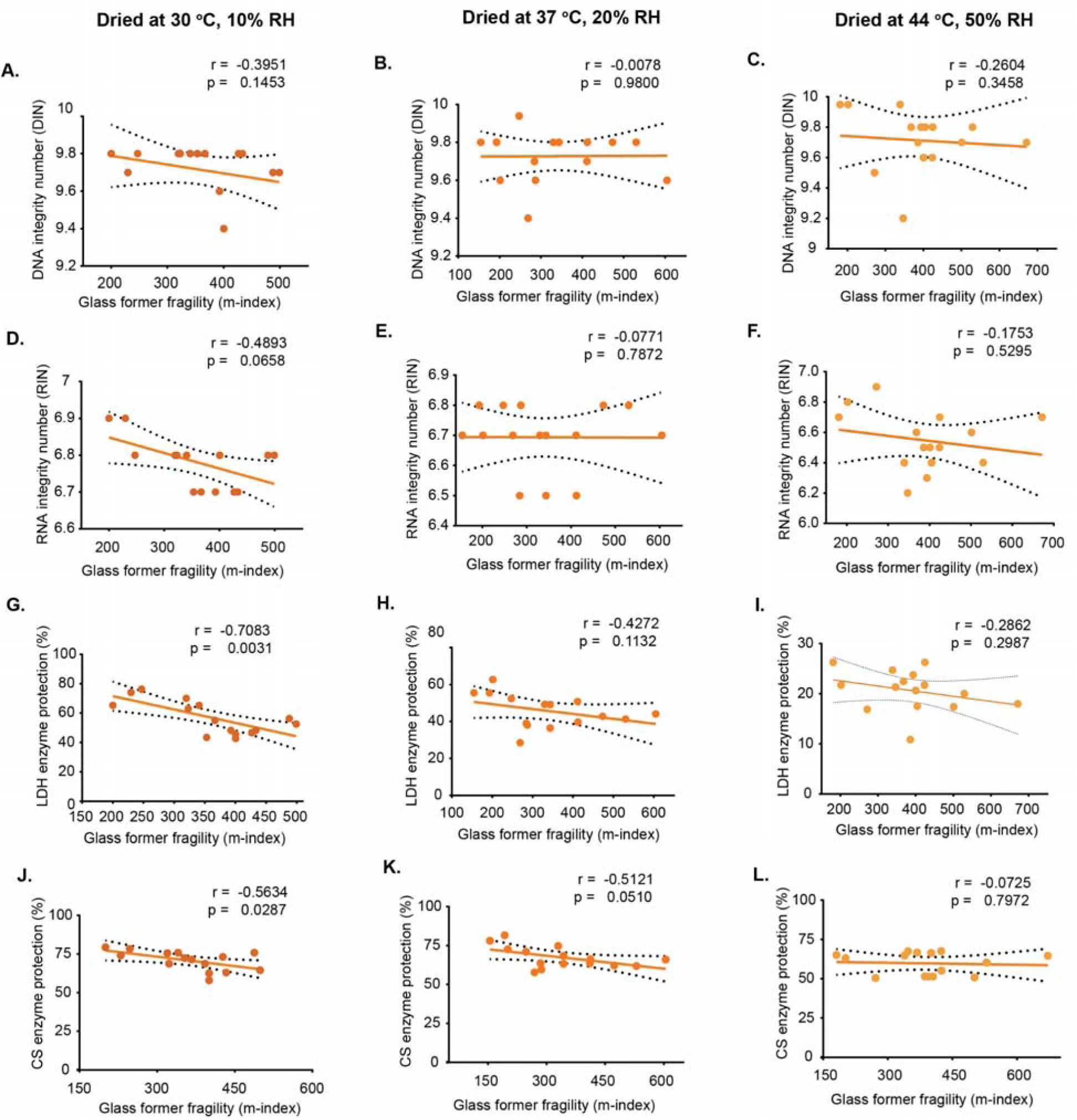
Correlation between glass former fragility (m-index) and nucleic acid integrity or protein activity under different drying conditions during storage. Samples were dried under three conditions: 30 °C, 10% humidity (A, D, G, J); 37 °C, 20% humidity (B, E, H, K); and 44 °C, 50% humidity (C, F, I, L). Correlations are shown for DNA integrity (A-C), RNA integrity (D-F), lactate dehydrogenase (LDH) activity (G-I), and citrate synthase (CS) activity (J-L). Pearson correlation was used for normally distributed data, and Spearman correlation was used for non-normally distributed data. Each data point represents an individual replicate. Dashed lines indicate 95% confidence interval (CI).

**Supplementary Figure S36.**
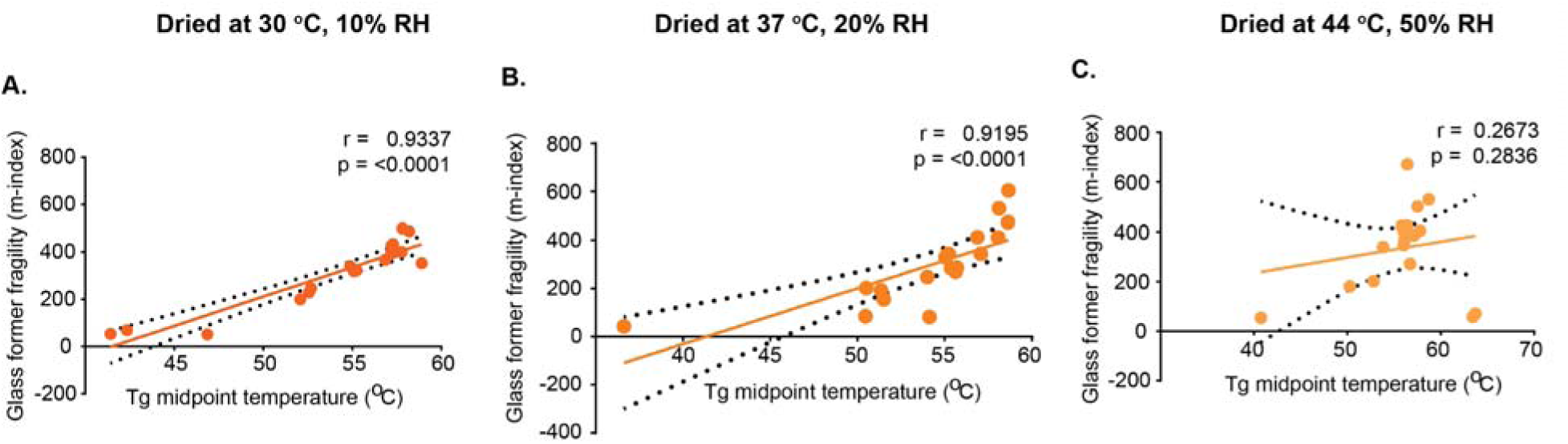
Correlation between glass transition (Tg) midpoint temperature and glass former fragility (m-index) under different drying conditions during storage. Samples were dried under three conditions: 30 °C, 10% humidity (A); 37 °C, 20% humidity (B); and 44 °C, 50% humidity (C). Correlation coefficients (r) and significance values (p) were calculated using Pearson correlation for normally distributed data and Spearman correlation for non-normally distributed data. Each data point represents an individual replicate. Dashed lines indicate 95% confidence interval (CI).

